# Convergent adaptation of true crabs (Decapoda: Brachyura) to a gradient of terrestrial environments

**DOI:** 10.1101/2022.12.09.519815

**Authors:** Joanna M. Wolfe, Lauren Ballou, Javier Luque, Victoria M. Watson-Zink, Shane T. Ahyong, Joëlle Barido-Sottani, Tin-Yam Chan, Ka Hou Chu, Keith A. Crandall, Savel R. Daniels, Darryl L. Felder, Harrison Mancke, Joel W. Martin, Peter K.L. Ng, Javier Ortega-Hernández, Emma Palacios Theil, N. Dean Pentcheff, Rafael Robles, Brent P. Thoma, Ling Ming Tsang, Regina Wetzer, Amanda M. Windsor, Heather D. Bracken-Grissom

## Abstract

For much of terrestrial biodiversity, the evolutionary pathways of adaptation from marine ancestors are poorly understood, and have usually been viewed as a binary trait. True crabs, the decapod crustacean infraorder Brachyura, comprise over 7,600 species representing a striking diversity of morphology and ecology, including repeated adaptation to non-marine habitats. Here, we reconstruct the evolutionary history of Brachyura using new and published sequences of 10 genes for 344 tips spanning 88 of 109 brachyuran families. Using 36 newly vetted fossil calibrations, we infer that brachyurans most likely diverged in the Triassic, with family-level splits in the late Cretaceous and early Paleogene. By contrast, the root age is underestimated with automated sampling of 328 fossil occurrences explicitly incorporated into the tree prior, suggesting such models are a poor fit under heterogeneous fossil preservation. We apply recently defined trait-by-environment associations to classify a gradient of transitions from marine to terrestrial lifestyles. We estimate that crabs left the marine environment at least seven and up to 17 times convergently, and returned to the sea from non-marine environments at least twice. Although the most highly terrestrial- and many freshwater-adapted crabs are concentrated in Thoracotremata, Bayesian threshold models of ancestral state reconstruction fail to identify shifts to higher terrestrial grades due to the degree of underlying change required. Lineages throughout our tree inhabit intertidal and marginal marine environments, corroborating the inference that the early stages of terrestrial adaptation have a lower threshold to evolve. Our framework and extensive new fossil and natural history datasets will enable future comparisons of non-marine adaptation at the morphological and molecular level. Crabs provide an important window into the early processes of adaptation to novel environments, and different degrees of evolutionary constraint that might help predict these pathways.

## Introduction

Over 80% of estimated species comprising extant multicellular life inhabit terrestrial and freshwater (“non-marine”) settings (Román-Palacios et al. 2022). Microbial life began to populate terrestrial habitats in the Precambrian, with eukaryotes potentially originating in non-marine settings around 1.6 Ga (Jamy et al. 2022), although major multicellular groups such as animals and plants were ancestrally marine. Their terrestrialization followed in the early Paleozoic (approximately 538–444 Ma), led by arthropods entering coastal and marginal marine settings (e.g., estuaries, lagoons), and plants that transformed the land and its sediments (Buatois et al. 2022). Although molecular divergence time estimates infer early Paleozoic ages for terrestrial arthropod crown groups (e.g., Bernot et al. 2023; Benavides et al. 2023), recognizable body fossils of millipedes, arachnids, and hexapods have recorded their presence on land by the onset of the Silurian–Devonian (443–359 Ma). Subsequently, these groups radiated to become prominent components of terrestrial biodiversity. Fossil evidence suggests potential transitions through marginal marine settings (Edgecombe et al. 2020; Lamsdell et al. 2020), but transitions for many modern groups lack such clues (e.g., the remipede sister group is now predominantly restricted to marine layers within anchialine caves), hinting at complex ecological pathways. Here, we examine the evolutionary history of a clade, the true crabs (Decapoda: Brachyura), that might provide insights into the early phases of adaptation from marine to non-marine environments, now obscured by extinction.

As with life in general, crabs have an unequivocally marine ancestor (Watson-Zink 2021). The largest group of Brachyura, called Eubrachyura, which contains all non-marine members, could be as old as the mid-Jurassic (183–161 Ma) based on phylogenomic divergence time estimates (Wolfe et al. 2019). During the “Cretaceous Crab Revolution” (145–66 Ma), many now-extinct lineages appeared briefly, accompanied by the divergence of many extant superfamilies (Wolfe et al. 2019; Luque et al. 2019b; Wolfe et al. 2021). Although the direct record of fossil crabs from non-marine sediments is depauperate, one well-preserved example of a completely extinct non-marine eubrachyuran lineage is known from around 100 Ma (Luque et al. 2021), and chelipeds of uncertain affinity from non-marine sediments around 74 Ma (Robin et al. 2019). Together, these fossils suggest that crabs have been entering non-marine habitats for the majority of their evolutionary history.

Complementary to direct fossil evidence, dated phylogenies and character mapping have recently been applied to investigate the evolution of crab terrestriality (Davis et al. 2022; Tsang et al. 2022). Eubrachyura has been previously divided into two presumed clades based on the position of the male gonopores: “Heterotremata” and Thoracotremata. In a 10-gene molecular study focused on the relationships of the clade Thoracotremata, the common ancestor of this clade was found to be “semi-terrestrial” (in Tsang et al. [2022], this referred to intertidal habitats) and Cretaceous in origin, with at least four transitions to terrestrial and two or three transitions to freshwater lifestyles, all within the Cenozoic (Tsang et al. 2022). In one instance, the authors estimated at least six returns to subtidal marine habitats, and hypothesized that sesarmid crabs (specifically *Geosesarma*, vampire crabs) transitioned from terrestrial to freshwater habitats (Tsang et al. 2022). A separate supertree-based study across Decapoda inferred three transitions to terrestriality and three to freshwater, and one reversal from terrestrial to marine habitats, within all of Brachyura (Davis et al. 2022). The oldest event, encompassing the freshwater heterotreme groups Potamoidea, Gecarcinucoidea, and Pseudothelphusoidea, occurred in the upper Cretaceous, with others in the Cenozoic. Additionally, Davis et al. (2022) inferred higher rates of speciation in non-marine crabs, but habitat shifts were not found to be a significant causal factor driving crab diversity.

The aforementioned phylogenetic studies, however, treated marine, terrestrial, and freshwater lifestyles as largely discrete ecologies for crabs. Indeed, previous studies have described a gradient of terrestrial change based on independence from standing water (e.g., Bliss 1968; Powers and Bliss 1983; Hartnoll 1988), with the caveat that no known crab is completely independent from water throughout its entire life cycle. Others (e.g., Yeo et al. 2008; Cumberlidge and Ng 2009; Cumberlidge et al. 2009) focused on the seven exclusively (“primary”) freshwater crab families and their vicariant biogeography leading to high endemicity and risks of extinction, but rarely drew comparisons with terrestrial crabs. Recently, Watson-Zink (2021) unified the conceptualization of the terrestrial and freshwater crab lifestyles as a series of ecological, morphological, and physiological traits describing grades of terrestriality (described in **Table S1**). Crabs can leave fully marine lifestyles (**Fig. 1a-e**) along either of two transition pathways: through marine-associated environments (e.g., the “direct” pathway of Tsang et al. [2022], via intertidal, mangroves, beaches: **Fig. 1f-j**) or through freshwater environments (e.g., the “indirect” pathway via estuaries, rivers: **Fig. 1k-o**, akin to the transition in amphibians). Each grade of terrestriality is loosely associated with habitats: lower intertidal and estuaries (grade 1), upper intertidal and freshwater (grade 2), beaches and riverbanks (grade 3), and coastal forests and jungles, including tree climbing (grades 4-5; Watson-Zink 2021). Less terrestrial crabs (grades 1 and 2) in either pathway can tolerate fluctuating environments, with osmoregulatory ability likely playing a major role in these lifestyles (Watson-Zink 2021). Crabs of higher terrestriality (grades 3-5) possess further morphological and developmental adaptations, such as branchiostegal lungs and water-wicking setae to prevent desiccation, and increasingly abbreviated larval development and parental care (Watson-Zink 2021). Note that the grades do not represent an ultimate “goal” of terrestriality, as many groups successfully remain and diversify within lower grades. Indeed, it is evident that multiple brachyuran families have repeatedly evolved members of both transition pathways and various grades, but their distribution across crab phylogeny (within and beyond Thoracotremata) over time remains unclear.

**Figure 1.**
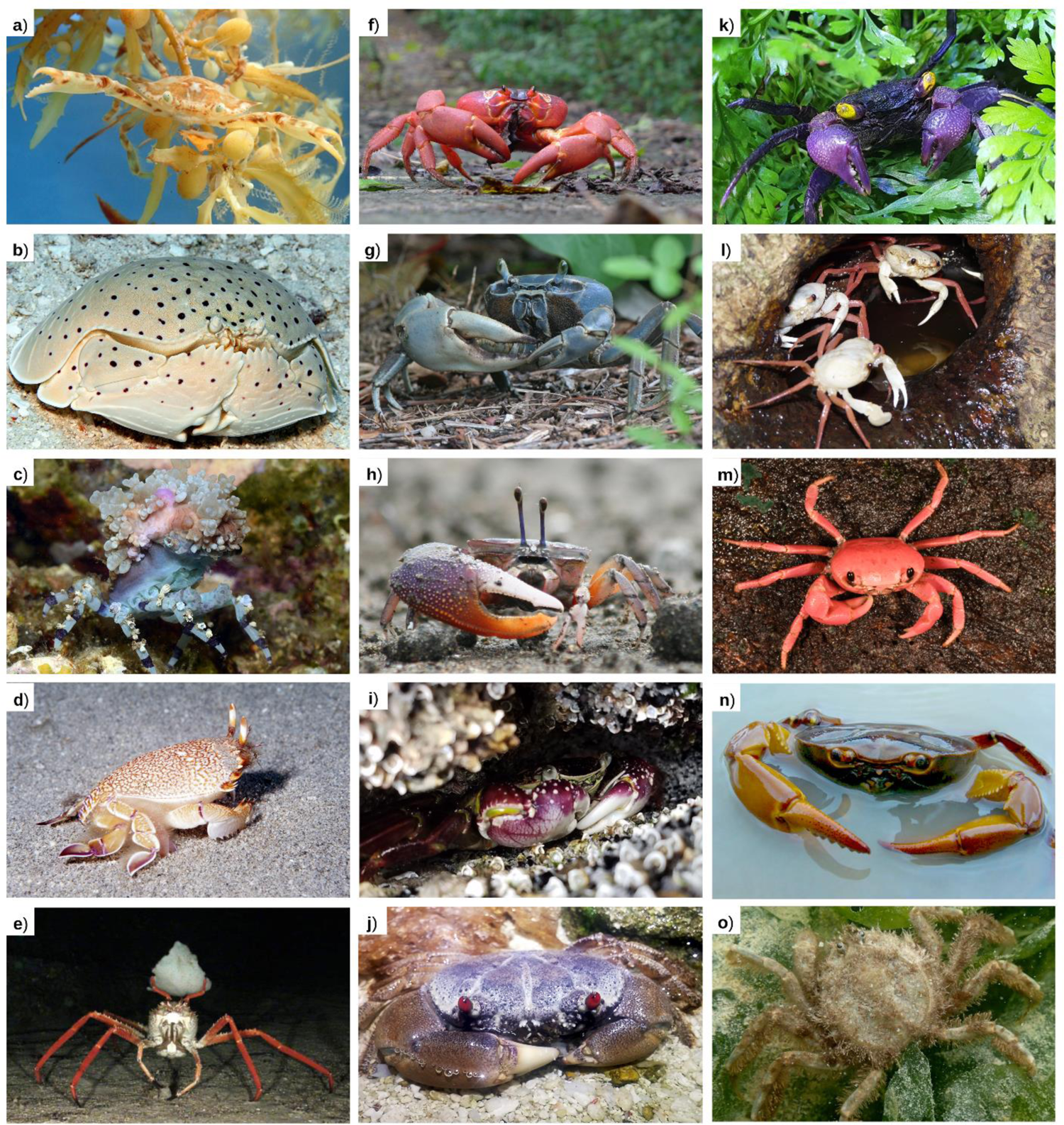
Representative brachyurans displaying different lifestyles and grades of terrestriality. (**a-e**) Fully marine lifestyle, grade 0; (**f-j**) direct marine transition pathway, grades 1-5 bottom to top; (**k-o**) indirect freshwater transition pathway, grades 1-5 bottom to top. (**a**) Portunidae: *Portunus sayi* (Bermuda); (**b**) Calappidae: *Calappa calappa* (Kwajalein Atoll, Marshall Islands); (**c**) Epialtidae: *Cyclocoeloma tuberculatum* (Anilao, Philippines); (**d**) Raninidae: *Ranina ranina* (Oahu, Hawaii, USA); (**e**) Homolidae: *Paromola cuvieri* (Gorringe Ridge, Portugal); (**f**) Gecarcinidae: *Gecarcoidea natalis* (Christmas Island, Australia); (**g**) Gecarcinidae: *Cardisoma guanhumi* (Fort Lauderdale, Florida, USA); (**h**) Ocypodidae: *Uca heteropleura* (Pacific coast, Panama); (**i**) Grapsidae: *Leptograpsus variegatus* (Tasmania, Australia); (**j**) Eriphiidae: *Eriphia sebana* (Heron Island, Queensland, Australia); (**k**) Sesarmidae: *Geosesarma dennerle* (aquarium specimen); (**l**) Deckeniidae: *Madagapotamon humberti* (Montagne de Français Reserve, Madagascar); (**m**) Gecarcinucidae: *Ghatiana botti* (Sindhudurg, India); (**n**) Pseudothelphusidae indet. (Santander, Colombia); (**o**) Hymenosomatidae: *Hymenosoma orbiculare* (Langebaan Lagoon, South Africa). Photo credits: (a) Jessica Riederer; (b,c) Jeanette and Scott Johnson; (d) John Hoover; (e) © OCEANA; (f) John Tann, license CC-BY; (g) Tom Friedel, license CC-BY 3.0; (h) Kecia Kerr and Javier Luque; (i) Joanna Wolfe; (j,n) Javier Luque; (k) Henry Wong; (l) Sara Ruane; (m) Tejas Thackeray; (o) Charles Griffiths.

To resolve the convergent evolution and timing of terrestriality, we present the most robust molecular taxon sampling to date for Brachyura, representing 333 species and 88 of 109 families. As our data represent only Sanger sequences of 10 loci, revision of brachyuran systematics is beyond the scope of the current study (Timm and Bracken-Grissom 2015), and efforts are currently underway to clarify deep relationships using phylogenomics. Furthermore, to partially ameliorate false confidence in the topology, we provide additional metrics describing the degree of nodal uncertainty. Using these data, we contrast divergence times inferred using 36 newly vetted detailed calibrations (Luque et al. 2023) with traditional node dating, and 328 calibrations sampled from the Paleobiology Database under the fossilized birth-death (FBD) and skyline models of tree evolution, one of only a few empirical comparisons of its type. Finally, we summarize the natural history traits of each sampled crab to assign each along a gradient of transitions from marine to terrestrial lifestyles (Watson-Zink 2021), and use Bayesian threshold models of ancestral state reconstruction to estimate convergent events.

## Materials and Methods

### Taxon and gene sampling, DNA extraction, PCR, and sequencing

The molecular dataset includes 88 families, 263 genera, 333 species and 338 individuals within the infraorder Brachyura (Ng et al. 2008; Poore and Ahyong 2023; WoRMS 2022), as well as six outgroups for a total of 344 tips. 41% of sequence data were new, with the remainder obtained from GenBank (**Table S2**). A total of 10 genes were selected based on previous phylogenetic research on decapods (Spears and Abele 1998; Schubart et al. 2000; Tsang et al. 2008; Bracken-Grissom et al. 2013; Tsang et al. 2014). These included two mitochondrial ribosomal RNA (rRNA) coding genes, two nuclear rRNA genes, and six nuclear protein-coding genes (**Text S1**). A minimum of two genes were required for each taxon included in the analysis, with an average of seven genes available per tip.

Genomic DNA was extracted from the gills, abdomen, pereopod, or pleopod, using the Qiagen DNeasy® Blood and Tissue Kit, QIAamp DNA Mini Kit, or QIAamp DNA Micro Kit. Gene regions were amplified with polymerase chain reaction (PCR) using one or more sets of primers (**Text S1**). PCR amplifications and sequencing reactions were performed as described in **Text S1**.

### Phylogenetic analysis

Sequences (n = 2249) were assembled and trimmed within Geneious Prime (Kearse et al. 2012). Protein-coding genes were checked for pseudogenes following Song et al. (2008); these were individually aligned in Geneious v.2021.0.1 using MAFFT (Katoh and Standley 2013), as were rRNA genes. To remove regions of questionable homology, rRNA alignments were masked in GBlocks v.0.91b (Castresana 2000) under “less stringent” parameters. Alignments were concatenated in Geneious Prime.

Best-fitting partitions (Chernomor et al. 2016) and substitution models (Kalyaanamoorthy et al. 2017) were selected in IQ-TREE v.2.1.2 (Minh et al. 2020b). The best-fitting scheme was used to estimate the concatenated maximum likelihood (ML) phylogeny, also in IQ-TREE. Ultrafast (UF) bootstrap values were calculated from 1000 replicates (Hoang et al. 2018). We conducted a Bayesian inference (BI) analysis of the concatenated loci using MrBayes v.3.2.7 (Ronquist et al. 2012). Two runs and four chains were run for 35 million generations with 25% burnin. Convergence was assessed by reaching effective sample size >200 for every parameter, and by evaluating posterior distributions in Tracer v.1.7.1 (Rambaut et al. 2018).

### Fossil calibration and divergence time inference

We compared two strategies for fossil calibration: (1) 36 newly vetted node calibrations, and (2) 328 fossil occurrences from the Paleobiology Database (PBDB; http://paleobiodb.org/). For node calibration, all calibrations followed best practices regarding specimen data, morphological diagnosis, and stratigraphy (Parham et al. 2012; Wolfe et al. 2016; extensive details in Luque et al. 2023, summary in **Table S3** herein), and were assigned to a crown group node at the family level or higher. This node dating strategy used a birth-death tree prior and uniform calibration age distributions.

We downloaded fossil occurrences from the PBDB on March 23, 2022, for Brachyura at family-level taxonomic resolution (details in **Text S1**). We randomly subsampled 10% of the 3,276 remaining occurrences, resulting in a computationally tractable 328 occurrences. The subsample was a reasonable sample of ages (including the oldest possible age), represented 69% of fossil families, and slightly overrepresented non-marine paleoenvironments (details in **Table S4**). All fossil occurrences were assigned age ranges from the PBDB, each with a uniform distribution (Barido-Sottani et al. 2019, 2020). To incorporate these fossil samples as part of the inferred evolutionary process, we used the unresolved time-homogeneous fossilized birth-death (FBD) tree prior (Stadler 2010; Heath et al. 2014; Bapst et al. 2016; O’Reilly and Donoghue 2020), with parameters described in **Text S1**.

To reflect the complete absence of brachyuran fossils from earlier than the Jurassic, which represents a known ghost lineage when compared to the diversity and abundance of outgroup anomuran fossils, thus earlier diversification of the sister group (Hegna et al. 2020; Wolfe et al. 2021), we also analyzed the fossil occurrence calibration set using a birth-death skyline tree prior with sequential sampling (BDSS; Stadler et al. 2013; Culshaw et al. 2019). Fossil sampling proportion was modeled as time-heterogeneous with time slices before and after the oldest fossil sample (details in **Text S1**) using the TreeSlicer function from the *skylinetools* package (https://github.com/laduplessis/skylinetools).

All divergence time analyses were conducted in BEAST2 v.2.6.7 (Bouckaert et al. 2019) using a fixed starting tree derived from our ML concatenated results (detailed parameters in **Text S1**). Fossil occurrences (in FBD and BDSS analyses only) were added to the starting tree as “rogues” (able to move within pre-assigned family level constraints following Barido-Sottani et al. [2022], as most decapod fossils are fragmentary and cannot be confidently assigned). For each calibration strategy and tree prior, we compared two clock models: relaxed lognormal (Drummond et al. 2006) and random local (Drummond and Suchard 2010). Analyses used four to six runs for at least 450 million generations with 25% burnin. Convergence was assessed as above. We visualized the results from different parameter sets using *chronospace* scripts (Mongiardino Koch et al. 2022), which provide a multidimensional representation of inferred node ages that can be broken down by different factors and models.

### Ancestral state reconstruction

To code character states representing a gradient of terrestriality, we used a modified version of the trait-by-environment associations defined by Watson-Zink (2021), additional details in **Table S1** and **Text S1**. Distinct transition pathways were defined for marine and freshwater routes. For both, we added a grade 0, indicating that the ancestral state for all crabs is fully marine (**Fig. 1a-e**). Using this framework, we coded discrete grades of terrestriality following two schemes. The first scheme coded the taxa that were sequenced in our molecular phylogeny and required justification from natural history literature on: adult habitat, osmoregulatory status, larval developmental strategy, primary respiratory structure, water-wicking setae, burrow type, and diurnal activity period (**Table S5**, **Text S2**). As our phylogeny sampled 4% of brachyuran species as tips, we also constructed a scheme to estimate grades for unsampled species for which the phylogenetic positions are unknown. For this scheme, we downloaded all taxonomic data, including non-marine taxa, from WoRMS as of June 8, 2021. For families sampled in our tree, we used WoRMS to estimate the number of species that fall into each grade and accordingly assigned prior distributions to each tip on the molecular phylogeny (**Table S6**).

First, we used stochastic character mapping (Bollback 2006; Revell 2012) to infer ancestral states at each node, using a simplified single dataset (details in **Text S1**). Next, we used Bayesian threshold models (Felsenstein 2012; Revell 2014) to account for gradients of change. A character coded with discrete ordered states (i.e., our grades) was assumed to evolve according to an unobserved continuous trait called “liability” (here representing the coded natural history traits combined with additional unobserved factors). Following Sallan et al. (2018), we assume thresholds represent the amount of change in terrestriality traits that allows a habitat shift. As there are two independent transition pathways, these were analyzed separately for each pathway (from marine to non-marine). Threshold models for ancestral state reconstruction were implemented as the *ancThresh* function in *phytools* (Revell 2012). Each *ancThresh* model was run for 150 million generations with 20% burnin (Revell 2014; Sallan et al. 2018).

## Results

### Phylogenetic relationships

The concatenated alignment length comprised 7,516 bp in total from two mitochondrial rRNA, two nuclear rRNA, and six nuclear protein-coding genes (gene trees visualized in **Figs. S1-10**). Results using ML and BI were similar, with some deeper nodes (higher than family level) maintaining low to moderate support (UF bootstrap = 50-94; **Figs. 2, S11**) with ML and generally stronger support (most posterior probabilities ≥ 0.98; **Fig. S12**) with BI. For each node of the ML tree, the gene concordance factor (gCF) reflects the percentage of loci containing all the descendant taxa, and site concordance factor (sCF) the percentage of sites supporting the node (Minh et al. 2020a). Both concordance factors illustrate a spectrum of support across nodes that are fully supported with UF bootstraps (**Fig. S13**). The average gCF was 32.73 (**Figs. S11, S13**), indicating one third of loci support the average node. However, nearly half of sites support the average node (average sCF = 45.64), demonstrating the benefit of concatenation for small numbers of loci.

**Figure 2.**
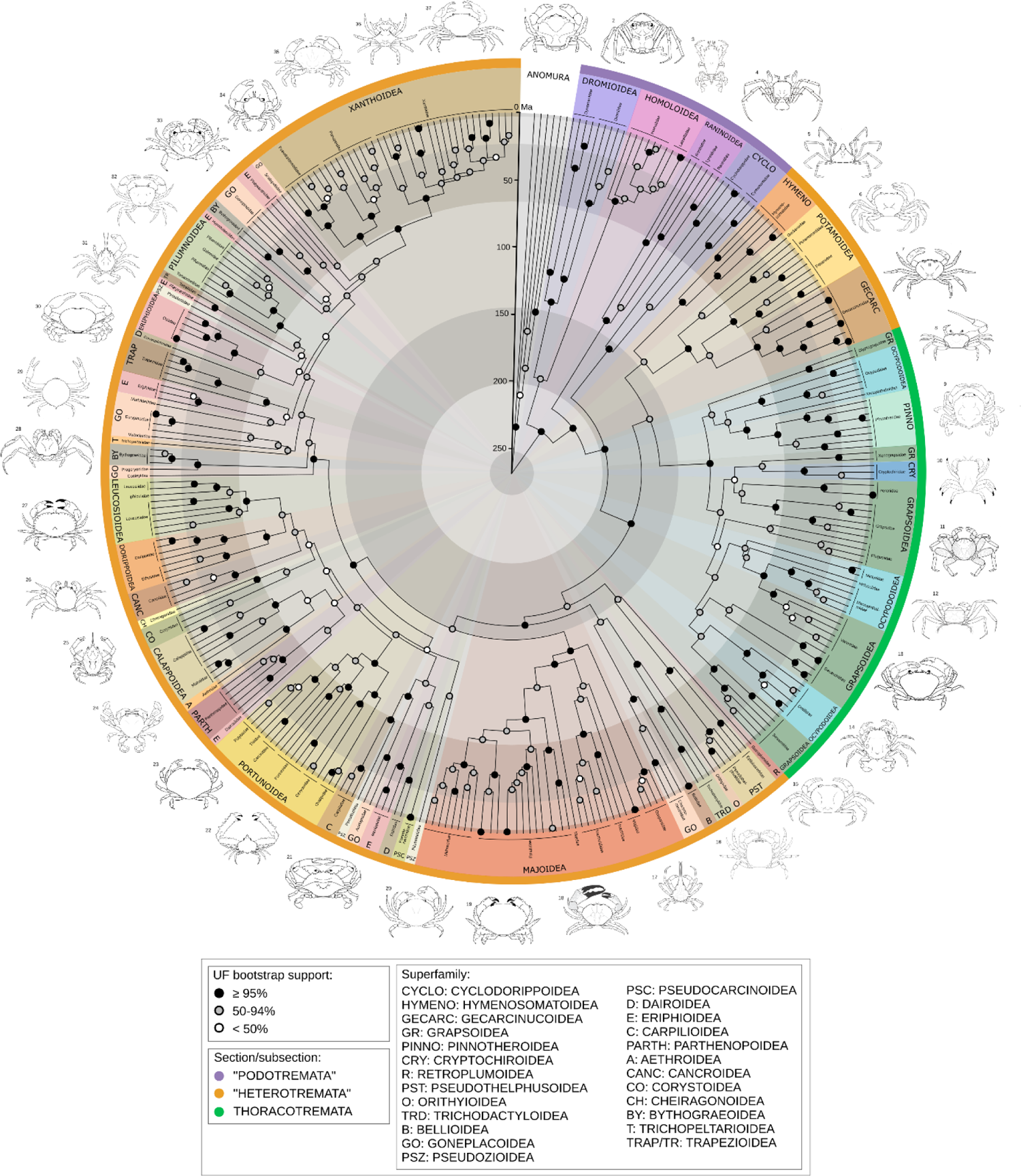
Summary of phylogeny and divergence time estimates for Brachyura (88 brachyuran families, 263 genera, 333 species, 338 individuals plus six outgroups). Posterior ages were estimated in BEAST2 using a fixed topology resulting from the concatenated ML analysis in IQ-TREE, 36 vetted node calibrations, a birth-death tree prior, and relaxed lognormal clock model. Shaded circles at nodes represent ultrafast bootstraps. Pie slices are colored by superfamily, with the outermost ring colored by taxonomic section. Line drawings, one representative per superfamily (numbers corresponding to taxa in Table S7), by Javier Luque and Harrison Mancke.

Revision of brachyuran systematics at nodes above the family level would best be undertaken with phylogenomic scale data (Wolfe et al. 2019 and phylogenetic informativeness profiles, **Fig. S14**), therefore we only briefly summarize the topology results here and in **Figure 2**. Podotremes are paraphyletic with respect to Eubrachyura, forming the following successive clades: Dromioidea + Homoloidea (this pairing has low support from ML, but strong from BI), Raninoidea, and Cyclodorippoidea (latter two clades with full support). Within Eubrachyura, subsection Heterotremata is paraphyletic with respect to monophyletic Thoracotremata. The so-called primary freshwater crabs are polyphyletic. The African and Eurasian groups (Potamoidea and Gecarcinucoidea) form a clade (UF bootstrap = 95, posterior probability = 0.89), and are themselves the sister clade of the Gondwanan Hymenosomatoidea (UF bootstrap = 92, posterior probability = 1). Together, this group comprises the sister group of Thoracotremata, with moderate support from ML (albeit with low concordance factors) and full support from BI. Within Thoracotremata, some higher-level relationships are weakly supported by ML (UF bootstraps < 75%, low gCF), but most nodes are similar to BI (where they have moderate to high support). Both Grapsoidea and Ocypodoidea are polyphyletic.

Meanwhile, the American freshwater groups branch off within clades including the deepest (Pseudothelphusoidea) and second deepest (Trichodactyloidea) divergences within the main heterotreme group, although these nodes are not strongly supported by traditional metrics in either analysis (concordance factors for both are > 60, some of the highest in our data). The remaining heterotremes are subdivided into Majoidea, and two large supported clades containing 24 and 23 families, respectively. Within these latter clades, the superfamilies Eriphioidea and Goneplacoidea are strongly polyphyletic. Some deep splits within both clades are poorly supported (some nodes UF bootstrap < 50, concordance factors = 0, posterior probability < 0.8).

We find that 76 (about 70%) of all sequenced families are monophyletic (or are represented by a single terminal), with the same exceptions in both ML and BI trees. Paraphyletic families are: the podotremes Homolidae (containing Latreillidae), Raninidae (containing Lyreididae), and Cyclodorippidae (containing Cymonomidae), and the heterotremes Epialtidae (containing Mithracidae), Carcinidae (containing Thiidae and Polybiidae), Corystidae (containing Cheiragonidae), Leucosiidae (containing Iphiculidae), and Pilumnidae (containing Galenidae). Polyphyletic families, all within heterotremes, are: Majidae, Bythograeidae, Platyxanthidae, and Pseudoziidae.

### Divergence times

Results of divergence time inference vary depending on parameters used, with results of the vetted calibration strategy distinct on the major axis (**Fig. 3a**), and the FBD and BDSS results being similar to one another. Although none of the random local clock analyses converged after extensive runtime, we plotted samples from their individual chains with burnin of 50% to reduce the effect of poor mixing; when included, the choice of clock model also differs significantly (**Fig. 3b**), but mostly in the same direction at the same nodes as the calibration strategy (**Fig. 3c**). In case the unconverged analyses were skewing the results, we plotted results from relaxed lognormal clocks alone (**Fig. S15**), finding the groupings by calibration strategy were mostly upheld, although the spread along axis 2 decreased for vetted calibrations especially.

**Figure 3.**
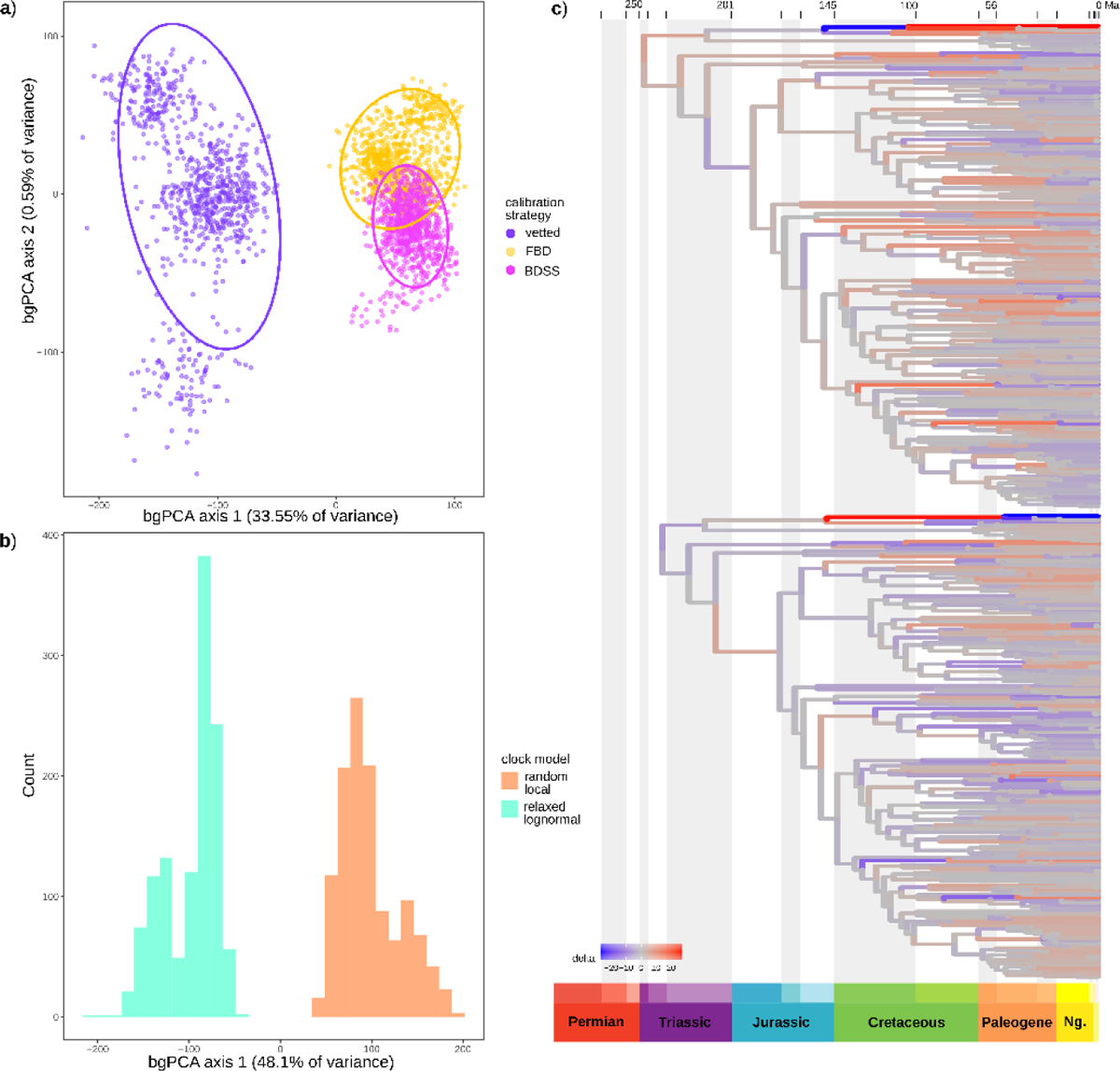
Sensitivity of divergence time estimates to inference strategy plotted with *chronospace*, with outgroup taxa removed from these analyses. (**a**) Between-group principal component analysis (bgPCA) separating chronograms by calibration strategy and (**b**) by clock model (note: analyses with the random local clock model did not converge, so individual chains are sampled here with 50% burnin). (**c**) Theoretical topologies representing change in branch lengths from the mean along the major bgPCA axis, discriminating based on calibration strategy. Negative extreme (−1 SD branch length) above, positive extreme (+1 SD) below.

Similar to Wolfe et al. (2019), using the vetted node calibrations, the divergence of Meiura (i.e., the root node) is inferred in the Permian (mean age at 268 Ma), while crown group Brachyura diverged in the Triassic (mean age 231 Ma), and crown group Eubrachyura in the Jurassic (mean age 172 Ma). Superfamily level divergences were inferred in the Jurassic for podotremes, and almost entirely within the Cretaceous within Eubrachyura (**Fig. S16**).

Divergence estimates using fossil occurrence sampling were both considerably younger than with the vetted calibrations, with the root estimate in the Jurassic in both cases (mean ages at 180 Ma for FBD, and 191 Ma for BDSS; **Figs. S17-18**). The PBDB calibrated analyses are relatively immune to the root prior (**Fig. S19a**). In the BDSS analysis, the inferred parameters were: sampling proportion 0.062 (95% HPD of 2.72e-3 to 0.23), death rate 0.091 (95% HPD of 0.024 to 0.23), and birth rate shown in **Figure S20**. Most other nodes were similarly compressed, with crown group Brachyura in the Jurassic and superfamily level divergences pushed to the Upper Cretaceous and Paleogene, although the posterior did not follow the marginal prior at some nodes (**Fig. S19b**). The PBDB analyses often inferred the placement of “rogue” fossils within the stem groups of their families. Consequently, a number of family level crown group ages were underestimated relative to their known vetted calibrations (e.g., Dromiidae, Dynomenidae, Lyreididae, Percnidae, Varunidae, Euryplacidae, and Panopeidae; **Table S3**). These families are among the most sensitive nodes, in addition to several nodes within Majoidea (**Fig. S19b**).

### Evolution of terrestriality

In the summary of stochastic character mapping, six total shifts from marine to non-marine were inferred (**Fig. S21**). These were at the base of Pseudothelphusoidea, Trichodactylidae, Menippidae, Eriphiidae, and Oziidae, and at the base of the clade of (Thoracotremata, Hymenosomatidae, Potamoidea, Gecarcinucidae). The latter node was split, with slightly higher posterior probability of being freshwater than terrestrial or marine, leading to a freshwater node for the base of (Hymenosomatidae, Potamoidea, Gecarcinucidae) and a terrestrial common ancestor of Thoracotremata. As individual stochastic character maps inferred shifts on branches leading to a single coded tip (e.g., *Carcinus maenas*), the median number of shifts to non-marine was 13. One shift from the freshwater to terrestrial pathway was found in Hymenosomatidae (note, all non-zero grades were collapsed, so the “terrestrial” members of this family are intertidal). Two shifts from terrestrial to freshwater were found in Glyptograpsidae and Varunidae, respectively. Two reversals to marine were inferred at the clades of Xenophthalmidae and Pinnotheridae, and Xenograpsidae and Cryptochiridae.

Using threshold models for both transition pathways, the best-fitting model was OU based on the lowest output Deviance Information Criterion (DIC) (**Table S8**). For the direct pathway, three shifts to non-marine grades were inferred at nodes: one in Ocypodidae, one at the base of the thoracotreme clade (Percnidae, Grapsidae, Plagusiidae, Mictyridae, Heloeciidae, Macrophthalmidae, Varunidae, Gecarcinidae, Sesarmidae, and Dotillidae), and one in Menippidae (**Figs. 4, S22**). If we consider grades 1 and 2 to be “semi-terrestrial” (similar to a character state from Tsang et al. [2022] referring to intertidal habitats), then the number of node origins for grades 3–5 is three or four: potentially twice in Ocypodidae, once in Mictyridae, and once at the base of the clade formed by (Gecarcinidae, Sesarmidae, and Dotillidae). Based on the estimated liabilities (i.e., thresholds of change required to transition to a different grade), it is 8– 36 times easier to move to grades 1 and 2 than to grades 3 and above (**Table S8**, **Fig. S23a**).

**Figure 4.**
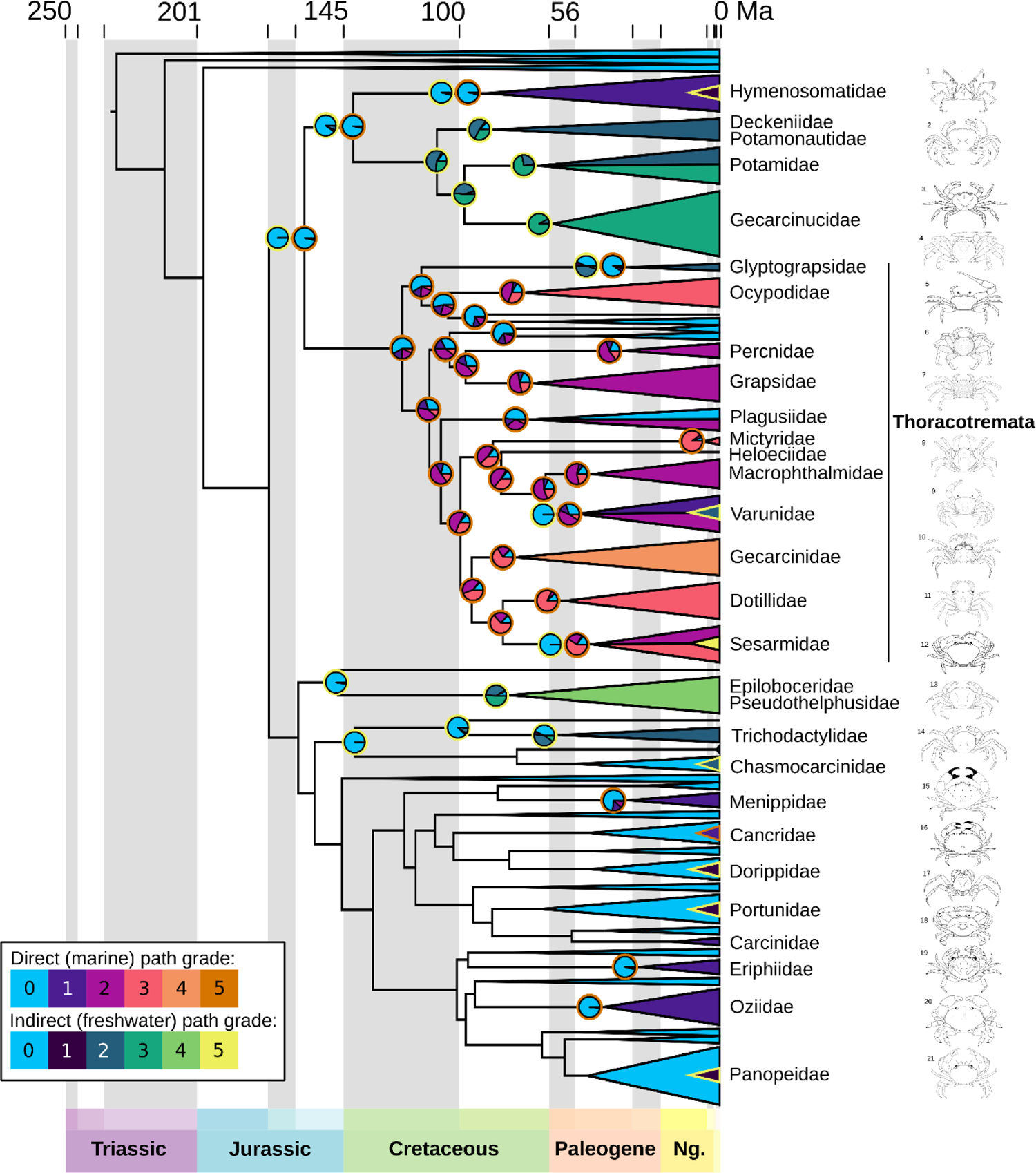
Composite of ancestral state reconstructions for the two transition pathways under best-fitting OU models in *ancThresh*, with fully marine clades (all families that are not labeled) reduced for clarity and outgroups removed. Legend for grades and colors representing each pathway at bottom left: fully marine crabs (grade 0), lower intertidal and estuaries (grade 1), upper intertidal and freshwater (grade 2), beaches and riverbanks (grade 3), and coastal forests and jungles (grades 4-5). Pies at nodes represent the estimated ancestral state with the outer ring indicating the pathway (at some nodes, both pathways are shown; when node is inferred marine, no pie is shown). Tip codings are based on estimates by family (Table S6), with the collapsed clades showing the color that represents the largest slice of their prior probabilities (split in the case of equal probabilities for two grades). For clades that have a small number of taxa in a grade from the opposite pathway, a small triangle is added. Line drawings at right (numbers corresponding to taxa in Table S7), by Javier Luque and Harrison Mancke.

Three reversals to marine or shifts to freshwater via this pathway (i.e., scored as grade 0) are inferred in: the clade of Xenograpsidae and Cryptochiridae, Plagusiidae, and Varunidae. In the case of Varunidae, the shift is to the indirect freshwater pathway (**Fig. 4**). Some tips that were coded with a majority of the prior probability failed to infer a shift at any nodes, such as Hymenosomatidae, Eriphiidae, and Oziidae (60% grade 1 for the former, 100% for the latter two), and Gecarcinidae (entirely grades 3–5, with 70% at grades 4 and 5).

For the indirect pathway, four shifts to non-marine grades were inferred at nodes: one at the base of the clades Potamoidea + Gecarcinucidae, and one each for Glyptograpsidae, Pseudothelphusoidea, and Trichodactylidae (**Figs. 4, S24**). If we consider grades 1 and 2 to be “semi-terrestrial”, then only Pseudothelphusoidea and Potamidae + Gecarcinucidae are inferred (each with nodes at grade 3). The liabilities indicate that it is extremely easy to move to grade 1, but 11 times harder to move to grade 2, and nearly 100 times harder to move to grade 3 (**Table S8**, **Fig. S23b**). Hymenosomatidae has 25% tip prior probabilities at grade 1, but no shifts were inferred. When analyzed as subclades, a transition was inferred from ancestrally freshwater to non-freshwater in Hymenosomatidae (**Fig. S25**), but we could not infer a transition from ancestrally terrestrial associated (i.e., intertidal/mangroves) to freshwater for Sesarmidae (**Fig. S26**).

## Discussion

### Relationships and divergence of true crabs

Previous molecular phylogenies of Brachyura have been constructed from eight to 10 Sanger loci (Tsang et al. 2014, 2022), mitochondrial genomes (Tan et al. 2019; Sun et al. 2022; Zhang et al. 2022), transcriptomics (Ma et al. 2019) and genomic target capture (Wolfe et al. 2019). However, the deep relationships among families and superfamilies remain uncertain, as the most extensive study (Tsang et al. 2014) sampled only 58 of 109 families with eight genes and low support at deep nodes, and more extensive gene sampling was coupled with even lower taxon sampling (Timm and Bracken-Grissom 2015; Wolfe et al. 2019). Although it is evident that the genes we and others have used are insufficient to resolve deep relationships even with current taxon sampling (**Fig. S14**), many regions of the tree are strongly supported. For example, the broadest strokes of our topological results (**Fig. 2**) contribute to the chorus of molecular and morphological analyses rejecting the monophyly of podotremes (e.g., Ahyong et al. 2007; Tsang et al. 2014, 2022; Luque et al. 2019a, 2019b; Tan et al. 2019) and heterotremes (e.g., Scholtz and Richter 1995; von Sternberg and Cumberlidge 2001; Tsang et al. 2014; Ma et al. 2019; Tan et al. 2019). Our divergence time estimates are older than most previous publications, except the deeper ages inferred by Wolfe et al. (2019) and the hypotheses of Guinot et al. (2019), but see below for evaluation. The relationships among thoracotremes were recently examined by Tsang et al. (2022).

As in their study, we find polyphyly of Ocypodoidea (fiddler, ghost crabs, and relatives), although our Grapsoidea (shore crabs, land crabs, and relatives) are separated into five clades, as opposed to four in Tsang et al. (2022). The main nodes where our results differ are: (1) the derived position of Dotillidae (sand bubbler crabs), (2) separation of the symbiotic groups Cryptochiridae (coral gall crabs) and Pinnotheridae (pea crabs), and (3) the position of Plagusiidae (different between ML and BI in our data: see **Fig. S12**). For points (2) and (3), we do recover weak support using all metrics. Ultimately, both studies use eight of the same loci.

Our data incorporates the nuclear rRNA genes, with relatively low phylogenetic informativeness above the family level (**Fig. S14**), yet both studies produce strong support at the base of thoracotreme families (note that Tsang et al. [2022] designated UF bootstraps > 90% as strong, which would add several nodes to our “strongly supported” category in **Figure 2** if we followed this metric). Finally, the backbone of thoracotreme phylogeny has been briefly addressed in two phylogenomic studies (Ma et al. 2019; Wolfe et al. 2019), and our results are similar to both, including the position of Sesarmidae (e.g., mangrove and vampire crabs) under models analyzing nucleotide data. A more robust understanding of internal thoracotreme relationships may be derived with additional phylogenomic data, but our results are sufficient to infer ancestral states, with the caveat of lower confidence at the aforeementioned nodes.

Polyphyly of the “primary” freshwater crab families (Deckeniidae, Epiloboceridae, Gecarcinucidae, Potamidae, Potamonautidae, Pseudothelphusidae, Trichodactylidae) has been found previously (e.g., von Sternberg and Cumberlidge 2001; Tsang et al. 2022). The inclusion of Hymenosomatidae (pillbox crabs) with the African and Eurasian freshwater groups in our results is novel, as it is the first analysis to incorporate this family in a larger tree. The grouping of African and Eurasian freshwater crabs with thoracotremes is otherwise fairly well supported in previous studies (e.g., Ma et al. 2019; Tan et al. 2019; Zhang et al. 2022). We contradict previous analyses, where the American Pseudothelphusoidea were closely related to the freshwater group (Tsang et al. 2014) and share a number of morphological synapomorphies (Cumberlidge et al. 2021). The American Trichodactylidae were more closely related to other heterotremes. Our results weakly support convergent origins of Pseudothelphusoidea (**Fig. 2**), leading to subsequent inference of separate transitions to freshwater. Divergence time estimates under all models push the origin of all primary freshwater groups except Trichodactylidae older than 66 Ma, with the African and Eurasian group over 100 Ma, consistent with a deeper cryptic history (Wolfe et al. 2019), particularly in non-marine environments (Tsang et al. 2014; Luque et al. 2021). Nevertheless, the 95% CIs of our divergence time estimates post-date the complete breakup of Pangaea (approximately 175 Ma), and do not support the Gondwanan origin of freshwater crabs (Klaus et al. 2011; Tsang et al. 2014) with these data.

Many of the relationships among families and superfamilies of the remaining heterotremes are not critical for the question of terrestriality. However, most of the deep relationships that we do observe, even with poor nodal support from ML, are congruent with the broad results retrieved from target capture of over 400 loci (Wolfe et al. 2019), except for the position of Menippidae (stone crabs). Our topologies for the relationships within superfamilies are largely congruent with previous Sanger data (Tsang et al. 2014), with some families exchanging places (e.g., Evans 2018; Mendoza et al. 2022) and some other groups with previous conflict still unresolved (Hultgren and Stachowicz 2008; Lai et al. 2014; Windsor and Felder 2014). The position of Dorippoidea nested well within heterotremes is consistent with previous molecular analysis (Tsang et al. 2014), but may confound hypotheses of early fossils that assume dorippoids are the earliest eubrachyuran branch (e.g., Guinot et al. 2019). One potentially novel result is the polyphyly of Bythograeoidea (hydrothermal vent crabs), that had not previously been included in a global analysis with as many genera (absent from Tsang et al. [2014]; no outgroups in concatenated analyses of Mateos et al. [2012]). Several non-monophyletic families were broken up by the insertion of closely related families, perhaps representing morphological groups that could be redefined as subfamilies (e.g., Lyreididae, Iphiculidae, potentially the cylodorippid genus *Tymolus*). In summary, we observe many similarities with previous analyses, and some new hypotheses that await improved molecular sampling before suggesting new systematic changes.

### Which divergence time estimates are reliable?

The factors that we investigated for different methodological choices were the calibration strategy and the clock model. Although divergence time estimates have only been visualized previously for only one dataset of echinoids with *chronospace* (Mongiardino Koch et al. 2022), we see significant differences in our data. Our results were impacted by the tested methods even more substantially than in the echinoid data, for which the results were sensitive to clock model choice, but not to substitution model or subsets of loci (Mongiardino Koch et al. 2022). In our data, a strong separation of variance of both clock model and calibration strategy were observed (**Fig. 3a-b**), with a similar pattern of posterior ages from both factors (**Fig. 3c**). However, the random local clock model, which accounts for the evolution of evolutionary rates in a clade-specific manner (Drummond and Suchard 2010; Ho and Duchêne 2014), failed to converge after months of runtime, so we cannot be certain of the ultimate effect on divergence time estimates.

We were initially interested in using FBD and BDSS because these tree models describe the processes of speciation, extinction, and fossil sampling that led to the true tree, and could be more accurate (Wright et al. 2022). A precursor method that incorporates fossil counts, but not directly into the tree model, has been previously used for lobsters (Bracken-Grissom et al. 2014). The gold standard for improved age accuracy may be to couple process-based tree models with joint inference of topology and ages, particularly total evidence tip dating (i.e., morphological character data from all incorporated fossils; Wright et al. 2022). Brachyura preserve abundant biomineralized hard parts, but the majority of fossils are dorsal carapaces and cheliped fragments. It is challenging to identify phylogenetically diagnostic characters in fragmented fossils due to substantial convergence in carapace and cheliped morphology (Guinot 2019; Luque et al. 2019b, 2021; Wolfe et al. 2021), and extensive missing data that could compromise their placement within crown groups. For these reasons, it was not possible to include a morphological matrix containing over 300 extant taxa, plus numerous fossils. Nevertheless, our evaluation of the brachyuran fossil record in the context of vetted crown and stem groups identified calibration fossils that were much older than previously appreciated (Luque et al. 2023). Most crown group families have exemplars preserved in the Eocene and Paleogene (66– 34 Ma), as well as an increasing number of well-preserved Cretaceous fossils (Ossó 2016; Luque et al. 2017, 2021, 2023).

Although records are abundant in the PBDB, brachyuran fossil sampling is not as evenly distributed as in many clades used as test cases for the performance of FBD and skyline models (e.g., Gavryushkina et al. 2014; Heath et al. 2014; Renner et al. 2016; Barido-Sottani et al. 2019; O’Reilly and Donoghue 2020). Various biases are observed in PBDB for crabs, including some families and early time slices with few or no fossils, and a geographic bias towards higher latitudes (we are continually working to improve the latter; e.g., Luque et al. 2017). Subsampling a fossil record using the unresolved FBD can be accurate with an evenly sampled clade such as cetaceans (Barido-Sottani et al. 2019), but can be highly inaccurate if the fossils have limited phylogenetic information (O’Reilly and Donoghue 2020). An analysis of mammals (Luo et al. 2021), however, found that fossil sampling density does not have linear effects on divergence time estimates. Altogether, it is difficult to generalize about the best models, unless prior knowledge about node ages is contradicted.

A major issue was estimating the root age of Meiura. Brachyura have a known ghost lineage, with stem group members *Eocarcinus* and *Eoprosopon* from the early Jurassic (approximately 190 Ma; Haug and Haug 2014; Hegna et al. 2020; Scholtz 2020; Wolfe et al. 2021). The oldest crown group brachyuran fossil occurrences in PBDB were also from the early Jurassic, representing less than 1% of total occurrences (including one occurrence in our subsample). Meanwhile, multiple modern anomuran families were already present in the late Jurassic (164–145 Ma; e.g., Fraaije et al. 2019, 2022; Robins and Klompmaker 2019), when their divergence likely took place (Bracken-Grissom et al. 2013; Wolfe et al. 2019). This strongly suggests the divergence of the common ancestor of Meiura, and probably of Brachyura, occurred at least by the very earliest Jurassic. Yet, the FBD and BDSS analyses, despite incorporating one Jurassic occurrence and a relatively representative subsample (**Table S4**), inferred impossibly young ages, as observed by O’Reilly and Donoghue (2020). Even the BDSS analysis including a time slice with no fossil sampling before the oldest occurrence (allowing a ghost lineage prior to the Jurassic: Culshaw et al. 2019; O’Reilly and Donoghue 2020) seemed to be a poor fit, and could not avoid estimating an unreasonably young root age (evident from the BDSS marginal prior in **Fig. S19a**). As such, given the data currently available for brachyurans, we recommend caution if using FBD models (and their extensions) to estimate divergences when a morphological matrix is unavailable, because such models may not perform according to expectations in all clades.

### How many times and when did crabs terrestrialize?

The number of estimated shifts to terrestriality (i.e., any shift from state 0) changes when considering the full diversity of known trait states within each family. We inferred seven shifts to non-marine lifestyles at nodes, fewer than Tsang et al. (2022), which estimated at least six transitions within Thoracotremata alone. Most shifts occur at well-supported nodes (**Figs. 2, 4**), although the common ancestor of (Thoracotremata, Hymenosomatidae, Potamoidea, Gecarcinucidae) has only moderate support, perhaps contributing to the uncertainty at this node with stochastic mapping (**Fig. S21**) and the differences we find between stochastic mapping and *ancThresh*. However, at least nine additional families have some proportion of their prior probability assigned to grades 1–2 (some all the way up to 100%, but most with lower probabilities), nested among marine sisters (**Fig. 4**), even though *ancThresh* does not infer a change at any node. Adding these brings the number of transitions to 16, distributed from Cretaceous to the last 10 myr. Although it is not included in the molecular phylogeny, there was likely another convergent transition to non-marine lifestyle in the Cretaceous fossil *Cretapsara* (Luque et al. 2021), resulting in a total of at least 17 terrestrialization events across at least 30 families.

The two or three losses of terrestriality that we estimate include nodes with poor support (the Cretaceous Xenograpsidae and Cryptochiridae, and the Eocene Plagusiidae). Character sequence reversals are rarely favored by threshold models (Revell 2014), so it is intriguing that we find these nodes. Across the tree of eukaryotic life, reversals to marine from non-marine lifestyles are more common than expected (Jamy et al. 2022), and more have been found in Thoracotremata (Tsang et al. 2022), so we could indeed be underestimating the phenomenon of returning to a marine environment.

Across plants and animals, more terrestrial species come from freshwater ancestors than directly from marine ancestors (Román-Palacios et al. 2022), a pathway we only observe potentially once, in Hymenosomatidae. We estimate only one instance of freshwater crabs evolving from intertidal ancestors, in Varunidae. Owing to the ordering of grades through the transition pathways, and the implementation of *ancThresh*, it is very challenging to infer these changes. The estimated liabilities are too high to infer freshwater Sesarmidae from an intertidal/terrestrial ancestor, although this is further complicated by all sesarmids starting from a higher base grade (i.e., starting at grade 2–3) within the two transition pathways, and possibly by the number of species that have convergently evolved arboreal lifestyles in mangroves (Fratini et al. 2005; Naruse and Ng 2020) and in freshwater derived habitats (Diesel 1989). These four clades are, however, the only brachyuran groups where the data suggest transitions between the main pathways, so the overall number of convergent events is not affected.

### Implications for early phases of arthropod terrestrialization

An outstanding question is the lifestyle of the common ancestor of the clade containing the majority of non-marine crabs: Thoracotremata, Hymenosomatoidea, Potamoidea, Gecarcinucoidea. Stochastic character mapping suggests the state of this node is uncertain (**Fig. S20**), although the most probable common ancestor may have been estuarine and likely lived in the Jurassic. It is also quite possible that the common ancestor was marine (**Fig. 4**), and direct and indirect pathways were established independently in the thoracotreme and freshwater clades. If estuarine, the Early Cretaceous common ancestor of Thoracotremata may have transitioned from the indirect pathway to an intertidal grade on the direct pathway, or alternatively this ancestor may have returned to fully marine life. Crabs in low grades of both transition pathways have different osmoregulatory adaptations (Watson-Zink 2021), so perhaps the thoracotreme common ancestor ecologically resembled some modern Hymenosomatidae (**Table S5**), or hypothetically resembled the only non-marine Cretaceous body fossil (which likely lived in brackish or fresh waters and may have been amphibious; Luque et al. 2021). The common ancestors at these early uncertain nodes could have had some degree of osmoregulatory ability, perhaps experiencing early development in marine or estuarine environments (Watson-Zink 2021).

The hypothetical common ancestor of many terrestrialized crabs, developed above, offers some lessons for understanding the Paleozoic terrestrialization of other arthropod groups. Despite a discrepancy of 100–150 myr between divergence time estimates (Cambrian) and body fossils (Silurian-Devonian), there are examples of Cambrian and Ordovician trace fossils that reveal limited excursions into non-marine environments, and perhaps more extensive life in marginal marine settings (e.g., Collette et al. 2010; Mángano et al. 2021; Buatois et al. 2022). Therefore, the fluidity of crab transitions into lower grades of terrestriality could hint at the types of adaptations other arthropods experienced in these early periods: osmoregulation and abbreviated and/or migratory larval development (Watson-Zink 2021). Other ecologies may resemble fiddler crabs that have adapted to coastal hypersaline environments, building their burrows well inland (Thurman 1984). Of course, other arthropods such as insects and arachnids continued on to surpass crabs in their terrestrial adaptations, as they became completely independent of water. Even the most terrestrial crab grades rely on water for, at the minimum, reproduction.

Additional instances of small numbers of taxa entering grades 1–2 through either pathway (e.g., Carcinidae, Panopeidae, freshwater Varunidae in **Fig. 4**) could provide some insights on the early stages of terrestrial adaptation. In particular, the above examples harbor some of the most persistent introduced crab taxa: *Carcinus maenas* (European green crab), *Hemigrapsus sanguineus* (Asian shore crab), and *Rhithropanopeus harrisii*. These species are notable for tolerating exceptionally wide-ranging salinities as larvae, and have wide and migratory habitat preferences as adults (e.g., Young and Elliott 2020). The introduced *Eriocheir sinensis* (Chinese mitten crab) can tolerate estuarine to full freshwater (Zhang et al. 2019).

Another example is the hymenosomatid *Halicarcinus planatus*, with a wide salinity tolerance that could help this species adapt and move into warming Antarctic waters (López-Farrán et al. 2021). Many other hymenosomatid genera have members in both freshwater and low salinity estuarine/mangrove habitats, and many have plastic osmoregulatory capabilities (Chuang and Ng 1994). Perhaps the ancestors of diverse non-marine groups originated with lifestyles similar to successful introduced species.

Groups sharing convergent morphological adaptations to higher grades of terrestriality, such as branchiostegal lungs in Gecarcinucidae, Gecarcinidae, Ocypodidae, and Pseudothelphusidae, or water-wicking setae in Gecarcinidae and Sesarmidae (Watson-Zink 2021 and **Table S5**), are deeply separated by over 100 Ma of evolution. Convergent terrestrial morphology in crabs, with likely pathways through a habitat gradient, perhaps with some traits of introduced taxa, could illuminate the hypothesis that arachnids convergently transitioned to terrestriality (Ballesteros et al. 2022). However, it is possible that horseshoe crabs (chelicerates, not decapods) returned to marine habitats even if they are phylogenetically nested within arachnids, as our crab analyses have inferred at least two reversals that involved likely intertidal (grade 2) ancestors.

## Conclusions

Herein, we inferred a large molecular phylogeny of true crabs, estimated divergence times that were older than previously thought, and estimated the number of transitions from marine to non-marine lifestyles. We found up to 17 convergent transitions through direct and indirect pathways, with at least three climbing to higher degrees of terrestrial adaptation. The most highly terrestrial clades were some of the oldest non-marine inferences in our data, with their common ancestors having diverged over 66 Ma. At least nine more recent events throughout the Cenozoic led to crabs living in intertidal and marginal marine environments, a shift that is estimated to be much easier based on lower threshold liability and likely fewer traits required. As instances of convergent evolution provide emerging models in the form of “natural experiments”, the framework we have developed to compare the gradient of adaptations will enable future research that aims to “predict” the constraints leading to repeated trait evolution and better understand the drivers of biodiversity across related groups.

## Author Contributions

JMW, KAC, DLF, HDBG conceived the project; STA, TYC, KHC, KAC, SRD, DLF, JWM, PKLN, EPT, NDP, RR, BPT, LMT, RW, AMW, HDBG provided samples; HDBG, EPT, RR, BPT, LMT, AMW extracted DNA and conducted sequencing; LB, HDBG curated sequence data; LB conducted phylogenetic analysis; JL vetted fossil calibrations; JMW, JBS developed and performed divergence time analysis; JMW, VMWZ coded natural history with input from JL, SRD, PKLN; JMW performed comparative methods and wrote the manuscript with input from all authors.

## Supplementary Material

Data available from the Dryad Digital Repository: https://doi.org/10.5061/dryad.tmpg4f52z

## Acknowledgments

We dedicate this paper to the memory of Christoph Schubart, whose contributions to brachyuran ecology, systematics, taxonomy and phylogeny we honor. The captain and crew of the R/V Pelican, R/V Weatherbird, R/V Urraca, R/V Hogarth and R/V Point Sur are recognized for assisting in specimen collections over the past 15 years. The staff and scientists at the US National Museum of Natural History: Smithsonian Institution, NIWA Invertebrate Collection, Natural History Museum of Los Angeles County, Universidad Nacional Autónoma de México, Smithsonian Tropical Marine Station: Bocas del Toro kindly assisted with shipping of loans, or hosting our visits. We thank Z. Nickell and many undergraduate students at Brigham Young University and Florida International University that assisted with museum curation, tissue plucking, DNA extractions and sequencing. We thank N. Mongiardino Koch and P. Milla Carmona for assistance with chronospace scripts, and L. Sallan for assistance with threshold models. Computations in this paper were run on the FASRC Cannon cluster supported by the FAS Division of Science Research Computing Group at Harvard University and the Florida International University High-Performance Computing Cluster (HPCC). This paper is contribution #1603 from the Coastlines and Oceans Division of the Institute of Environment at Florida International University, and contribution #223 from the UL Laboratory for Crustacean Research. Publication costs supported by a grant from the Wetmore Colles Fund.

## References

Ahyong S.T., Lai J.C.Y., Sharkey D., Colgan D.J., Ng P.K.L. 2007. Phylogenetics of the brachyuran crabs (Crustacea: Decapoda): The status of Podotremata based on small subunit nuclear ribosomal RNA. Molecular Phylogenetics and Evolution. 45:576–586.

Ballesteros J.A., Santibáñez-López C.E., Baker C.M., Benavides L.R., Cunha T.J., Gainett G., Ontano A.Z., Setton E.V.W., Arango C.P., Gavish-Regev E., Harvey M.S., Wheeler W.C., Hormiga G., Giribet G., Sharma P.P. 2022. Comprehensive species sampling and sophisticated algorithmic approaches refute the monophyly of Arachnida. Molecular Biology and Evolution. 39:msac021.

Bapst D.W., Wright A.M., Matzke N.J., Lloyd, G.T. 2016. Topology, divergence dates, and macroevolutionary inferences vary between different tip-dating approaches applied to fossil theropods (Dinosauria). Biology Letters. 12: 20160237.

Barido-Sottani J., Aguirre-Fernández G., Hopkins M.J., Stadler T., Warnock R.C. 2019. Ignoring stratigraphic age uncertainty leads to erroneous estimates of species divergence times under the fossilized birth-death process. Proceedings of the Royal Society B: Biological Sciences. 286:20190685.

Barido-Sottani J., Pohle A., De Baets K., Murdock D., Warnock R.C.M. 2022. Putting the F in FBD analyses: tree constraints or morphological data? BioRxiv.

Barido-Sottani J., van Tiel N., Hopkins M.J., Wright D.F., Stadler T., Warnock R.C.M. 2020. Ignoring fossil age uncertainty leads to inaccurate topology and divergence times in time calibrated tree inference. Frontiers in Ecology and Evolution. 8:183.

Benavides L.R., Edgecombe G.D., Giribet G. 2023. Re-evaluating and dating myriapod diversification with phylotranscriptomics under a regime of dense taxon sampling. Molecular Phylogenetics and Evolution. 178:107621.

Bernot J.P., Owen C.L., Wolfe J.M., Meland K., Olesen J., Crandall A. 2023. Major revisions in pancrustacean phylogeny and evidence of sensitivity to taxon sampling. Molecular Biology and Evolution. 40:msad175.

Bliss D.E. 1968. Transition from water to land in decapod crustaceans. American Zoologist. 8:355–392.

Bollback J.P. 2006. SIMMAP: stochastic character mapping of discrete traits on phylogenies. BMC Bioinformatics. 7:88.

Bouckaert R., Vaughan T.G., Barido-Sottani J., Duchêne S., Fourment M., Gavryushkina A., Heled J., Jones G., Kühnert D., De Maio N., Matschiner M., Mendes F.K., Müller N.F., Ogilvie H.A., du Plessis L., Popinga A., Rambaut A., Rasmussen D., Siveroni I., Suchard M.A., Wu C.-H., Xie D., Zhang C., Stadler T., Drummond A.J. 2019. BEAST 2.5: An advanced software platform for Bayesian evolutionary analysis. PLoS Comput Biol. 15:e1006650.

Bracken-Grissom H.D., Ahyong S.T., Wilkinson R.D., Feldmann R.M., Schweitzer C.E., Breinholt J.W., Bendall M., Palero F., Chan T.-Y., Felder D.L., Robles R., Chu K.-H., Tsang L.-M., Kim D., Martin J.W., Crandall K.A. 2014. The emergence of lobsters: Phylogenetic relationships, morphological evolution and divergence time comparisons of an ancient group (Decapoda: Achelata, Astacidea, Glypheidea, Polychelida). Systematic Biology. 63:457–479.

Bracken-Grissom H.D., Cannon M.E., Cabezas P., Feldmann R.M., Schweitzer C.E., Ahyong S.T., Felder D.L., Lemaitre R., Crandall K.A. 2013. A comprehensive and integrative reconstruction of evolutionary history for Anomura (Crustacea: Decapoda). BMC Evolutionary Biology. 13:128.

Buatois L.A., Davies N.S., Gibling M.R., Krapovickas V., Labandeira C.C., MacNaughton R.B., Mángano M.G., Minter N.J., Shillito A.P. 2022. The invasion of the land in deep time: Integrating Paleozoic records of paleobiology, ichnology, sedimentology, and geomorphology. Integrative and Comparative Biology. 62:297–331.

Castresana J. 2000. Selection of conserved blocks from multiple alignments for their use in phylogenetic analysis. Molecular Biology and Evolution. 17:540–552.

Chernomor O., von Haeseler A., Minh B.Q. 2016. Terrace aware data structure for phylogenomic inference from supermatrices. Systematic Biology. 6:997–1008.

Chuang C.T.N., Ng P.K.L. 1994. The ecology and biology of Southeast Asian false spider crabs (Crustacea: Decapoda: Brachyura: Hymenosomatidae). Hydrobiologia. 285:85–92.

Collette J.H., Hagadorn J.W., Lacelle M.A. 2010. Dead in their tracks--Cambrian arthropods and their traces from intertidal sandstones of Quebec and Wisconsin. PALAIOS. 25:475–486.

Culshaw V., Stadler T., Sanmartín I. 2019. Exploring the power of Bayesian birth-death skyline models to detect mass extinction events from phylogenies with only extant taxa. Evolution. 73:1133–1150.

Cumberlidge N., Johnson E.C., Leever E.M., Soma J.B., Ahles K.M., Kamanli S.A., Clark P.F. 2021. The significance of female dimorphic characters of primary freshwater crabs for the systematics of Eubrachyura (Crustacea: Decapoda: Brachyura). Journal of Crustacean Biology. 41:1–27.

Cumberlidge N., Ng P. 2009. Systematics, Evolution, and Biogeography of Freshwater Crabs. In: Martin J., Crandall K., Felder D., editors. Crustacean Issues: Decapod Crustacean Phylogenetics. Leiden: CRC Press. p. 491–508.

Cumberlidge N., Ng P.K.L., Yeo D.C.J., Magalhães C., Campos M.R., Alvarez F., Naruse T., Daniels S.R., Esser L.J., Attipoe F.Y.K., Clotilde-Ba F.-L., Darwall W., McIvor A., Baillie J.E.M., Collen B., Ram M. 2009. Freshwater crabs and the biodiversity crisis: Importance, threats, status, and conservation challenges. Biological Conservation. 142:1665–1673.

Davis K.E., De Grave S., Delmer C., Payne A.R.D., Mitchell S., Wills M.A. 2022. Ecological transitions and the shape of the Decapod Tree of Life. Integrative and Comparative Biology. 62:332–344.

Diesel R. 1989. Parental care in an unusual environment: *Metopaulias depressus* (Decapoda: Grapsidae), a crab that lives in epiphytic bromeliads. Animal Behaviour. 38:561–575.

Drummond A.J., Ho S.Y.W., Phillips M.J., Rambaut A. 2006. Relaxed phylogenetics and dating with confidence. PLoS Biology. 4:e88.

Drummond A.J., Suchard M.A. 2010. Bayesian random local clocks, or one rate to rule them all. BMC Biol. 8:114.

Edgecombe G.D., Strullu-Derrien C., Góral T., Hetherington A.J., Thompson C., Koch M. 2020. Aquatic stem group myriapods close a gap between molecular divergence dates and the terrestrial fossil record. Proc Natl Acad Sci USA. 117:8966–8972.

Evans N. 2018. Molecular phylogenetics of swimming crabs (Portunoidea Rafinesque, 1815) supports a revised family-level classification and suggests a single derived origin of symbiotic taxa. PeerJ. 6:e4260.

Felsenstein J. 2012. A comparative method for both discrete and continuous characters using the threshold model. American Naturalist. 179:145–156.

Fraaije R., Robins C., van Bakel B.W.M., Jagt J.W.M., Bachmayer F. 2019. Paguroid anomurans from the Tithonian Ernstbrunn Limestone, Austria – the most diverse extinct paguroid assemblage on record. Annalen des Naturhistorischen Museums in Wien. 121:257–289.

Fraaije R.H., Klompmaker A.A., Jagt J.W., Krobicki M., van Bakel B.W. 2022. A new, highly diverse paguroid assemblage from the Oxfordian (Upper Jurassic) of southern Poland and its environmental distribution. Neues Jahrbuch für Geologie und Paläontologie-Abhandlungen. 304:1–12.

Fratini S., Vannini M., Cannicci S., Schubart C.D. 2005. Tree-climbing mangrove crabs: a case of convergent evolution. Evolutionary Ecology Research. 7:219–233.

Gavryushkina A., Welch D., Stadler T., Drummond A.J. 2014. Bayesian Inference of Sampled Ancestor Trees for Epidemiology and Fossil Calibration. PLoS Computational Biology. 10:e1003919.

Guinot D. 2019. New hypotheses concerning the earliest brachyurans (Crustacea, Decapoda, Brachyura). Geodiversitas. 41:747–796.

Guinot D., Carbot-Chanona G., Vega F.J. 2019. Archaeochiapasidae n. fam., a new early Cenomanian brachyuran family from Chiapas, Mexico, new hypothesis on Lecythocaridae Schweitzer & Feldmann, 2009, and phylogenetic implications (Crustacea, Decapoda, Brachyura, Eubrachyura). Geodiversitas. 41:285–322.

Hartnoll R.G. 1988. Evolution, Systematics, and Geographical Distribution. In: Burggren W.W., McMahon B.R., editors. Biology of the Land Crabs. Cambridge: Cambridge University Press. p. 6–54.

Haug J.T., Haug C. 2014. *Eoprosopon klugi* (Brachyura)–the oldest unequivocal and most “primitive” crab reconsidered. Palaeodiversity. 7:149–158.

Heath T.A., Huelsenbeck J.P., Stadler T. 2014. The fossilized birth-death process for coherent calibration of divergence-time estimates. Proceedings of the National Academy of Sciences. 111:E2957–E2966.

Hegna T.A., Luque J., Wolfe J.M. 2020. The fossil record of the Pancrustacea. In: Poore G.C.B., Thiel M., editors. Evolution and Biogeography. Oxford: Oxford University Press. p. 21–52.

Ho S.Y.W., Duchêne S. 2014. Molecular-clock methods for estimating evolutionary rates and timescales. Molecular Ecology. 23:5947–5965.

Hoang D.T., Chernomor O., von Haeseler A., Minh B.Q., Vinh L.S. 2018. UFBoot2: Improving the ultrafast bootstrap approximation. Molecular Biology and Evolution. 35:518–522.

Hultgren K.M., Stachowicz J.J. 2008. Molecular phylogeny of the brachyuran crab superfamily Majoidea indicates close congruence with trees based on larval morphology. Molecular Phylogenetics and Evolution. 48:986– 996.

Jamy M., Biwer C., Vaulot D., Obiol A., Jing H., Peura S., Massana R., Burki F. 2022. Global patterns and rates of habitat transitions across the eukaryotic tree of life. Nat Ecol Evol. 6:1458–1470.

Kalyaanamoorthy S., Minh B.Q., Wong T.K.F., von Haeseler A., Jermiin L.S. 2017. ModelFinder: fast model selection for accurate phylogenetic estimates. Nature Methods. 14:587–589.

Katoh K., Standley D.M. 2013. MAFFT Multiple sequence alignment software version 7: Improvements in performance and usability. Molecular Biology and Evolution. 30:772–780.

Kearse M., Moir R., Wilson A., Stones-Havas S., Cheung M., Sturrock S., Buxton S., Cooper A., Markowitz S., Duran C., Thierer T., Ashton B., Meintjes P., Drummond A. 2012. Geneious Basic: An integrated and extendable desktop software platform for the organization and analysis of sequence data. Bioinformatics. 28:1647–1649.

Klaus S., Yeo D.C.J., Ahyong S.T. 2011. Freshwater crab origins—Laying Gondwana to rest. Zoologischer Anzeiger. 250:449–456.

Lai J.C.Y., Thoma B.P., Clark P.F., Felder D.L., Ng P.K.L. 2014. Phylogeny of eriphioid crabs (Brachyura, Eriphioidea) inferred from molecular and morphological studies. Zoologica Scripta. 43:52–64.

Lamsdell J.C., McCoy V.E., Perron-Feller O.A., Hopkins M.J. 2020. Air breathing in an exceptionally preserved 340-million-year-old sea scorpion. Current Biology. 30:4316–4321.

López-Farrán Z., Guillaumot C., Vargas-Chacoff L., Paschke K., Dulière V., Danis B., Poulin E., Saucède T., Waters J., Gérard K. 2021. Is the southern crab Halicarcinus planatus (Fabricius, 1775) the next invader of Antarctica? Glob Change Biol. 27:3487–3504.

Luo A., Zhang C., Zhou Q.-S., Ho S.Y.W., Zhu C.-D. 2021. Impacts of taxon-sampling schemes on Bayesian molecular dating under the unresolved Fossilized Birth-Death process. BioRxiv.

Luque J., Allison W.T., Bracken-Grissom H.D., Jenkins K.M., Palmer A.R., Porter M.L., Wolfe J.M. 2019a. Evolution of crab eye structures and the utility of ommatidia morphology in resolving phylogeny. BioRxiv.

Luque, J., Bracken-Grissom H.D., Ortega-Hernández J., Wolfe J.M. 2023.Fossil calibrations for molecular analyses and divergence time estimation for true crabs (Decapoda: Brachyura). BioRxiv.

Luque J., Feldmann R.M., Vernygora O., Schweitzer C.E., Cameron C.B., Kerr K.A., Vega F.J., Duque A., Strange M., Palmer A.R., Jaramillo C. 2019b. Exceptional preservation of mid-Cretaceous marine arthropods and the evolution of novel forms via heterochrony. Science Advances. 5:eaav3875.

Luque J., Schweitzer C.E., Santana W., Portell R.W., Vega F.J., Klompmaker A.A. 2017. Checklist of fossil decapod crustaceans from tropical America. Part I: Anomura and Brachyura. Nauplius. 25:1–85.

Luque J., Xing L., Briggs D.E.G., Clark E.G., Duque A., Hui J., Mai H., McKellar R.C. 2021. Crab in amber reveals an early colonization of nonmarine environments during the Cretaceous. Science Advances. 7:eabj5689.

Ma K.Y., Qin J., Lin C.-W., Chan T.-Y., Ng P.K.L., Chu K.H., Tsang L.M. 2019. Phylogenomic analyses of brachyuran crabs support early divergence of primary freshwater crabs. Molecular Phylogenetics and Evolution. 135:62–66.

Mángano M.G., Buatois L.A., Waisfeld B.G., Muñoz D.F., Vaccari N.E., Astini R.A. 2021. Were all trilobites fully marine? Trilobite expansion into brackish water during the early Palaeozoic. Proc. R. Soc. B. 288:20202263.

Mateos M., Hurtado L.A., Santamaria C.A., Leignel V., Guinot D. 2012. Molecular systematics of the deep-sea hydrothermal vent endemic brachyuran family Bythograeidae: A comparison of three Bayesian species tree methods. PLoS ONE. 7:e32066.

Mendoza J.C.E., Chan K.O., Lai J.C.Y., Thoma B.P., Clark P.F., Guinot D., Felder D.L., Ng P.K.L. 2022. A comprehensive molecular phylogeny of the brachyuran crab superfamily Xanthoidea provides novel insights into its systematics and evolutionary history. Molecular Phylogenetics and Evolution. 177:107627.

Minh B.Q., Hahn M.W., Lanfear R. 2020a. New methods to calculate concordance factors for phylogenomic datasets. Molecular Biology and Evolution. 37:2727–2733.

Minh B.Q., Schmidt H., Chernomor O., Schrempf D., Woodhams M., von Haeseler A., Lanfear R. 2020b. IQ-TREE 2: New models and efficient methods for phylogenetic inference in the genomic era. Molecular Biology and Evolution. 37:1530–1534.

Mongiardino Koch N., Thompson J.R., Hatch A.S., McCowin M.F., Armstrong A.F., Coppard S.E., Aguilera F., Bronstein O., Kroh A., Mooi R., Rouse G.W. 2022. Phylogenomic analyses of echinoid diversification prompt a re-evaluation of their fossil record. eLife. 11:e72460.

Naruse T., Ng P.K.L. 2020. Revision of the sesarmid crab genera Labuanium Serène and Soh, 1970, Scandarma Schubart, Liu and Cuesta, 2003 and Namlacium Serène and Soh, 1970 (Crustacea: Decapoda: Brachyura: Sesarmidae), with descriptions of four new genera and two new species. Journal of Natural History. 54:445–532.

Ng P.L.K., Guinot D., Davie P.J.F. 2008. Systema Brachyurorum: Part I. An annotated checklist of extant brachyuran crabs of the world. Raffles Bulletin of Zoology. 17:1–286.

O’Reilly J.E., Donoghue P.C.J. 2020. The effect of fossil sampling on the estimation of divergence times with the Fossilized Birth–Death process. Systematic Biology. 69:124–138.

Ossó À. 2016. Eogeryon elegius n. gen. and n. sp. (Decapoda: Eubrachyura: Portunoidea), one of the oldest modern crabs from late Cenomanian of the Iberian Peninsula. Boletín de la Sociedad Geológica Mexicana. 68:231– 246.

Parham J.F., Donoghue P.C.J., Bell C.J., Calway T.D., Head J.J., Holroyd P.A., Inoue J.G., Irmis R.B., Joyce W.G., Ksepka D.T., Patane J.S.L., Smith N.D., Tarver J.E., van Tuinen M., Yang Z., Angielczyk K.D., Greenwood J.M., Hipsley C.A., Jacobs L., Makovicky P.J., Muller J., Smith K.T., Theodor J.M., Warnock R.C.M., Benton M.J. 2012. Best practices for justifying fossil calibrations. Systematic Biology. 61:346– 359.

Poore G.C.B., Ahyong S.T. 2023. Marine decapod Crustacea: a guide to the families and genera of the world. CSIRO Publishing.

Powers L.W., Bliss D.E. 1983. Terrestrial adaptations. In: Bliss D.E., editor. The Biology of Crustacea. Academic Press. p. 271–334.

Rambaut A., Drummond A.J., Xie D., Baele G., Suchard M.A. 2018. Posterior summarisation in Bayesian phylogenetics using Tracer 1.7. Systematic Biology. 67:901–904.

Renner S.S., Grimm G.W., Kapli P., Denk T. 2016. Species relationships and divergence times in beeches: new insights from the inclusion of 53 young and old fossils in a birth–death clock model. Philosophical Transactions of the Royal Society B: Biological Sciences. 371:20150135.

Revell L.J. 2012. phytools: an R package for phylogenetic comparative biology (and other things). Methods in Ecology and Evolution. 3:217–223.

Revell L.J. 2014. Ancestral character estimation under the threshold model from quantitative genetics. Evolution. 68:743–759.

Robin N., van Bakel B.W.M., Hyžný M., Cincotta A., Garcia G., Charbonnier S., Godefroit P., Valentin X. 2019. The oldest freshwater crabs: claws on dinosaur bones. Scientific Reports. 9:20220.

Robins C.M., Klompmaker A.A. 2019. Extreme diversity and parasitism of Late Jurassic squat lobsters (Decapoda: Galatheoidea) and the oldest records of porcellanids and galatheids. Zoological Journal of the Linnean Society. 187:1131–1154.

Román-Palacios C., Moraga-López D., Wiens J.J. 2022. The origins of global biodiversity on land, sea and freshwater. Ecology Letters. 25:1376–1386.

Ronquist F., Teslenko M., van der Mark P., Ayres D.L., Darling A., Hohna S., Larget B., Liu L., Suchard M.A., Huelsenbeck J.P. 2012. MrBayes 3.2: Efficient Bayesian phylogenetic inference and model choice across a large model space. Systematic Biology. 61:539–542.

Sallan L., Friedman M., Sansom R.S., Bird C.M., Sansom I.J. 2018. The nearshore cradle of early vertebrate diversification. Science. 362:460–464.

Scholtz G. 2020. Eocarcinus praecursor Withers, 1932 (Malacostraca, Decapoda, Meiura) is a stem group brachyuran. Arthropod Structure & Development. 59:100991.

Schubart C.D., Neigel J.E., Felder D.L. 2000. The use of the mitochondrial 16S rDNA gene for phylogenetics and biogeographic studies of Crustacea. Crustacean Issues: The Biodiversity Crisis and Crustacea. Boca Raton: CRC Press. p. 817–830.

Song H., Buhay J.E., Whiting M.F., Crandall K.A. 2008. Many species in one: DNA barcoding overestimates the number of species when nuclear mitochondrial pseudogenes are coamplified. Proc. Natl. Acad. Sci. U.S.A. 105:13486–13491.

Spears T., Abele L.G. 1998. Crustacean phylogeny inferred from 18S rDNA. In: Fortey R.A., Thomas R.H., editors. Arthropod Relationships. Dordrecht: Springer. p. 169–187.

Stadler T. 2010. Sampling-through-time in birth–death trees. Journal of Theoretical Biology. 267:396–404.

Stadler T., Kuhnert D., Bonhoeffer S., Drummond A.J. 2013. Birth-death skyline plot reveals temporal changes of epidemic spread in HIV and hepatitis C virus (HCV). Proceedings of the National Academy of Sciences. 110:228–233.

von Sternberg R., Cumberlidge N. 2001. Notes on the position of the true freshwater crabs within the brachyrhynchan Eubrachyura (Crustacea: Decapoda: Brachyura). Hydrobiologia. 449:21–39.

Sun S., Jiang W., Yuan Z., Sha Z. 2022. Mitogenomes provide insights into the evolution of Thoracotremata (Brachyura: Eubrachyura). Front. Mar. Sci. 9:848203.

Tan M.H., Gan H.M., Lee Y.P., Bracken-Grissom H., Chan T.-Y., Miller A.D., Austin C.M. 2019. Comparative mitogenomics of the Decapoda reveals evolutionary heterogeneity in architecture and composition. Scientific Reports. 9:10756.

Thurman C.L. 1984. Ecological notes on fiddler crabs of South Texas, with special reference to *Uca subcylindrica*. Journal of Crustacean Biology. 4:665–681.

Timm L., Bracken-Grissom H.D. 2015. The forest for the trees: evaluating molecular phylogenies with an emphasis on higher-level Decapoda. Journal of Crustacean Biology. 35:577–592.

Tsang C.T.T., Schubart C.D., Chu K.H., Ng P.K.L., Tsang L.M. 2022. Molecular phylogeny of Thoracotremata crabs (Decapoda, Brachyura): Toward adopting monophyletic superfamilies, invasion history into terrestrial habitats and multiple origins of symbiosis. Molecular Phylogenetics and Evolution. 177:107596.

Tsang L.M., Ma K.Y., Ahyong S.T., Chan T.-Y., Chu K.H. 2008. Phylogeny of Decapoda using two nuclear protein-coding genes: Origin and evolution of the Reptantia. Molecular Phylogenetics and Evolution. 48:359–368.

Tsang L.M., Schubart C.D., Ahyong S.T., Lai J.C.Y., Au E.Y.C., Chan T.-Y., Ng P.K.L., Chu K.H. 2014. Evolutionary history of true crabs (Crustacea: Decapoda: Brachyura) and the origin of freshwater crabs. Molecular Biology and Evolution. 31:1173–1187.

Watson-Zink V.M. 2021. Making the grade: Physiological adaptations to terrestrial environments in decapod crabs. Arthropod Structure & Development. 64:101089.

Windsor A.M., Felder D.L. 2014. Molecular phylogenetics and taxonomic reanalysis of the family Mithracidae MacLeay (Decapoda: Brachyura: Majoidea). Invertebrate Systematics. 28:145–173.

Wolfe J.M., Breinholt J.W., Crandall K.A., Lemmon A.R., Moriarty Lemmon E., Timm L.E., Siddall M.E., Bracken-Grissom H.D. 2019. A phylogenomic framework, evolutionary timeline, and genomic resources for comparative studies of decapod crustaceans. Proceedings of the Royal Society B: Biological Sciences. 286:20190079.

Wolfe J.M., Daley A.C., Legg D.A., Edgecombe G.D. 2016. Fossil calibrations for the arthropod Tree of Life. Earth-Science Reviews. 160:43–110.

Wolfe J.M., Luque J., Bracken-Grissom H.D. 2021. How to become a crab: Phenotypic constraints on a recurring body plan. BioEssays. 43:2100020.

WoRMS. 2022. Brachyura. Accessed at: https://www.marinespecies.org/aphia.php?p=taxdetails&id=106673 on 12-08-2022.

Wright A.M., Bapst D.W., Barido-Sottani J., Warnock R.C.M. 2022. Integrating fossil observations into phylogenetics using the Fossilized Birth–Death model. Annu. Rev. Ecol. Evol. Syst. 53:251–273.

Yeo D.C.J., Ng P.K.L., Cumberlidge N., Magalhães C., Daniels S.R., Campos M.R. 2008. Global diversity of crabs (Crustacea: Decapoda: Brachyura) in freshwater. Hydrobiologia. 595:275–286.

Young A.M., Elliott J.A. 2020. Life history and population dynamics of green crabs (*Carcinus maenas*). Fishes. 5:44.

Zhang Y., Gong L., Lu X., Miao Z., Jiang L., Liu B., Liu L., Li P., Zhang X., Lü Z. 2022. Comparative mitochondrial genome analysis of Varunidae and its phylogenetic implications. Acta Oceanol. Sin. 41:119– 131.

Zhang Z., Capinha C., Weterings R., McLay C.L., Xi D., Lü H., Yu L. 2019. Ensemble forecasting of the global potential distribution of the invasive Chinese mitten crab, *Eriocheir sinensis*. Hydrobiologia. 826:367–377.

