## Supplementary material for "Convergent adaptation of true crabs (Decapoda: Brachyura) to a gradient of terrestrial environments": Text S1

**Text S1.** Supplemental methods.
**Text S2.** References associated with natural history codings in Table S5.

**Figure S1.** Gene tree for 12S. Nodes are unlabeled as gene trees were calculated as part of the concordance factor analysis (labels visualized in Figure S11). Branches colored by superfamily, as for Figure 2.
**Figure S2.** Gene tree for 16S. Branches colored by superfamily, as for Figure 2.
**Figure S3.** Gene tree for 18S. Branches colored by superfamily, as for Figure 2.
**Figure S4.** Gene tree for 28S. Branches colored by superfamily, as for Figure 2.
**Figure S5.** Gene tree for AK. Branches colored by superfamily, as for Figure 2.
**Figure S6.** Gene tree for enolase. Branches colored by superfamily, as for Figure 2.
**Figure S7.** Gene tree for GADPH. Branches colored by superfamily, as for Figure 2.
**Figure S8.** Gene tree for H3. Branches colored by superfamily, as for Figure 2.
**Figure S9.** Gene tree for NaK. Branches colored by superfamily, as for Figure 2.
**Figure S10.** Gene tree for PEPCK. Branches colored by superfamily, as for Figure 2.
**Figure S11.** Full phylogenetic hypothesis for Brachyura based on the topology from the ML concatenated analysis. Values at nodes represent ultrafast bootstraps / gene concordance factors (gCF) by partition / updated site concordance factors (sCF). Branches colored by superfamily, as for Figure 2.
**Figure S12.** Full phylogenetic hypothesis for Brachyura based on the topology from the Bayesian concatenated analysis. Values at nodes represent posterior probabilities. Branches colored by superfamily, as for Figure 2.
**Figure S13.** Plots of concordance factors versus UF bootstrap values for the ML concatenated tree. (**A**) Gene concordance factors by partition. (**B**) Updated site concordance factors.
**Figure S14.** Results of phylogenetic informativeness profiling for 10 Sanger genes.
**Figure S15.** Sensitivity of divergence time estimates to inference strategy plotted with *chronospace*, with outgroup taxa removed from these analyses. This plot shows between-group principal component analysis (bgPCA) separating chronograms by calibration strategy, for only relaxed lognormal clock models (all fully converged).
**Figure S16.** Full maximum clade credibility divergence time estimates for Brachyura as depicted in Figure 2. Posterior ages were estimated in BEAST2 using 36 vetted node calibrations (represented by numbered circles following the order in Text S1) and the topology fixed from Figure S11, a birth-death tree prior, and relaxed lognormal clock model. Horizontal shaded bars represent 95% HPD intervals. Branches colored by superfamily, as for Figure 2.

**Figure S17.** Full maximum clade credibility divergence time estimates for Brachyura using the FBD. Posterior ages were estimated in BEAST2 using 328 calibrations from PBDB added as rogues to the topology fixed from Figure S11, an FBD tree prior, and relaxed lognormal clock model. Horizontal shaded bars represent 95% HPD intervals. Branches colored by superfamily, as for Figure 2.

**Figure S18.** Full maximum clade credibility divergence time estimates for Brachyura using BDSS. Posterior ages were estimated in BEAST2 using 328 calibrations from PBDB added as rogues to the topology fixed from Figure S11, a BDSS tree prior, and relaxed lognormal clock model. Horizontal shaded bars represent 95% HPD intervals. Branches colored by superfamily, as for Figure 2.

**Figure S19.** Comparison of posterior probability distributions for divergence time analyses under different calibration strategies in BEAST2, and the same analyses under the marginal prior (removing sequence data). The posterior analyses are shaded; marginal priors are superimposed on the same axes with a heavy line of the same color. (**A**) Root; (**B**) 19 most sensitive nodes, calculated based on Figure 3A.

**Figure S20.** Inferred posterior birth rate through time, from BDSS analysis.

**Figure S21.** Summary of 500 stochastic character maps coding tips in a simplified form as: fully marine, terrestrial (through direct marine pathway), and freshwater, under the best-fitting all rates different (ARD) model. These codings are based on the transition pathways scored from the literature supported data in Table S5.

**Figure S22.** Ancestral state reconstruction for the sequence from fully marine to terrestrial through direct marine pathway, under the best-fitting OU model in *ancThresh*. The tip codings are based on estimates by family as coded in Table S6, with their prior probabilities visualized as small pies at tips.

**Figure S23.** Posterior density of liabilities estimated in *ancThresh*, indicating the thresholds of change required to shift character states. As in Sallan et al. (2018), the threshold for exiting marine (grade 0) is held at 0, and values for exiting grade 5 are infinity as there are no subsequent states to transition into. (**A**) Sequence from fully marine to terrestrial through direct marine pathway, under OU model, and (**B**) Sequence from fully marine to terrestrial through indirect freshwater pathway, under OU model.

**Figure S24.** Ancestral state reconstruction for the sequence from fully marine to terrestrial through indirect freshwater pathway, under the best-fitting OU model in *ancThresh*. The tip codings are based on estimates by family as coded in Table S6, with their prior probabilities visualized as small pies at tips.

**Figure S25.** Ancestral state reconstructions for transitions between pathways for subclade Hymenosomatidae, under the OU model in *ancThresh*. (**A**) From ancestrally freshwater to direct pathway. In this case, state 0 reflects fully marine plus terrestrial prior probabilities. (**B**) From terrestrial through the direct pathway, to freshwater. Posterior density of liabilities from (**C**) ancestrally freshwater to direct pathway and (**D**) terrestrial through the direct pathway, to freshwater.

**Figure S26.** Ancestral state reconstructions for transitions between pathways for subclade Sesarmidae, under the OU model in ancThresh. (**A**) From ancestrally terrestrial to indirect freshwater pathway. In this case, state 0 reflects fully marine plus terrestrial prior probabilities. (**B**) From freshwater through the indirect pathway, to terrestrial. Posterior density of liabilities from (**C**) ancestrally terrestrial to indirect freshwater pathway and (**D**) freshwater through the indirect pathway, to terrestrial.

**Table S1.** Description of the pathways, grades, and characteristic traits of terrestriality, following Watson-Zink (2021).
**Table S2.** Sample information for new and previous sequence data.
**Table S3.** Fossil node calibrations with details following Wolfe et al. (2016, 2019) and elaborated in Luque et al. (2023). The topology defining the nodes is the concatenated ML tree.
**Table S4.** Properties of the 10% subsample of fossil occurrences from PBDB, ordered by age.
**Table S5.** Discrete coded terrestriality grades for all sequenced taxa, with evidence for each trait supported by literature. Transition pathway 0 represents fully marine lifestyle, 1 represents transition to terrestriality through marine pathway, 2 represents transition through freshwater pathway. Grades as in Watson-Zink (2021) and Table S1.
**Table S6.** Prior probabilities of terrestriality grades estimated for each family, accounting for states from taxa that are not sequenced, based on WoRMS. Transition pathway 0 represents fully marine lifestyle, 1 represents transition to terrestriality through marine pathway, 2 represents transition through freshwater pathway. Grades as in Watson-Zink (2021) and Table S1.
**Table S7.** Taxonomic key for line drawings in Figures 2 and 4.
**Table S8.** Model parameters estimated by *ancThresh*. Means reported for best-fitting model only (or OU model only for subclades), after excluding 20% burnin. As in Sallan et al. (2018), the threshold for exiting marine (grade 0) is held at 0, and values for exiting grade 5 are infinity as there are no subsequent states to transition into.

#### Text S1. Supplemental Methods

*Taxon sampling*

This dataset represents over 15 years of sampling crabs in the field or from museum collections (**Table S2**). The final dataset includes 88 families, 263 genera, 333 species and 338 individuals within the infraorder Brachyura, as well as six outgroups for a total of 344 tips, 45% representing new sequence data, with the remainder obtained directly from GenBank (**Table S2**). All taxonomy was checked against expert compilations (Ng et al. 2008; Poore and Ahyong 2023, WoRMS 2022).

*Gene selection*

A total of 10 genes were selected based on previous phylogenetic research of decapods (Spears and Abele 1998; Schubart et al. 2000; Tsang et al. 2008; Bracken-Grissom et al. 2013; Tsang et al. 2014). These included two mitochondrial ribosomal RNA (rRNA) coding genes, 12S (small ribosomal subunit) and 16S (large ribosomal subunit) as well as two nuclear rRNA genes, 18S (small ribosomal subunit) and 28S (large ribosomal subunit). The six nuclear protein-coding genes NaK (sodium–potassium ATPase α-subunit), PEPCK (phosphoenolpyruvate carboxykinase), GADPH (glyceraldehyde 3-phosphate dehydrogenase), H3 (histone 3), enolase (phosphopyruvate hydratase), and AK (arginine kinase) were also included. A minimum of two genes were required for each taxon included in the analysis, with an average of seven genes available per tip.

*DNA extraction, PCR, and sequencing*

Genomic DNA was extracted from the gills, abdomen, pereopod, or pleopod, using the Qiagen DNeasy® Blood and Tissue Kit (Cat. No. 69582), QIAamp DNA Mini Kit (Cat. No. 51304) or QIAamp DNA Micro Kit (Cat. No. 56304). Gene regions were amplified with polymerase chain reaction (PCR) using one or more sets of primers. Annealing temperatures and primer sequences are as follows: 12S rRNA (325 bp, Buhay et al. 2007), 16S rRNA (339 bp, Simon et al. 1994; Crandall and Fitzpatrick 1996; Schubart et al. 2002), 18S rRNA (1798 bp, Medlin et al. 1988; Whiting et al. 1997; Whiting 2002; Bracken et al. 2009), 28S rRNA (1892 bp, Whiting et al. 1997; Whiting 2002; Palero et al. 2008), histone 3 (H3) (336 bp, Colgan et al. 1998), NaK (606 bp, Tsang et al. 2008), PEPCK (582 bp, Tsang et al. 2008), GADPH (609 bp, Tsang et al. 2011), enolase (399 bp, Tsang et al. 2011), and AK (630 bp, Tsang et al. 2011).

PCR amplifications were performed in 25–50 µl volumes containing 1 U of Taq polymerase (HotMaster, AccuPrime or REDTaq), PCR Buffer, 2.5 mM of deoxyribonucleotide triphosphate mix (dNTPs), 0.5 µM forward and reverse primer, and 30–100 ng extracted DNA. The thermal profile used an initial denaturation for 1 min at 94ºC followed by 30–40 cycles of 30 sec-1 min at 94ºC, 45 sec-1 min at 46-58ºC (depending on gene region), 1 min at 72ºC and a final extension of 7 min at 72ºC. PCR products were purified using filters (PrepEase™ PCR Purification 96-well Plate Kit, USB Corporation) and sequenced with ABI BigDye® terminator mix (Applied Biosystems, Foster City, CA, USA). Cycle Sequencing reactions were performed in an Applied Biosystems 9800 Fast Thermal Cycler (Applied Biosystems, Foster City, CA, USA), and sequencing products were run (forward and reverse) on an ABI 3730xl DNA Analyzer 96-capillary automated sequencer.

*Phylogenetic analysis*

Sequences (n = 2251) were assembled and trimmed within Geneious Prime (Kearse et al. 2012). All protein coding genes (AK, enolase, GADPH, H3, NaK, PEPCK) were checked for pseudogenes following Song et al. (2008). Ribosomal RNA genes (12S, 16S, 18S, 28S), were individually aligned using the MAFFT via local iterative refinement algorithms in Geneious v.2021.0.1 (Kearse et al. 2012; Katoh and Standley 2013). Protein-coding genes were individually aligned in Geneious using the MAFFT global iterative refinement algorithm. To remove regions of questionable homology, rRNA alignments were masked in GBlocks v.0.91b (Castresana 2000) under "less stringent" parameters.

Alignments were concatenated in Geneious Prime. Best-fitting partitions were selected by an edge-linked proportional model (Chernomor et al. 2016) and substitution models with ModelFinder (Kalyaanamoorthy et al. 2017) in IQ-TREE v.2.1.2 (Minh et al. 2020b). The best-fitting scheme was used to estimate the concatenated maximum likelihood (ML) phylogeny, also in IQ-TREE, using 10 independent runs with perturbation strength 0.2 and stopping at 500 iterations, selecting the best topology by the highest log likelihood (Zhou et al. 2018). Ultrafast (UF) bootstrap values were calculated from 1000 replicates using the -bnni flag (Hoang et al. 2018). Gene and site concordance factors (gCF and sCF, respectively) were calculated in IQ-TREE v.2.2.2 (Minh et al. 2020a; Mo et al. 2022), simultaneously generating individual gene trees using ML. We further explored the resolution of individual loci evolving at different rates by using a metric for per-site phylogenetic informativeness (Townsend 2007) estimated by the online PhyDesign program (http://phydesign.townsend.yale.edu/; López-Giráldez and Townsend 2011) using the DNArates substitution model.

We conducted a Bayesian inference (BI) analysis of the concatenated loci using MrBayes v.3.2.7 (Ronquist et al. 2012). Two runs (analyses starting from different start trees) and four chains (three heated and one cold per run) were run for 35 million generations with 25% burnin. Convergence was assessed by reaching effective sample size >200 for every parameter, and by evaluating posterior distributions in Tracer v.1.7.1 (Rambaut et al. 2018).

*Fossil calibration and divergence time inference*

We compared two strategies for fossil calibration: (1) 36 newly vetted node calibrations (Luque et al. 2023), and (2) 328 fossil occurrences from the Paleobiology Database (PBDB; http://paleobiodb.org/). For node calibration, all calibrations followed best practices regarding specimen data, morphological diagnosis, and stratigraphy (Parham et al. 2012; Wolfe et al. 2016; details in **Table S3** and Luque et al. 2023), and were assigned to a crown group node at the family level or higher. All internal calibration age distributions were uniform. We applied a root prior with a fossil-informed lognormal distribution (mean = 282 Ma, standard deviation = 0.25, offset = 25; based on the same node in Wolfe et al. [2019]). This node dating strategy used a birth-death tree prior.

We downloaded fossil occurrences from the PBDB on March 23, 2022, for Brachyura at family-level taxonomic resolution, excluding uncertain genera and species, with accepted names only, from Jurassic to Pleistocene. Issues are known with the entry of decapod fossil taxonomy in the PBDB, primarily with typos falsely creating multiple taxa for different occurrences that should belong to the same taxon (M. Uhen and M. Clapham, pers. comm.). Our efforts to improve the accuracy of PBDB data included: spot checking occurrences, resolving all family level taxonomy (wherever we found errors) in the full sample of brachyuran occurrences, resolving genera in the subsampled occurrences (see below), and manually removing from our downloads 30 families that could not be placed in our tree (extant families without molecular data, or extinct families with no known phylogenetic context). We randomly subsampled 10% of the 3,276 remaining occurrences, resulting in a computationally tractable 328 occurrences. The subsample was a reasonable sample of ages (including the oldest possible age, at 201.3-145 Ma), represented 69% of fossil families, and slightly overrepresented non-marine paleoenvironments (details in **Table S4**). All fossil occurrences were assigned age ranges from the PBDB, each with a uniform distribution (Barido-Sottani et al. 2019, 2020). To incorporate these fossil samples as part of the inferred evolutionary process, we used the unresolved time-homogeneous fossilized birth-death (FBD) tree prior (Stadler 2010; Heath et al. 2014; O'Reilly and Donoghue 2020), conditioned on the same root prior as above. The diversification rate was assigned a uniform prior (0, infinity) to reflect the unknown proportion of extinct relative to extant species, and an initial rate of 0.01. The turnover rate and sampling proportion were both assigned uniform priors with distribution (0, 1). For rho, the proportion of living species sampled, the value was fixed at 0.04 based on 7,636 valid brachyuran species in WoRMS (WoRMS 2021).

To reflect the complete absence of brachyuran fossils from earlier than the Jurassic, which represents a known ghost lineage when compared to the diversity and abundance of outgroup anomuran fossils, thus earlier diversification of the sister group (Hegna et al. 2020; Wolfe et al. 2021), we also analyzed the fossil occurrence calibration set using a birth-death skyline tree prior with sequential sampling (BDSS; Stadler et al. 2013; Culshaw et al. 2019). Fossil sampling proportion was modeled as time-heterogeneous with time slices before and after the oldest fossil sample (initial values of 0 and 0.1, respectively), and a uniform prior distribution (0, 1). This was done using the TreeSlicer function from the *skylinetools* package (https://github.com/laduplessis/skylinetools). While we used the same root prior, the origin time was estimated with a uniform prior (182, 372) and an initial value of 325 Ma (Wolfe et al. 2019). The reproductive number R0 (i.e., birth rate/total death rate) had a dimension of 5; this parameter and the total death rate were both assigned lognormal prior distributions.

All divergence time analyses were conducted in BEAST2 v.2.6.7 (Bouckaert et al. 2019) using a fixed starting tree derived from our ML concatenated results. When using fossil occurrences (FBD and BDSS analyses only), these were added to the starting tree as "rogues" (able to move within pre-assigned family level constraints following Barido-Sottani et al. [2022], as most decapod fossils are fragmentary and cannot be confidently assigned). All analyses linked the substitution models selected by ModelFinder, and the clock models of those partitions (4 categories total). For each calibration strategy and tree prior, we compared two clock models: relaxed lognormal (Drummond et al. 2006) and random local (Drummond and Suchard 2010), using lognormal priors on the mean branch rates (lower and upper values of 0.001 and 1, respectively). Analyses used four to six independent runs for at least 450 million generations with 25% burnin, visualized as a maximum clade credibility tree. Convergence was assessed as above. Having explored a total of six divergence time analyses, we visualized the results from different parameter sets using *chronospace* scripts (Mongiardino Koch et al. 2022), which provide a multidimensional representation of inferred node ages that can be broken down by different factors and models.

*Ancestral state reconstruction*

To code character states representing a gradient of terrestriality, we used a modified version of the trait-by-environment associations defined by Watson-Zink (2021), additional details in **Table S1**. Distinct transition pathways were defined for marine and freshwater routes. For each pathway, we used five of the six grades of increasing terrestriality; the highest level does not appear in our data (Watson-Zink 2021). We added a grade 0, to indicate that the ancestral state for all crabs is fully marine (**Fig. 1a-e**). Using this framework, we coded discrete grades of terrestriality following two schemes. The first scheme coded the taxa that were sequenced in our molecular phylogeny and required detailed justification from natural history literature (**Table S5**, **Text S2**). Justifications accounted for available data on adult habitat, osmoregulatory status, larval developmental strategy, primary respiratory structure, water-wicking setae, burrow type, and diurnal activity period (Watson-Zink 2021). As our phylogeny sampled 4% of brachyuran species as tips, we also constructed a scheme to estimate grades for unsampled species for which the phylogenetic positions are unknown. For this scheme, we downloaded all taxonomic data, including non-marine taxa, from WoRMS as of June 8, 2021. For families sampled in our tree, we used the WoRMS data to estimate the number of species that fall into each grade (without detailed literature survey per species) and accordingly assigned prior distributions to each tip on the molecular phylogeny (**Table S6**).

First, we used stochastic character mapping (Bollback 2006; Revell 2012) to infer ancestral states at each node on the chronogram with vetted calibrations. For this we used a single dataset, where the natural history codings were simplified to marine, and marine/direct and freshwater/indirect transition paths (i.e., three unordered states of a single character), with the posterior distribution simulated 500 times. Next, we used Bayesian threshold models (Felsenstein 2012; Revell 2014) to account for gradients of change. A character coded with discrete ordered states (i.e., our grades) was assumed to evolve according to an unobserved continuous trait called “liability” (here representing the coded natural history traits combined with additional unobserved factors). Following Sallan et al. (2018), we assume thresholds represent the amount of change in terrestriality traits that allows a habitat shift. As there are two independent transition pathways with their own gradients, these were analyzed separately for each pathway from marine to non-marine (and on subclades for transitions between direct and indirect pathways). Threshold models for ancestral state reconstruction were implemented as the *ancThresh* function in *phytools* (Revell 2012). Each *ancThresh* model (Brownian Motion: BM, Ornstein-Uhlenbeck: OU, and Pagel’s Lambda) was run for 50 million generations (5 million for subclades) with 20% burnin (Revell 2014; Sallan et al. 2018).

#### References


Barido-Sottani J., Aguirre-Fernández G., Hopkins M.J., Stadler T., Warnock R.C. 2019. Ignoring stratigraphic age uncertainty leads to erroneous estimates of species divergence times under the fossilized birth-death process. Proceedings of the Royal Society B: Biological Sciences. 286:20190685.

Barido-Sottani J., Pohle A., De Baets K., Murdock D., Warnock R.C.M. 2022. Putting the F in FBD analyses: tree constraints or morphological data? BioRxiv.

Barido-Sottani J., van Tiel N., Hopkins M.J., Wright D.F., Stadler T., Warnock R.C.M. 2020. Ignoring fossil age uncertainty leads to inaccurate topology and divergence times in time calibrated tree inference. Frontiers in Ecology and Evolution. 8:183.

Bollback J.P. 2006. SIMMAP: stochastic character mapping of discrete traits on phylogenies. BMC Bioinformatics. 7:88.

Bouckaert R., Vaughan T.G., Barido-Sottani J., Duchêne S., Fourment M., Gavryushkina A., Heled J., Jones G., Kühnert D., De Maio N., Matschiner M., Mendes F.K., Müller N.F., Ogilvie H.A., du Plessis L., Popinga A., Rambaut A., Rasmussen D., Siveroni I., Suchard M.A., Wu C.-H., Xie D., Zhang C., Stadler T., Drummond A.J. 2019. BEAST 2.5: An advanced software platform for Bayesian evolutionary analysis. PLoS Comput Biol. 15:e1006650.

Bracken H.D., Toon A., Felder D.L., Martin J.W., Finley M., Rasmussen J., Palero F., Crandall K.A. 2009. The decapod tree of life: compiling the data and moving toward a consensus of decapod evolution. Arthropod Systematics & Phylogeny. 67:99–116.

Bracken-Grissom H.D., Cannon M.E., Cabezas P., Feldmann R.M., Schweitzer C.E., Ahyong S.T., Felder D.L., Lemaitre R., Crandall K.A. 2013. A comprehensive and integrative reconstruction of evolutionary history for Anomura (Crustacea: Decapoda). BMC Evolutionary Biology. 13:128.

Buhay J.E., Moni G., Mann N., Crandall K.A. 2007. Molecular taxonomy in the dark: Evolutionary history, phylogeography, and diversity of cave crayfish in the subgenus *Aviticambarus*, genus *Cambarus*. Molecular Phylogenetics and Evolution. 42:435–448.

Castresana J. 2000. Selection of conserved blocks from multiple alignments for their use in phylogenetic analysis. Molecular Biology and Evolution. 17:540–552.

Chernomor O., von Haeseler A., Minh B.Q. 2016. Terrace aware data structure for phylogenomic inference from supermatrices. Systematic Biology. 6:997–1008.

Colgan D.J., McLauchlan A., Wilson G.D.F., Livingston S.P., Edgecombe G.D., Macaranas J., Cassis G., Gray M.R. 1998. Histone H3 and U2 snRNA DNA sequences and arthropod molecular evolution. Aust. J. Zool. 46:419.

Crandall K.A., Fitzpatrick J.F. 1996. Crayfish molecular systematics: Using a combination of procedures to estimate phylogeny. Systematic Biology. 45:1–26.

Culshaw V., Stadler T., Sanmartín I. 2019. Exploring the power of Bayesian birth-death skyline models to detect mass extinction events from phylogenies with only extant taxa. Evolution. 73:1133–1150.

Drummond A.J., Ho S.Y.W., Phillips M.J., Rambaut A. 2006. Relaxed phylogenetics and dating with confidence. PLoS Biology. 4:e88.

Drummond A.J., Suchard M.A. 2010. Bayesian random local clocks, or one rate to rule them all. BMC Biol. 8:114.

Felsenstein J. 2012. A comparative method for both discrete and continuous characters using the threshold model. American Naturalist. 179:145-156.

Heath T.A., Huelsenbeck J.P., Stadler T. 2014. The fossilized birth-death process for coherent calibration of divergence-time estimates. Proceedings of the National Academy of Sciences. 111:E2957–E2966.

Hegna T.A., Luque J., Wolfe J.M. 2020. The fossil record of the Pancrustacea. In: Poore G.C.B., Thiel M., editors. Evolution and Biogeography. Oxford: Oxford University Press. p. 21–52.

Hoang D.T., Chernomor O., von Haeseler A., Minh B.Q., Vinh L.S. 2018. UFBoot2: Improving the ultrafast bootstrap approximation. Molecular Biology and Evolution. 35:518–522.

Kalyaanamoorthy S., Minh B.Q., Wong T.K.F., von Haeseler A., Jermiin L.S. 2017. ModelFinder: fast model selection for accurate phylogenetic estimates. Nature Methods. 14:587–589.

Katoh K., Standley D.M. 2013. MAFFT Multiple sequence alignment software version 7: Improvements in performance and usability. Molecular Biology and Evolution. 30:772–780.

Kearse M., Moir R., Wilson A., Stones-Havas S., Cheung M., Sturrock S., Buxton S., Cooper A., Markowitz S., Duran C., Thierer T., Ashton B., Meintjes P., Drummond A. 2012. Geneious Basic: An integrated and extendable desktop software platform for the organization and analysis of sequence data. Bioinformatics. 28:1647–1649.

López-Giráldez F., Townsend J.P. 2011. PhyDesign: an online application for profiling phylogenetic informativeness. BMC Evolutionary Biology. 11:152.
Luque, J., Bracken-Grissom H.D., Ortega-Hernández J., Wolfe J.M. 2023. Fossil calibrations for molecular analyses and divergence time estimation for true crabs (Decapoda: Brachyura). BioRxiv.
Medlin L., Elwood H.L., Stickel S., Sogin M.L. 1988. The characterization of enzymatically amplified eukaryotic 16S-like rRNA-coding regions. Gene. 71:491–499.
Minh B.Q., Hahn M.W., Lanfear R. 2020a. New methods to calculate concordance factors for phylogenomic datasets. Molecular Biology and Evolution. 37:2727–2733.
Minh B.Q., Schmidt H., Chernomor O., Schrempf D., Woodhams M., von Haeseler A., Lanfear R. 2020b. IQ-TREE 2: New models and efficient methods for phylogenetic inference in the genomic era. Molecular Biology and Evolution. 37:1530–1534.
Mo Y.K., Lanfear R., Hahn M.W., Minh B.Q. 2022. Updated site concordance factors minimize effects of homoplasy and taxon sampling. Bioinformatics.:1–2.
Mongiardino Koch N., Thompson J.R., Hatch A.S., McCowin M.F., Armstrong A.F., Coppard S.E., Aguilera F., Bronstein O., Kroh A., Mooi R., Rouse G.W. 2022. Phylogenomic analyses of echinoid diversification prompt a re-evaluation of their fossil record. eLife. 11:e72460.
Ng P.L.K., Guinot D., Davie P.J.F. 2008. Systema Brachyurorum: Part I. An annotated checklist of extant brachyuran crabs of the world. Raffles Bulletin of Zoology. 17:1–286.
O'Reilly J.E., Donoghue P.C.J. 2020. The effect of fossil sampling on the estimation of divergence times with the Fossilized Birth–Death process. Systematic Biology. 69:124–138.
Palero F., Guerao G., Abello P. 2008. Morphology of the final stage phyllosoma larva of *Scyllarus pygmaeus* (Crustacea: Decapoda: Scyllaridae), identified by DNA analysis. Journal of Plankton Research. 30:483–488.
Parham J.F., Donoghue P.C.J., Bell C.J., Calway T.D., Head J.J., Holroyd P.A., Inoue J.G., Irmis R.B., Joyce W.G., Ksepka D.T., Patane J.S.L., Smith N.D., Tarver J.E., van Tuinen M., Yang Z., Angielczyk K.D., Greenwood J.M., Hipsley C.A., Jacobs L., Makovicky P.J., Muller J., Smith K.T., Theodor J.M., Warnock R.C.M., Benton M.J. 2012. Best practices for justifying fossil calibrations. Systematic Biology. 61:346–359.
Poore G.C.B., Ahyong S.T. 2023. Marine decapod Crustacea: a guide to the families and genera of the world. CSIRO Publishing.
Rambaut A., Drummond A.J., Xie D., Baele G., Suchard M.A. 2018. Posterior summarisation in Bayesian phylogenetics using Tracer 1.7. Systematic Biology. 67:901–904.
Revell L.J. 2012. phytools: an R package for phylogenetic comparative biology (and other things). Methods in Ecology and Evolution. 3:217–223.
Revell L.J. 2014. Ancestral character estimation under the threshold model from quantitative genetics. Evolution. 68:743–759.
Ronquist F., Teslenko M., van der Mark P., Ayres D.L., Darling A., Hohna S., Larget B., Liu L., Suchard M.A., Huelsenbeck J.P. 2012. MrBayes 3.2: Efficient Bayesian phylogenetic inference and model choice across a large model space. Systematic Biology. 61:539–542.
Sallan L., Friedman M., Sansom R.S., Bird C.M., Sansom I.J. 2018. The nearshore cradle of early vertebrate diversification. Science. 362:460–464.
Schubart C.D., Cuesta J.A., Felder D.L. 2002. Glyptograpsidae, a new brachyuran family from Central America: Larval and adult morphology, and a molecular phylogeny of the Grapsoidea. Journal of Crustacean Biology. 22:28–44.
Schubart C.D., Neigel J.E., Felder D.L. 2000. The use of the mitochondrial 16S rDNA gene for phylogenetics and biogeographic studies of Crustacea. Crustacean Issues: The Biodiversity Crisis and Crustacea. Boca Raton: CRC Press. p. 817–830.
Simon C., Frati F., Beckenbach A., Crespi B., Liu H., Flook P. 1994. Evolution, weighting, and phylogenetic utility of mitochondrial gene sequences and a compilation of conserved Polymerase Chain Reaction primers. Annals of the Entomological Society of America. 87:651–701.
Song H., Buhay J.E., Whiting M.F., Crandall K.A. 2008. Many species in one: DNA barcoding overestimates the number of species when nuclear mitochondrial pseudogenes are coamplified. Proc. Natl. Acad. Sci. U.S.A. 105:13486–13491.
Spears T., Abele L.G. 1998. Crustacean phylogeny inferred from 18S rDNA. In: Fortey R.A., Thomas R.H., editors. Arthropod Relationships. Dordrecht: Springer. p. 169–187.
Stadler T. 2010. Sampling-through-time in birth–death trees. Journal of Theoretical Biology. 267:396–404.

Stadler T., Kuhnert D., Bonhoeffer S., Drummond A.J. 2013. Birth-death skyline plot reveals temporal changes of epidemic spread in HIV and hepatitis C virus (HCV). Proceedings of the National Academy of Sciences. 110:228–233.

Townsend J.P. 2007. Profiling phylogenetic informativeness. Systematic Biology. 56:222–231.

Tsang L.M., Chan T.-Y., Ahyong S.T., Chu K.H. 2011. Hermit to king, or hermit to all: Multiple transitions to crab-like forms from hermit crab ancestors. Systematic Biology. 60:616–629.

Tsang L.M., Ma K.Y., Ahyong S.T., Chan T.-Y., Chu K.H. 2008. Phylogeny of Decapoda using two nuclear protein-coding genes: Origin and evolution of the Reptantia. Molecular Phylogenetics and Evolution. 48:359–368.

Tsang L.M., Schubart C.D., Ahyong S.T., Lai J.C.Y., Au E.Y.C., Chan T.-Y., Ng P.K.L., Chu K.H. 2014. Evolutionary history of true crabs (Crustacea: Decapoda: Brachyura) and the origin of freshwater crabs. Molecular Biology and Evolution. 31:1173–1187.

Watson-Zink V.M. 2021. Making the grade: Physiological adaptations to terrestrial environments in decapod crabs. Arthropod Structure & Development. 64:101089.

Whiting M.F. 2002. Mecoptera is paraphyletic: multiple genes and phylogeny of Mecoptera and Siphonaptera. Zoologica Scripta. 31:93–104.

Whiting M.F., Carpenter J.C., Wheeler Q.D., Wheeler W.C. 1997. The Strepsiptera problem: Phylogeny of the holometabolous insect orders inferred from 18S and 28S ribosomal DNA sequences and morphology. Systematic Biology. 46:1–68.

Wolfe J.M., Breinholt J.W., Crandall K.A., Lemmon A.R., Moriarty Lemmon E., Timm L.E., Siddall M.E., Bracken-Grissom H.D. 2019. A phylogenomic framework, evolutionary timeline, and genomic resources for comparative studies of decapod crustaceans. Proceedings of the Royal Society B: Biological Sciences. 286:20190079.

Wolfe J.M., Daley A.C., Legg D.A., Edgecombe G.D. 2016. Fossil calibrations for the arthropod Tree of Life. Earth-Science Reviews. 160:43–110.

Wolfe J.M., Luque J., Bracken-Grissom H.D. 2021. How to become a crab: Phenotypic constraints on a recurring body plan. BioEssays. 43:2100020.

WoRMS. 2021. Brachyura. Accessed at: https://www.marinespecies.org/aphia.php?p=taxdetails&id=106673 on 06-08-2021.

WoRMS. 2022. Brachyura. Accessed at: https://www.marinespecies.org/aphia.php?p=taxdetails&id=106673 on 12-08-2022.

Zhou X., Shen X.-X., Hittinger C.T., Rokas A. 2018. Evaluating fast Maximum Likelihood-based phylogenetic programs using empirical phylogenomic data sets. Molecular Biology and Evolution. 35:486–503.

#### Text S2. Natural History References

These are the references listed in Table S5.


Aagaard A., Styrishave B., Warman C.G., Depledge M.H. 2000. The use of cardiac monitoring in the assessment of mercury toxicity in the subtropical pebble crab *Gaetice depressus* (Brachyura: Grapsidae: Varuninae). Sci. Mar. 64:381–386.

Abele L.G. 1979. The community structure of coral-associated decapod crustaceans in variable environments. In: Livingston R.J., editor. Ecological Processes in Coastal and Marine Systems. Boston, MA: Springer US. p. 265–287.

Adamczewska A.M., Morris S. 2000. Locomotion, respiratory physiology, and energetics of amphibious and terrestrial crabs. Physiological and Biochemical Zoology. 73:706–725.

Agusto L.E., Fratini S., Jimenez P.J., Quadros A., Cannicci S. 2021. Structural characteristics of crab burrows in Hong Kong mangrove forests and their role in ecosystem engineering. Estuarine, Coastal and Shelf Science. 248:106973.

Al-Aidaroos A.M., Al-Haj A.E., Kumar A.A.J. 2014. Complete larval development of the Brachyuran crab (*Epixanthus frontalis* H. Milne Edwards, 1834) (Crustacea, Decapoda, Eriphioidea, Oziidae) under laboratory conditions. Zootaxa. 3815:29.

Arias-Pineda J.Y., Gómez D.A., Magalhães C. 2015a. Range extension of the genus *Fredilocarcinu*s Pretzmann, 1978 (Crustacea: Decapoda: Trichodactylidae) to Colombia. Checklist: The Journal of Biodiversity Data. 11:1586.

Augusto A., Greene L.J., Laure H.J., Mcnamara J.C. 2007. Adaptive shifts in osmoregulatory strategy and the invasion of freshwater by brachyuran crabs: evidence from *Dilocarcinus pagei* (Trichodactylidae). J. Exp. Zool. 307A:688–698.

Banerjee S., Ramakrishnan S.C. 2022. The first record of *Ozius tuberculosus* H. Milne Edwards (Crustacea: Decapoda: Brachyura) from the East Coast of Indian subcontinent. Research Square.

Barnes R.S.K. 1967. The osmotic behaviour of a number of grapsoid crabs with respect to their differential penetration of an estuarine system. Journal of Experimental Biology. 47:535–551.

Barnwell F.H. 1968. The role of rhythmic systems in the adaptation of fiddler crabs to the intertidal zone. Am Zool. 8:569–583.

Bello-González O.C., Wehrtmann I.S., Magalhães C. 2016. First confirmed report of a primary freshwater crab (Brachyura: Pseudothelphusidae) associated with bromeliads in the Neotropics. Journal of Crustacean Biology. 36:303–309.

Bennett D.B. 1995. Factors in the life history of the edible crab (*Cancer pagurus* L.) that influence modelling and management. ICES Mar. Sci. Symp., 199: 89-98.

Bianchini A., Gilles R. 1996. Toxicity and accumulation of mercury in three species of crabs with different osmoregulatory capacities. Bulletin of Environmental Contamination and Toxicology. 57:91–98.

Branch G.M., Grindley J.R. 1979. Ecology of southern African estuaries Part XI. Mngazana: a mangrove estuary in Transkei. South African Journal of Zoology. 14:149–170.

Bright D.B., Hogue C.L. 1972. A synopsis of the burrowing land crabs of the world and list of their arthropod symbionts and burrow associates. Contributions in Science. 220:1–58.

Brown S., Bert T. 1993. The effects of temperature and salinity on molting and survival of *Menippe adina* and *M. mercenaria* (Crustacea, Decapoda) postsettlement juveniles. Mar. Ecol. Prog. Ser. 99:41–49.

Brown S.D., Bert T.M., Tweedale W.A., Torres J.J., Lindberg W.J. 1992. The effects of temperature and salinity on survival and development of early life stage Florida stone crabs *Menippe mercenaria* (Say). Journal of Experimental Marine Biology and Ecology. 157:115–136.

Cannicci S., Ruwa R.K., Giuggioli M. 1998. Predatory activity and spatial strategies of *Epixanthus dentatus* (Decapoda: Oziidae), an ambush predator among the mangroves. Journal of Crustacean Biology. 18(1): 57-63.

Charmantier G., Charmantier-Daures M. 1991. Ontogeny of osmoregulation and salinity tolerance in *Cancer irroratus* ; elements of comparison with *C. borealis* (Crustacea, Decapoda). The Biological Bulletin. 180:125–134.

Chen T.-Y., Hwang G.-W., Mayfield A.B., Chen C.-P., Lin H.-J. 2019. The development of habitat suitability models for fiddler crabs residing in subtropical tidal flats. Ocean & Coastal Management. 182:104931.

Cieluch U., Anger K., Aujoulat F., Buchholz F., Charmantier-Daures M., Charmantie G. 2004. Ontogeny of osmoregulatory structures and functions in the green crab *Carcinus maenas* (Crustacea, Decapoda). Journal of Experimental Biology. 207:325–336.
Clark P.F., Paula J. 2003. Descriptions of ten xanthoidean (Crustacea: Decapoda: Brachyura) first stage zoeas from Inhaca island, Mozambique. Raffles Bulletin of Zoology. 51:323-378.
Cook B.D., Pringle C.M., Hughes J.M. 2008. Phylogeography of an island endemic, the Puerto Rican Freshwater Crab (*Epilobocera sinuatifrons*). Journal of Heredity. 99:157–164.
Cuesta J.A., Guerao G., Liu H.-C., Schubart C.D. 2006. Morphology of the first zoeal stages of eleven Sesarmidae (Crustacea, Brachyura, Thoracotremata) from the Indo-West Pacific, with a summary of familial larval characters. Invertebrate Reproduction & Development. 49:151–173.
Cumberlidge N. 1991. The Respiratory System of *Globonautes macropus* (Rathbun, 1898), a terrestrial freshwater crab from Liberia (Gecarcinucoidea, Gecarcinucidae). Crustac. 61:69–80.
Cumberlidge N. 2008. Insular species of Afrotropical freshwater crabs (Crustacea: Decapoda: Brachyura: Potamonautidae and Potamidae) with special reference to Madagascar and the Seychelles. Contributions to Zoology. 77:71–81.
Cumberlidge N. 2009. The status and distribution of freshwater crabs [West Africa]. In: *The Status and Distribution of Freshwater Biodiversity in Western Africa*. Compilers: Smith, K.G., Diop, M.D., Niane, M. and Darwall, W.R.T. IUCN & Wetlands International report. Gland, Switzerland and Cambridge, UK: pp56-72
Curtis D.L., McGaw I.J. 2008. A year in the life of a Dungeness crab: methodology for determining microhabitat conditions experienced by large decapod crustaceans in estuaries. Journal of Zoology. 274:375–385.
Davie P.J.F., Guinot D., Ng P.K.L. 2015. Anatomy and functional morphology of Brachyura. Crustacea 9C (71-2): 11-163.
Kunsook C., Dumrongrojwatthana P. 2017. Species diversity and abundance of marine crabs (Portunidae: Decapoda) from a collapsible crab trap fishery at Kung Krabaen Bay, Chanthaburi Province, Thailand. Tropical Life Sciences Research. 28:45–67.
Depledge M.H. 1989. Observations on the feeding behaviour of *Gaetice depressus* (Grapsidae: Varuninae) with special reference to suspension feeding. Marine Biology. 100:253–259.
Diamond D.W., Scott L.K., Forward R.B., Kirby-Smith W. 1989. Respiration and osmoregulation of the estuarine crab, *Rhithropanopeus harrisii* (Gould): Effects of the herbicide, Alachlor. Comparative Biochemistry and Physiology Part A: Physiology. 93:313–318.
Dornelas M., Paula J., Macia A. 2003. The larval development of *Hymenosoma orbiculare* Desmarest, 1825 (Crustacea: Decapoda: Brachyura: Hymenosomatidae). Journal of Natural History. 37:2579–2597.
Edkins M.T., Teske P.R., Papadopoulos I., Griffiths C.L. 2007. Morphological and genetic analyses suggest that southern African crown crabs, *Hymenosoma orbiculare*, represent five distinct species. Crustaceana. 80:667–683.
Edwards M. 1986. Bimodal respiration, water balance and acid-base regulation in a high-shore crab, *Cyclograpsus lavauxi* H. Milne Edwards. J. Exp. Mar. Bio. Ecol., Vol. 100, pp. 127 - 145.
Erkan M., Balkis H., Tunali Y., Oliveria E. 2009. Morphology of testis and vas deferens in the xanthoid crab, *Eriphia verrucosa* (Forskål, 1775) (Decapoda: Brachyura). Journal of Crustacean Biology. 29:458–465.
Eshky A., Atkinson R., Taylor A. 1995. Physiological ecology of crabs from Saudi Arabian mangrove. Mar. Ecol. Prog. Ser. 126:83–95.
Esser L.J., Cumberlidge N. 2011. Evidence that salt water may not be a barrier to the dispersal of Asian freshwater crabs (Decapoda: Brachyura: Gecarcinucidae and Potamidae). Raffles Bulletin of Zoology, 59(2): 259-268.
Fadlaoui S., Melhaoui M. 2019. Population structure of the freshwater crab *Potamon algeriense* (Bott, 1967) inhabiting Oued Zegzel, (Northeast of Morocco). International Journal of Ecology. 2019:1–7.
Felder D.L., Martin J.W., Goy J.W. 2017. Patterns in early postlarval development of decapods. In: Schram F.R., Wenner A.M., editors. Crustacean Issues 2. Routledge. p. 163–225.
Fernandes L.S., Sant'Anna B.S., Hattori G.Y. 2020. Amazon freshwater crab *Dilocarcinus pagei* (Decapoda: Trichodactylidae): a view about burrow construction behavior. Lat. Am. J. Aquat. Res. 48:85–93.
Figueiredo J., Penha-Lopes G., Anto J., Narciso L., Lin J. 2008. Fecundity, brood loss and egg development through embryogenesis of *Armases cinereum* (Decapoda: Grapsidae). Mar Biol. 154:287–294.
Farhadi A., Harlioglu M.M. 2018. Distribution and diversity of freshwater crabs (Decapoda: Brachyura: Potamidae, Gecarcinucidae) in Iranian inland waters. Aquatic Sciences and Engineering:110–116.
Felder D.L., Rabalais N.N. 1986. The Genera *Chasmocarcinus* Rathbun and *Speocarcinus* Stimpson on the Continental Shelf of the Gulf of Mexico, with Descriptions of Two New Species (Decapoda: Brachyura: Goneplacidae). Journal Of Crustacean Biology. 6:30.

Foale S. 1999. Local Ecological Knowledge and Biology of the Land Crab *Cardisoma hirtipes* (Decapoda: Gecarcinidae) at West Nggela, Solomon Islands. Pacific Science. 53:13.
Forward R.B. 2009. Larval Biology of the Crab *Rhithropanopeus harrisii* (Gould): A Synthesis. The Biological Bulletin. 216:243–256.
Fowler A., Forsström T., von Numers M., Vesakoski O. 2013. The North American mud crab *Rhithropanopeus harrisii* (Gould, 1841) in newly colonized Northern Baltic Sea: distribution and ecology. Aquatic Invasions. 8:89–96.
Rodriguez G., Ng P.KL. 1995. Freshwater Crabs as Poor Zoogeographical Indicators: A Critique of Bănărescu (1990). Crustaceana. 68:636-645.
Fukui Y. 1988. Comparative Studies on the Life History of the Grapsid Crabs (Crustacea, Brachyura) Inhabiting Intertidal Cobble and Boulder Shores. Publ. SMBL. 33:121–162.
Full R.J., II C.F.H., Assad J.A. 1985. Energetics of the Exercising Wharf Crab *Sesarma cinereum*. Physiological Zoology. 58:605–615.
Galil B., Vannini M. 1990. Research on the coast of Somalia. Xanthidae Trapeziidae Carpiliidae Menippidae (Crustacea Brachyura). Tropical Zoology. 3:21–56.
Giraldes B.W., Chatting M., Smyth D. 2019. The fishing behaviour of *Metopograpsus messor* (Decapoda: Grapsidae) and the use of pneumatophore-borne vibrations for prey-localizing in an arid mangrove setting. J. Mar. Biol. Ass. 99:1353–1361.
Goodman S.H., Cavey M.J. 1990. Organization of a phyllobranchiate gill from the green shore crab *Carcinus maenas* (Crustacea, Decapoda). Cell Tissue Res. 260:495–505.
Guerao G. 2004. Complete larval and early juvenile development of the mangrove crab *Perisesarma fasciatum* (Crustacea: Brachyura: Sesarmidae) from Singapore, with a larval comparison of *Parasesarma* and *Perisesarma*. Journal of Plankton Research. 26:1389–1408.
Guinot D. 2007. A new genus of the family Plagusiidae Dana, 1851, close to *Plagusia* Latreille, 1804 (Crustacea, Decapoda, Brachyura). Zootaxa. 1498:27–33.
Hartnoll R. 2009. Sexual Maturity and Reproductive Strategy of the Rock Crab *Grapsus adscensionis* (Osbeck, 1765) (Brachyura, Grapsidae) on Ascension Island. Crustac. 82:275–291.
Hartnoll R.G. 1976. The ecology of some rocky shores in tropical East Africa. Estuarine and Coastal Marine Science. 4:1–21.
Hegerl E.J., Davie J.D.S. 1970. The mangrove forests of Cairns, Northern Australia. Mar. Res. Indonesia. 18:23–57.
Herreid C.F., Gifford C.A. 1963. The Burrow Habitat of the Land Crab, *Cardisoma guanhumi* (Latreille). Ecology. 44:773–775.
Hill B.J., Forbes A.T. 1979. Biology of *Hymenosoma orbiculare* Desm in Lake Sibaya. South African Journal of Zoology. 14:75–79.
Holsman K., McDonald P., Armstrong D. 2006. Intertidal migration and habitat use by subadult Dungeness crab *Cancer magister* in a NE Pacific estuary. Mar. Ecol. Prog. Ser. 308:183–195.
Hsueh P.-W. 2018. A new species of *Neorhynchoplax* (Crustacea: Decapoda: Brachyura: Hymenosomatidae) from Taiwan. Zootaxa. 4461:350.
Huang J.-F., Yang S.-L., Ng P.K.L. 1998. Notes On the Taxonomy and Distribution of Two Closely Related Species of Ghost Crabs, *Ocypode sinensis* and *O. cordimanus* (Decapoda, Brachyura, Ocypodidae). Crustac. 71:942–954.
Hunter K.C., Rudy P.P. 1975. Osmotic and ionic regulation in the Dungeness crab, *Cancer magister* Dana. Comparative Biochemistry and Physiology Part A: Physiology. 51:439–447.
Vázquez-López H., Vega-Villasante F., Rodríguez-Varela A.C., Cruz-Gómez A. 2014. Population density of the land crab *Cardisoma crassum* Smith, 1870 (Decapoda: Gecarcinidae) in the estuary El Salado, Puerto Vallarta, Jalisco, Mexico. International Journal of Innovative and Applied Research. 2:1-9.
IUCN. 2008a. *Ceylonthelphusa alpina*: Bahir, M.M., Ng, P.K.L., Crandall, K. & Pethiyagoda, R. and Cumberlidge, N.: The IUCN Red List of Threatened Species 2008: e.T61690A12525900. .
IUCN. 2008b. *Potamonautes perlatus*: Cumberlidge, N.: The IUCN Red List of Threatened Species 2008: e.T64389A12767481. .
IUCN. 2018. *Syntripsa flavichela*: Schubart, C.: The IUCN Red List of Threatened Species 2018: e.T134942A109683110.
Jones D.A. 1972. Aspects of the ecology and behaviour of *Ocypode ceratophthalmus* (Pallas) and *O. kuhlii* de Haan (Crustacea: Ocypodidae). Journal of Experimental Marine Biology and Ecology. 8:31-43.
Kennish R., Williams G. 1997. Feeding preferences of the herbivorous crab *Grapsus albolineatus*: The differential influence of algal nutrient content and morphology. Mar. Ecol. Prog. Ser. 147:87–95.

Kepel' A.A., Tsareva L.A. 2005. First Record of Tropical Crabs *Portunus sanguinolentus* (Herbst, 1783) and *Plagusia depressa tuberculata* Lamarck, 1818 in Peter the Great Bay, Sea of Japan. Russ J Mar Biol. 31:124–125.

Kim S.Y., Yi C.H., Kim J.M., Choi W.Y., Kim H.S., Kim M.-S. 2020. DNA Barcoding of the Marine Protected Species *Parasesarma bidens* (Decapoda: Sesarmidea) from the Korean Waters. Animal Systematics, Evolution and Diversity. 36:159–163.

Kitaura J., Wada K., Nishida M. 2002. Molecular phylogeny of grapsoid and ocypodoid crabs with special reference to the genera *Metaplax* and *Macrophthalmus*. Journal Of Crustacean Biology. 22:12.

Kobayashi S., Archdale M.V. 2022. Burrow utilization by the Japanese mitten crab *Eriocheir japonica* in the clay sediment of a freshwater river of Kyushu, Japan. Crust Res. 51:17–29.

Kobayashi S., Vazquez Archdale M. 2021. Habitat preference and locomotive traits of the Japanese mitten crab *Eriocheir japonica* within the stream unit of the freshwater river environment. Limnologica. 88:125870.

Krediet C.J., Donahue M.J. 2009. Growth-mortality trade-offs along a depth gradient in *Cancer borealis*. Journal of Experimental Marine Biology and Ecology. 373:133–139.

Krishnan T., Kannupandi T. 1988. Larval development of *Elamena (Trigonoplax) cimex* Kemp, 1915 in the laboratory: The most unusual larvae known in the Brachyura (Crustacea: Decapoda). Bulletin Of Marine Science. 43:14.

Krishnan T., Kannupandi T. 1989. Laboratory cultured zoeae and megalopa of the mangrove crab *Metaplax distincta* H.Milne Edwards, 1852 (Brachyura: Sesarminae). J Plankton Res. 11:633–648.

Le Vay L., Ut V.N., Walton M. 2007. Population ecology of the mud crab *Scylla paramamosain* (Estampador) in an estuarine mangrove system; a mark-recapture study. Mar Biol. 151:1127–1135.

Lee S.H. 2022. Morphology of First Zoea Stage of *Sphaerozius nitidus* (Decapoda: Eriphioidea: Menippidae) Reared in the Laboratory Material. Animal Systematics, Evolution and Diversity. 38:83–90.

Lewis L., Ayers J. 2014. Temperature preference and acclimation in the Jonah Crab, *Cancer borealis*. Journal of Experimental Marine Biology and Ecology. 455:7–13.

Li J.-J., Shih H.-T., Ng P.K.L. 2019. Three New Species and Two New Records of *Parasesarma* De Man, 1895 (Crustacea: Brachyura: Sesarmidae) from Taiwan and the Philippines from Morphological and Molecular Evidence. Zoological Studies.:23.

Liu H.-C., Jeng M.-S. 2007. Some Reproductive Aspects of *Gecarcoidea lalandii* (Brachyura: Gecarcinidae) in Taiwan. Zoological Studies.:8.

Lucas J.S. 1980. Spider crabs of the family Hymenosomatidae (Crustacea; Brachyura) with particular reference to Australian species: Systematics and biology. Rec. Aust. Mus. 33:148–247.

Lucas J.S. 1971. The larval stages of some Australian species of *Halicarcinus* (Crustacea, Brachyura, Hymenosomatidae). I. Morphology. Bulletin of Marine Science. 21:471-490.

Lucas J.S. 1972. The larval stages of some Australian species of *Halicarcinus* (Crustacea, Brachyura, Hymenosomatidae). II. Physiology. Bulletin of Marine Science. 25:824-840.

Madeira D., Narciso L., Cabral H.N., Diniz M.S., Vinagre C. 2012. Thermal tolerance of the crab *Pachygrapsus marmoratus*: intraspecific differences at a physiological (CTMax) and molecular level (Hsp70). Cell Stress and Chaperones. 17:707–716.

Magalhaes C. 1996. Taxonomy of the Neotropical freshwater crab family Trichodactylidae III. The genera *Fredilocarcinus* and *Goyazana*. Senckenbergiana biologica:12.

Magalhães C., Barbosa U.C., Py-Daniel V. 2006. Decapod crustaceans used as food by the Yanomami Indians of the Balawa-ú village, State of Amazonas, Brazil. Acta Amaz. 36:369–374.

Maitland D.P. 1990. Aerial respiration in the semaphore crab, *Heloecius cordiformis*, with or without branchial water. Comparative Biochemistry and Physiology Part A: Physiology. 95:267–274.

Marijnissen S.A.E., Michel E., Daniels S.R., Erpenbeck D., Menken S.B.J., Schram F.R. 2006. Molecular evidence for recent divergence of Lake Tanganyika endemic crabs (Decapoda: Platythelphusidae). Molecular Phylogenetics and Evolution. 40:628–634.

Matsumasa M., Kikuchi S., Takeda S., Poovachiranon S., Yong H.-S., Murai M. 2001. Blood Osmoregulation and Ultrastructure of the Gas Windows ('Tympana') of Intertidal Ocypodid Crabs: *Dotilla* vs. *Scopimera*. Benthos Research. 56:47–55.

McGaw I.J. 2005. Burying behaviour of two sympatric crab species: *Cancer magister* and *Cancer productus*. Sci. Mar. 69:375–381.

Melrose M.J. 1975. The Marine Fauna of New Zealand: Family Hymenosomatidae (Crustacea, Decapoda, Brachyura). :123.

Miculka B. 2009. Burrowing Habits, Habitat Selections, and Behaviors of Four Common Dominican Land Crabs; *Guinotia dentata*, *Gecarcinus lateralis*, *Gecarcinus ruricola*, and *Cardisoma guanhumi*. :16.
Molina Z.S., Neri J.B., Cañete R.M.P., Metillo E.B. 2020. Body Size, Habitat, and Diet of Freshwater Crabs *Isolapotamon mindanaoense* and *Sundathelphusa miguelito* (Crustacea: Brachyura) in the Municipality of Lake Sebu, South Cotabato, Philippines. Science Diliman:21.
Morris S. 2002. The ecophysiology of air-breathing in crabs with special reference to *Gecarcoidea natalis*. Comparative Biochemistry and Physiology Part B: Biochemistry and Molecular Biology. 131:559–570.
Murai M., Henmi Y., Matsumasa M., Backwell P.R.Y., Takeshita F. 2022. Attraction waves of male fiddler crabs: A visual display designed for efficacy. Journal of Experimental Marine Biology and Ecology. 546:151665.
Naderloo R., Fatemi Y. 2015. New record of the rare alien grapsoid crab *Percnon affine* (H. Milne Edwards, 1853) (Crustacea, Decapoda, Grapsoidea, Plagusidae) from the northern Indian Ocean. Mar. Biodivers. Rec. 8:e92.
Naderloo R., Türkay M. 2009. A new species of the genus *Nanosesarma* (Crustacea: Decapoda: Brachyura: Sesarmidae), and redescription of *Nanosesarma jousseaumei* (Nobili, 1906), including new records from the Persian Gulf. Journal of Natural History. 43:2911–2923.
Naderloo R., Türkay M., Sari A. 2013. Intertidal habitats and decapod (Crustacea) diversity of Qeshm Island, a biodiversity hotspot within the Persian Gulf. Mar Biodiv. 43:445–462.
Ng P.K.L., Dudgeon D. 1992. The Potmidae and Parathelphusidae (Crustacea: Decapoda: Brachyura) (Crustacea: Decapoda: Brachyura) of Hong Kong. Invertebrate Systematics:28.
Ng P.K.L. 1995. *Somanniathelphusa lacuvita*, a new ricefield crab from Tonle Sap, Cambodia. Asian Journal of Tropical Biology:6.
Ng P.K.L., Manuel-Santos M.R. 2007. Establishment of the Vultocinidae, a new family for an unusual new genus and new species of Indo-West Pacific crab (Crustacea: Decapoda: Brachyura: Goneplacoidea), with comments on the taxonomy of the Goneplacidae. Zootaxa. 1558:39–68.
Ng P.K.L., Ng P.Y.C. 2021. Land crabs (Crustacea: Brachyura: Gecarcinidae) of Singapore. Nature in Singapore. 14:113.
Niu J., Hu X.L., Ip J.C.H., Ma K.Y., Tang Y., Wang Y., Qin J., Qiu J.-W., Chan T.F., Chu K.H. 2020. Multi-omic approach provides insights into osmoregulation and osmoconformation of the crab *Scylla paramamosain*. Sci Rep. 10:21771.
Novak M. 2004. Diurnal activity in a group of Gulf of Maine decapods. Crustac. 77:603–620.
Nye P.A. 1974. Burrowing and burying by the crab *Macrophthalmus hirtipes*. New Zealand Journal of Marine and Freshwater Research. 8:243–254.
de Oliveira Rocha C.A., de Lira J.J.P.R., de Lemos Santana J., Guimarães M.P., dos Santos Calado T.C. 2019. Biological aspects of the marine crab *Plagusia depressa* (Fabricius, 1775) on the northeast coast of Brazil. Marine Biology Research. 15:181–190.
Padate V.P., Vijaylaxmi J., Maurya S., Rivonker C.U. 2015. Morphology and morphometrics of *Dotilla myctiroides* (Decapoda: Brachyura: Dotillidae) from an impacted beach of Goa along west coast of India. 57:7.
Padghane S., Chavan S. Habitat preference and maintenance of freshwater crab *Barytelphusa cunicularis* in concrete tank culture model. Journal of Entomology and Zoology Studies.:7.
Paoli F., Wirkner C.S., Cannicci S. 2015. The branchiostegal lung of *Uca vocans* (Decapoda: Ocypodidae): Unreported complexity revealed by corrosion casting and MicroCT techniques. Arthropod Structure & Development. 44:622–629.
Papadopoulos I., Newman B.K., Schoeman D.S., Wooldridge T.H. 2006. Influence of salinity and temperature on the larval development of the crown crab, *Hymenosoma orbiculare* (Crustacea: Brachyura: Hymenosomatidae). African Journal of Aquatic Science. 31:43–52.
Pati S.K., Devi A.R.S. 2015. *Spiralothelphusa gibberosa*, a new freshwater crab (Brachyura: Gecarcinucidae) from Thrissur district, Kerala, India. Zootaxa. 3963:416.
Pati S.K., Sharma R.M. 2014. Description of *Ghatiana*, a new genus of freshwater crab, with two new species and a new species of *Gubernatoriana* (Crustacea: Decapoda: Brachyura: Gecarcinucidae) from the Western Ghat Mountains, India. Journal of Natural History. 48:1279–1298.
Paula J. 2004. Patterns of temporal occurrence of brachyuran crab larvae at Saco mangrove creek, Inhaca Island (South Mozambique): implications for flux and recruitment. Journal of Plankton Research. 26:1163–1174.
Paula J., Hartnoll R.G. 1989. The larval and post larval development of *Percnon gibbesi* (Crustacea, Brachyura, Grapsidae) and the identity of the larval genus *Pluteocaris*. Journal of Zoology. 218:17–37.
Peyrot-Clausade M. 1989. Crab cryptofauna (Brachyura and Anomura) of Tikehau, Tuamotu Archipelago, French Polynesia. Coral Reefs. 8:109–117.

Pinder A.W., Smits A.W. 1993. The Burrow Microhabitat of the Land Crab *Cardisoma guanhumi*: Respiratory/Ionic Conditions and Physiological Responses of Crabs to Hypercapnia. Physiological Zoology. 66:216–236.
Qhaji Y., van Vuuren B.J., Papadopoulos I., McQuaid C., Teske P. 2015. A comparison of genetic structure in two low-dispersal crabs from the Wild Coast, South Africa. African Journal of Marine Science. 37:345–351.
Rasmuson L.K. 2013. The Biology, Ecology and Fishery of the Dungeness crab, *Cancer magister*. Advances in Marine Biology. Elsevier. p. 95–148.
Rodríguez G., Rodriguez R. 1992. The freshwater crabs of America. Family Trichodactylidae and supplement to the family Pseudothelphusidae. Paris: Orstom.
Rolier Lara L., Wehrtmann I.S., Magalhaes C., Mantelatto F.L. 2013. Species diversity and distribution of freshwater crabs (Decapoda: Pseudothelphusidae) inhabiting the basin of the Rio Grande de Terraba, Pacific slope of Costa Rica. lajar. 41:685–695.
Rosa L.C. da, Santos R.B. dos, Freire K.M.F. 2018. First record of the cliff crab *Plagusia depressa* (Fabricius, 1775) (Decapoda: Plagusiidae) from the coast of Sergipe, Brazil. Nauplius. 26:e2018030.
Rosenberg M.S. 2020. A fresh look at the biodiversity lexicon for fiddler crabs (Decapoda: Brachyura: Ocypodidae). Part 2: Biogeography. Journal of Crustacean Biology. 40:364–383.
Sastry A.N. 1977. The Larval Development of the Jonah Crab, *Cancer borealis* Stimpson, 1859, Under Laboratory Conditions (Decapoda Brachyura). Crustac. 32:290–303.
Schlotterbeck R.E. 1976. The Larval Development of the Lined Shore Crab, *Pachygrapsus crassipes* Randall, 1840 (Decapoda Brachyura, Grapsidae) Reared in the Laboratory. Crustac. 30:184–200.
Schubart C.D., Cuesta J.A., Felder D.L. 2002. Glyptograpsidae, A New Brachyuran Family From Central America: Larval And Adult Morphology, And A Molecular Phylogeny Of The Grapsoidea. Journal Of Crustacean Biology. 22:17.
Senkman L.E., Negro C.L., Lopretto E.C., Collins P.A. 2015. Reproductive behaviour of three species of freshwater crabs of the family Trichodactylidae (Crustacea: Decapoda) including forced copulation by males. Marine and Freshwater Behaviour and Physiology. 48:77–88.
Shih H.-T., Fang S.-H., Ng P.K.L. 2007. Phylogeny of the freshwater crab genus *Somanniathelphusa* Bott (Decapoda:Parathelphusidae) from Taiwan and the coastal regions of China, with notes on their biogeography. Invert. Systematics. 21:29.
Shih H.-T., Hsu J.-W., Wong K.J.H., Ng N.K. 2019. Review of the mudflat varunid crab genus *Metaplax* (Crustacea, Brachyura, Varunidae) from East Asia and northern Vietnam. ZK. 877:1–29.
Spencer A.M., Fielding A.H., Kamemoto F.I. 1979. The Relationship between Gill NaK-ATPase Activity and Osmoregulatory Capacity in Various Crabs. Physiological Zoology. 52:1–10.
Stanton D.J., Leven M.R., Hui T.C.H. 2017. Distribution of *Cryptopotamon anacoluthon* (Kemp, 1918) (Crustacea: Brachyura: Potamidae), a freshwater crab endemic to Hong Kong. J. Threat. Taxa. 9:9786.
Staton J.L., Sulkin S.D. 1991. Nutritional requirements and starvation resistance in larvae of the brachyuran crabs *Sesarma cinereum* (Bose) and *S. reticulatum* (Say). Journal of Experimental Marine Biology and Ecology:14.
Stutz A.M. 1978. Tidal and diurnal activity rhythms in the striped shore crab *Pachygrapsus crassipes*. Journal of Interdisciplinary Cycle Research. 9:41–48.
Sudha K., Anilkumar G. 1996. Seasonal growth and reproduction in a highly fecund brachyuran crab, *Metopograpsus messor* (Forskal) (Grapsidae). Hydrobiologia. 319:15–21.
Sukmaningrum T., Adji B.K., Pratiwi E.M., Larasati B., Sayekti P.R., Maulana I., Eprilurahman R. 2018. Diversity of crabs in the intertidal zone at Sundak Beach, Gunungkidul, Indonesia. :020066.
Sutton P.G. 2009. The occurrence of the freshwater crab *Potamon fluviatile* (Decapoda: Brachyura) in Corfu. Mar. Biodivers. Rec. 2:e58.
Takeda M., Miyake S. 1969. Crabs from the East China Sea. II : Addition to Brachygnatha Brachyrhyncha. Journal of the Faculty of Agriculture, Kyushu University. 15:449–468.
Takeda S., Matsumasa M., Kikuchi S., Poovachiranon S., Murai M. 1996. Variation in the Branchial Formula of Semiterrestrial Crabs (Decapoda: Brachyura: Grapsidae and Ocypodidae) in Relation to Physiological Adaptations to the Environment. Journal of Crustacean Biology. 16:472.
Takeda S., Murai M. 2004. Microhabitat Use by the Soldier Crab *Mictyris brevidactylus* (Brachyura: Mictyridae): Interchangeability of Surface and Subsurface Feeding through Burrow Structure Alteration. Journal of Crustacean Biology. 24:327–339.
Thrupp T.J., Pope E.C., Whitten M.M.A., Bull J.C., Wootton E.C., Edwards M., Vogan C.L., Rowley A.F. 2015. Disease profiles of juvenile edible crabs (*Cancer pagurus* L.) differ at two geographically-close intertidal sites. Journal of Invertebrate Pathology. 128:1–5.

Tiralongo F., Messina G., Lombardo B.M. 2021. Invasive Species Control: Predation on the Alien Crab *Percnon gibbesi* (H. Milne Edwards, 1853) (Malacostraca: Percnidae) by the Rock Goby, *Gobius paganellus* Linnaeus, 1758 (Actinopterygii: Gobiidae). JMSE. 9:393.

Tomikawa N., Watanabe S. 1992. Reproductive Ecology of the Xanthid Crab *Eriphia smithii* Mcleay. Journal of Crustacean Biology. 12:57–67.

Trivedi J.N., Gadhavi M.K., Vachhrajani K.D. 2012. Diversity and habitat preference of brachyuran crabs in Gulf of Kutch, Gujarat, India. :11.

Tsai J.-R., Lin H.-C. 2012. A Shift to an Ion Regulatory Role by Gills of a Semi-Terrestrial Crab. Zoological Studies.:13.

Tseng K.-Y., Tsai J.-R., Lin H.-C. 2022. A Multi-Species Comparison and Evolutionary Perspectives on Ion Regulation in the Antennal Gland of Brachyurans. Front. Physiol. 13:902937.

Vannini M., Chelazzi G., Gherardi F. 1989. Feeding habits of the pebble crab *Eriphia smithi* (Crustacea, Brachyura, Menippidae). Marine Biology. 100:249–252.

Vannini M., Gherardi F. 1988. Studies on the pebble crab, *Eriphia smithi* MacLeay 1838 (Xanthoidea Menippidae): patterns of relative growth and population structure. Tropical Zoology. 1:203–216.

Venkatachari S.A.T., Vasantha N. 1981. Salinity tolerance and osmoregulation in the freshwater crab, *Barytelphusa guerini* (Milne Edwards). Hydrobiologia. 77:133–138.

Vermeiren P., Abrantes K., Sheaves M. 2015. Generalist and Specialist Feeding Crabs Maintain Discrete Trophic Niches Within and Among Estuarine Locations. Estuaries and Coasts. 38:2070–2082.

Vijaylaxmi J., Can A.A., Rivonker C.U. 2020. Population biology of some species of crabs along Goa, west coast of India. 49:8.

Wada K., Kuroda M., Kamada M. 2005. Factors Influencing Coexistence of Two Brachyuran Crabs, *Helice tridens* and *Parasesarma plicatum*, in an Estuarine Salt Marsh, Japan. Journal of Crustacean Biology. 25:146–153.

Wahyudi A.J., Wada S., Aoki M., Hama T. 2015. *Gaetice depressus* (Crustacea, Varunidae): Species profile and its role in organic carbon and nitrogen flow. Ocean Sci. J. 50:389–401.

Waiho K., Fazhan H., Quinitio E.T., Baylon J.C., Fujaya Y., Azmie G., Wu Q., Shi X., Ikhwanuddin M., Ma H. 2018. Larval rearing of mud crab (*Scylla*): What lies ahead. Aquaculture. 493:37–50.

Walker J.B., Rinehart S.A., White W.K., Grosholz E.D., Long J.D. 2021. Local and regional variation in effects of burrowing crabs on plant community structure. Ecology. 102.

Walton M.E., Le Vay L., Truong L.M., Ut V.N. 2006. Significance of mangrove–mudflat boundaries as nursery grounds for the mud crab, *Scylla paramamosain*. Mar Biol. 149:1199–1207.

Wang Z., Bai Y., Zhang D., Tang B. 2018a. Adaptive evolution of osmoregulatory-related genes provides insight into salinity adaptation in Chinese mitten crab, *Eriocheir sinensis*. Genetica. 146:303–311.

Wang Z., Wang Z., Shi X., Wu Q., Tao Y., Guo H., Ji C., Bai Y. 2018b. Complete mitochondrial genome of *Parasesarma affine* (Brachyura: Sesarmidae): Gene rearrangements in Sesarmidae and phylogenetic analysis of the Brachyura. International Journal of Biological Macromolecules. 118:31–40.

Warren J. 1990. Role of burrows as refuges from sub-tidal predators of temperate mangrove crabs. Mar. Ecol. Prog. Ser. 67:295–299.

Wear R.G., Fielder D.R. 1985. The marine fauna of New Zealand: larvae of the Brachyura (Crustacea, Decapoda). Wellington: New Zealand Dept. of Scientific and Industrial Research.

Wehrtmann I.S., Magalhães C., Hernáez P., Mantelatto F.L. 2010. Offspring production in three freshwater crab species (Brachyura: Pseudothelphusidae) from the Amazon region and Central America. Zoologia (Curitiba, Impr.). 27:965–972.

Wehrtmann I.S., Magalhães C., Orozco M.N. 2016. The primary freshwater crabs of Guatemala (Decapoda: Brachyura: Pseudothelphusidae), with comments on their conservation status. Journal of Crustacean Biology. 36:776–784.

Weiss M., Thatje S., Heilmayer O., Anger K., Brey T., Keller M. 2009. Influence of temperature on the larval development of the edible crab, *Cancer pagurus*. J. Mar. Biol. Ass. 89:753–759.

Wilson B.M., Pohle G.W. 2016. Northern range expansion of the American talon crab, *Euchirograpsus americanus* A. Milne-Edwards, 1880 (Decapoda, Grapsoidea, Plagusiidae), to the Bay of Fundy, Canada. Crustac. 89:163–173.

Woll A. 2003. In situ observations of ovigerous *Cancer pagurus* Linnaeus, 1758 in Norwegian waters (Brachyura, Cancridae). Crustac. 76:469–478.

Wong K.J.H., Tao L.S.R., Leung K.M.Y. 2021. Subtidal crabs of Hong Kong: Brachyura (Crustacea: Decapoda) from benthic trawl surveys conducted by the University of Hong Kong, 2012 to 2018. Regional Studies in Marine Science. 48:102013.

Wortham J.L., Pascual S. 2017. Grooming behaviors and gill fouling in the commercially important blue crab (*Callinectes sapidus*) and stone crab (*Menippe mercenaria*). Nauplius. 25.

Yeo D.C.J., Cai Y., Ng P.K.L. 1999. The freshwater and terrestrial decapod crustacea of Pulau Tioman, Peninsular Malaysia. Raffles Bulletin of Zoology:48.

Yeo D.C.J., Ng P.K.L., Cumberlidge N., Magalhães C., Daniels S.R., Campos M.R. 2008. Global diversity of crabs (Crustacea: Decapoda: Brachyura) in freshwater. Hydrobiologia. 595:275–286.

Young A.M., Elliott J.A. 2020. Life History and Population Dynamics of Green Crabs (*Carcinus maenas*). Fishes. 5:44.

Zalota A.K., Spiridonov V.A., Kolyuchkina G.A. 2016. In situ observations and census of invasive mud crab *Rhithropanopeus harrisii* (Crustacea: Decapoda: Panopeidae) applied in the Black Sea and the Sea of Azov. Arthropoda Selecta. 25:39062–0.

Zenone A., Badalamenti F., Giacalone V.M., Musco L., Pipitone C., Vega Fernández T., D'Anna G. 2017. Substrate preference and settlement behaviour of the megalopa of the invasive crab *Percnon gibbesi* (Decapoda, Percnidae) in the Mediterranean Sea. Helgol Mar Res. 70:21.

Zhang D., Liu J., Qi T., Ge B., Liu Q., Jiang S., Zhang H., Wang Z., Ding G., Tang B. 2018. Comparative transcriptome analysis of *Eriocheir japonica sinensis* response to environmental salinity. PLoS ONE. 13:e0203280.

Zimmermann B.L., Aued A.W., Machado S., Manfio D., Scarton L.P., Santos S. 2009. Behavioral repertory of *Trichodactylus panoplus* (Crustacea: Trichodactylidae) under laboratory conditions. Zoologia (Curitiba, Impr.). 26:5–11.

Fig. S1

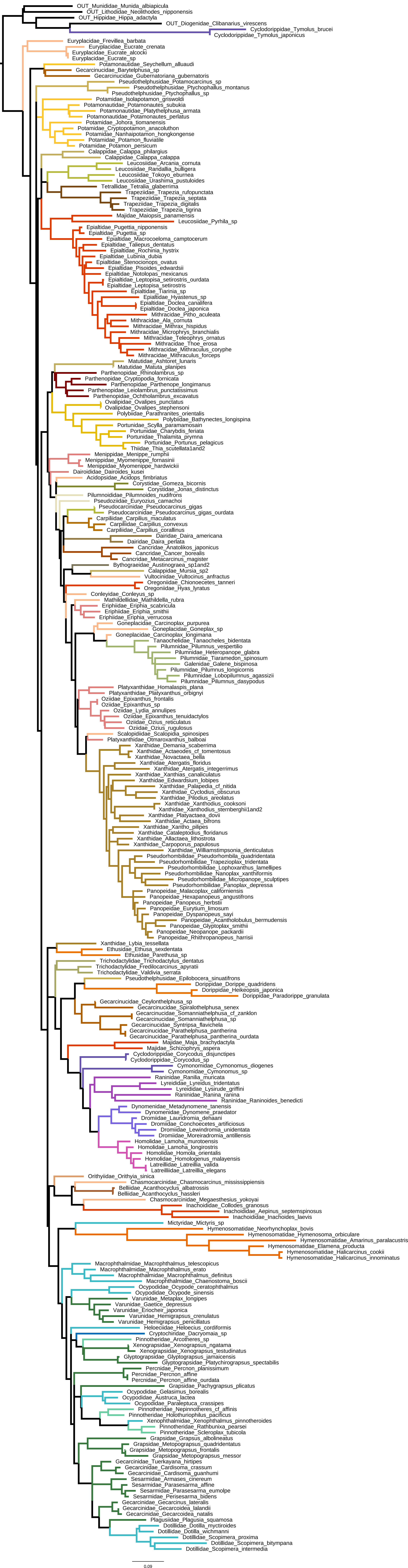
OUT_Munididae_Munida_albiapicula
OUT_Lithodidae_Neolithodes_nipponensis
OUT_Hippidae_Hippa_adactyla
OUT_Diogenidae_Clibanarius_virescens
Cyclodorippidae_Tymolus_brucei
Cyclodorippidae_Tymolus_japonicus
Euryplacidae_Frevillea_barbata
Euryplacidae_Eucrate_crenata
Euryplacidae_Eucrate_alcocki
Euryplacidae_Eucrate_sp
Potamonautidae_Seychellum_alluaudi
Gecarcinucidae_Barytelphusa_sp
Gecarcinucidae_Gubernatoriana_gubernatoris
Pseudothelphusidae_Potamocarcinus_sp
Pseudothelphusidae_Ptychophallus_montanus
Pseudothelphusidae_Ptychophallus_sp
Potamidae_Isolapotamon_griswoldi
Potamonautidae_Potamonautes_subukia
Potamonautidae_Platythelphusa_armata
Potamonautidae_Potamonautes_perlatus
Potamidae_Johora_tiomanensis
Potamidae_Cryptopotamon_anacoluthon
Potamidae_Nanhaipotamon_hongkongense
Potamidae_Potamon_fluviatile
Potamidae_Potamon_persicum
Calappidae_Calappa_philargius
Calappidae_Calappa_calappa
Leucosiidae_Arcania_cornuta
Leucosiidae_Randallia_bulligera
Leucosiidae_Tokoyo_eburnea
Leucosiidae_Urashima_pustuloides
Tetrallidae_Tetralia_glaberrima
Trapeziidae_Trapezia_rufopunctata
Trapeziidae_Trapezia_septata
Trapeziidae_Trapezia_digitalis
Trapeziidae_Trapezia_tigrina
Majidae_Maiopsis_panamensis
Leucosiidae_Pyrhila_sp
Epialtidae_Pugettia_nipponensis
Epialtidae_Pugettia_sp
Epialtidae_Macrocoeloma_camptocerum
Epialtidae_Taliepus_dentatus
Epialtidae_Rochinia_hystrix
Epialtidae_Lubinia_dubia
Epialtidae_Stenocionops_ovatus
Epialtidae_Pisoides_edwardsii
Epialtidae_Notolopas_mexicanus
Epialtidae_Leptopisa_setirostris_ourdata
Epialtidae_Leptopisa_setirostris
Epialtidae_Tiarinia_sp
Epialtidae_Hyastenus_sp
Epialtidae_Doclea_canalifera
Epialtidae_Doclea_japonica
Mithracidae_Pitho_aculeata
Mithracidae_Ala_cornuta
Mithracidae_Mithrax_hispidus
Mithracidae_Microphrys_branchialis
Mithracidae_Teleophrys_ornatus
Mithracidae_Thoe_erosa
Mithracidae_Mithraculus_coryphe
Mithracidae_Mithraculus_forceps
Matutidae_Ashtoret_lunaris
Matutidae_Matuta_planipes
Parthenopidae_Rhinolambrus_sp
Parthenopidae_Cryptopodia_fornicata
Parthenopidae_Parthenope_longimanus
Parthenopidae_Leiolambrus_punctatissimus
Parthenopidae_Ochtholambrus_excavatus
Ovalipidae_Ovalipes_punctatus
Ovalipidae_Ovalipes_stephensoni
Polybiidae_Parathranites_orientalis
Polybiidae_Bathynectes_longispina
Portunidae_Scylla_paramamosain
Portunidae_Charybdis_feriata
Portunidae_Thalamita_prymna
Portunidae_Portunus_pelagicus
Thiidae_Thia_scutellata1and2
Menippidae_Menippe_rumphii
Menippidae_Myomenippe_fornasinii
Menippidae_Myomenippe_hardwickii
Dairoididae_Dairoides_kusei
Acidopsidae_Acidops_fimbriatus
Corystidae_Gomeza_bicornis
Corystidae_Jonas_distinctus
Pilumnoididae_Pilumnoides_nudifrons
Pseudoziidae_Euryozius_camachoi
Pseudocarcinidae_Pseudocarcinus_gigas
Pseudocarcinidae_Pseudocarcinus_gigas_ourdata
Carpiliidae_Carpilius_maculatus
Carpiliidae_Carpilius_convexus
Carpiliidae_Carpilius_corallinus
Dairidae_Daira_americana
Dairidae_Daira_perlata
Cancridae_Anatolikos_japonicus
Cancridae_Cancer_borealis
Cancridae_Metacarcinus_magister
Bythograeidae_Austinograea_sp1and2
Calappidae_Mursia_sp2
Vultocinidae_Vultocinus_anfractus
Oregoniidae_Chionoecetes_tanneri
Oregoniidae_Hyas_lyratus
Conleyidae_Conleyus_sp
Mathildellidae_Mathildella_rubra
Eriphiidae_Eriphia_scabricula
Eriphiidae_Eriphia_smithii
Eriphiidae_Eriphia_verrucosa
Goneplacidae_Carcinoplax_purpurea
Goneplacidae_Goneplax_sp
Goneplacidae_Carcinoplax_longimana
Tanaochelidae_Tanaocheles_bidentata
Pilumnidae_Pilumnus_vespertilio
Pilumnidae_Heteropanope_glabra
Pilumnidae_Tiaramedon_spinosum
Galenidae_Galene_bispinosa
Pilumnidae_Pilumnus_longicornis
Pilumnidae_Lobopilumnus_agassizii
Pilumnidae_Pilumnus_dasypodus
Platyxanthidae_Homalaspis_plana
Platyxanthidae_Platyxanthus_orbignyi
Oziidae_Epixanthus_frontalis
Oziidae_Epixanthus_sp
Oziidae_Lydia_annulipes
Oziidae_Epixanthus_tenuidactylos
Oziidae_Ozius_reticulatus
Oziidae_Ozius_rugulosus
Scalopidiidae_Scalopidia_spinosipes
Platyxanthidae_Otmaroxanthus_balboai
Xanthidae_Demania_scaberrima
Xanthidae_Actaeodes_cf_tomentosus
Xanthidae_Novactaea_bella
Xanthidae_Atergatis_floridus
Xanthidae_Atergatis_integerrimus
Xanthidae_Xanthias_canaliculatus
Xanthidae_Edwardsium_lobipes
Xanthidae_Palapedia_cf_nitida
Xanthidae_Cyclodius_obscurus
Xanthidae_Pilodius_areolatus
Xanthidae_Xanthodius_cooksoni
Xanthidae_Xanthodius_sternberghii1and2
Xanthidae_Platyactaea_dovii
Xanthidae_Actaea_bifrons
Xanthidae_Xantho_pilipes
Xanthidae_Cataleptodius_floridanus
Xanthidae_Allactaea_lithostrota
Xanthidae_Carpoporus_papulosus
Xanthidae_Williamstimpsonia_denticulatus
Pseudorhombilidae_Pseudorhombila_quadridentata
Pseudorhombilidae_Trapezioplax_tridentata
Pseudorhombilidae_Lophoxanthus_lamellipes
Pseudorhombilidae_Nanoplax_xanthiformis
Pseudorhombilidae_Micropanope_sculptipes
Pseudorhombilidae_Panoplax_depressa
Panopeidae_Malacoplax_californiensis
Panopeidae_Hexapanopeus_angustifrons
Panopeidae_Panopeus_herbstii
Panopeidae_Eurytium_limosum
Panopeidae_Dyspanopeus_sayi
Panopeidae_Acantholobulus_bermudensis
Panopeidae_Glyptoplax_smithii
Panopeidae_Neopanope_packardii
Panopeidae_Rhithropanopeus_harrisii
Xanthidae_Lybia_tessellata
Ethusidae_Ethusa_sexdentata
Ethusidae_Parethusa_sp
Trichodactylidae_Trichodactylus_dentatus
Trichodactylidae_Fredilocarcinus_apyratii
Trichodactylidae_Valdivia_serrata
Pseudothelphusidae_Epilobocera_sinuatifrons
Dorippidae_Dorippe_quadridens
Dorippidae_Heikeopsis_japonica
Dorippidae_Paradorippe_granulata
Gecarcinucidae_Ceylonthelphusa_sp
Gecarcinucidae_Spiralothelphusa_senex
Gecarcinucidae_Somanniathelphusa_cf_zanklon
Gecarcinucidae_Somanniathelphusa_sp
Gecarcinucidae_Syntripsa_flavichela
Gecarcinucidae_Parathelphusa_pantherina
Gecarcinucidae_Parathelphusa_pantherina_ourdata
Majidae_Maja_brachydactyla
Majidae_Schizophrys_aspera
Cyclodorippidae_Corycodus_disjunctipes
Cyclodorippidae_Corycodus_sp
Cymonomidae_Cymonomus_diogenes
Cymonomidae_Cymonomus_sp
Raninidae_Ranilia_muricata
Lyreididae_Lyreidus_tridentatus
Lyreididae_Lysirude_griffini
Raninidae_Ranina_ranina
Raninidae_Raninoides_benedicti
Dynomenidae_Metadynomene_tanensis
Dynomenidae_Dynomene_praedator
Dromiidae_Lauridromia_dehaani
Dromiidae_Conchoecetes_artificiosus
Dromiidae_Lewindromia_unidentata
Dromiidae_Moreiradromia_antillensis
Homolidae_Lamoha_murotoensis
Homolidae_Lamoha_longirostris
Homolidae_Homola_orientalis
Homolidae_Homologenus_malayensis
Latreilliidae_Latreillia_valida
Latreilliidae_Latreillia_elegans
Orithyiidae_Orithyia_sinica
Chasmocarcinidae_Chasmocarcinus_mississippiensis
Belliidae_Acanthocyclus_albatrossis
Belliidae_Acanthocyclus_hassleri
Chasmocarcinidae_Megaesthesius_yokoyai
Inachoididae_Collodes_granosus
Inachoididae_Aepinus_septemspinosus
Inachoididae_Inachoides_laevis
Mictyridae_Mictyris_sp
Hymenosomatidae_Neorhynchoplax_bovis
Hymenosomatidae_Hymenosoma_orbiculare
Hymenosomatidae_Amarinus_paralacustris
Hymenosomatidae_Elamena_producta
Hymenosomatidae_Halicarcinus_cookii
Hymenosomatidae_Halicarcinus_innominatus
Macrophthalmidae_Macrophthalmus_telescopicus
Macrophthalmidae_Macrophthalmus_erato
Macrophthalmidae_Macrophthalmus_definitus
Macrophthalmidae_Chaenostoma_boscii
Ocypodidae_Ocypode_ceratophthalmus
Ocypodidae_Ocypode_sinensis
Varunidae_Metaplax_longipes
Varunidae_Gaetice_depressus
Varunidae_Eriocheir_japonica
Varunidae_Hemigrapsus_crenulatus
Varunidae_Hemigrapsus_penicillatus
Heloeciidae_Heloecius_cordiformis
Cryptochiridae_Dacryomaia_sp
Pinnotheridae_Arcotheres_sp
Xenograpsidae_Xenograpsus_ngatama
Xenograpsidae_Xenograpsus_testudinatus
Glyptograpsidae_Glyptograpsus_jamaicensis
Glyptograpsidae_Platychirograpsus_spectabilis
Percnidae_Percnon_planissimum
Percnidae_Percnon_affine
Percnidae_Percnon_affine_ourdata
Grapsidae_Pachygrapsus_plicatus
Ocypodidae_Gelasimus_borealis
Ocypodidae_Austruca_lactea
Ocypodidae_Paraleptuca_crassipes
Pinnotheridae_Nepinnotheres_cf_affinis
Pinnotheridae_Holothuriophilus_pacificus
Xenophthalmidae_Xenophthalmus_pinnotheroides
Pinnotheridae_Rathbunixa_pearsei
Pinnotheridae_Scleroplax_tubicola
Grapsidae_Grapsus_albolineatus
Grapsidae_Metopograpsus_quadridentatus
Grapsidae_Metopograpsus_frontalis
Grapsidae_Metopograpsus_messor
Gecarcinidae_Tuerkayana_hirtipes
Gecarcinidae_Cardisoma_crassum
Gecarcinidae_Cardisoma_guanhumi
Sesarmidae_Armases_cinereum
Sesarmidae_Parasesarma_affine
Sesarmidae_Parasesarma_eumolpe
Sesarmidae_Perisesarma_bidens
Gecarcinidae_Gecarcinus_lateralis
Gecarcinidae_Gecarcoidea_lalandii
Gecarcinidae_Gecarcoidea_natalis
Plagusiidae_Plagusia_squamosa
Dotillidae_Dotilla_myctiroides
Dotillidae_Dotilla_wichmanni
Dotillidae_Scopimera_proxima
Dotillidae_Scopimera_bitympana
Dotillidae_Scopimera_intermedia
0.09

**Fig. S2**

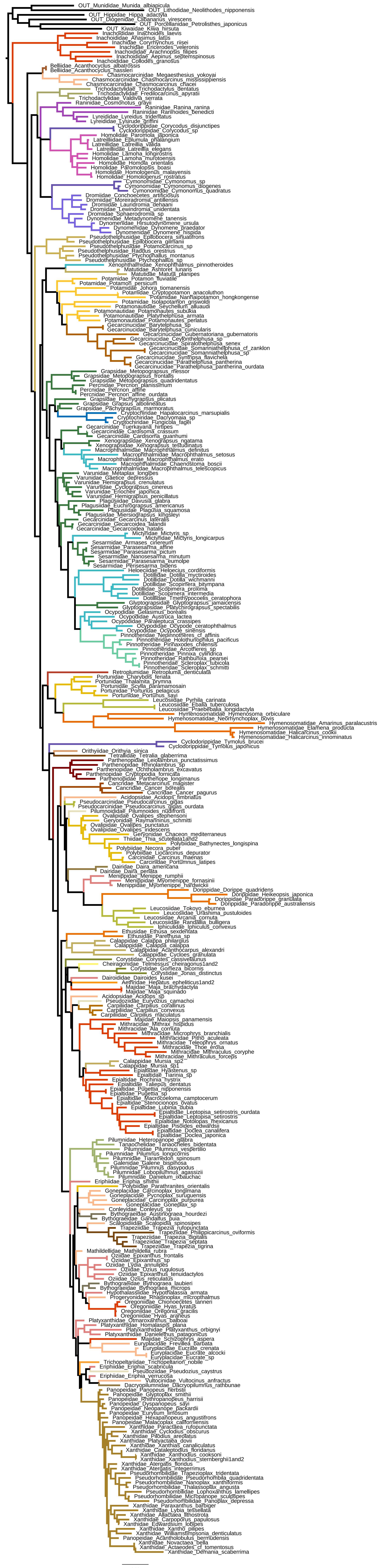

Fig. S3

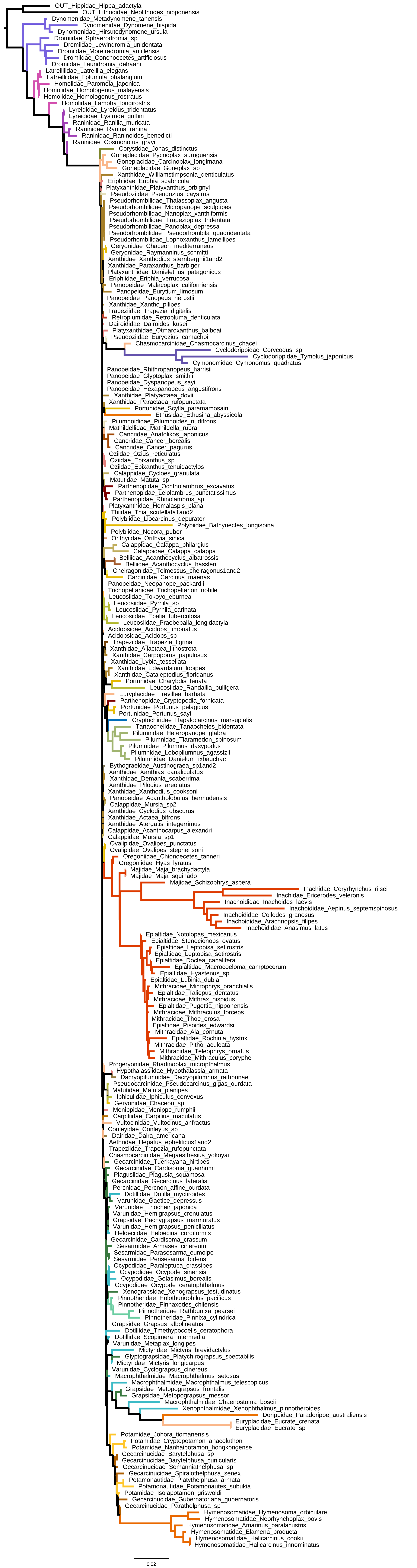

OUT_Hippidae_Hippa_adactyla
OUT_Lithodidae_Neolithodes_nipponensis
Dynomenidae_Metadynomene_tanensis
Dynomenidae_Dynomene_hispida
Dynomenidae_Hirsutodynomene_ursula
Dromiidae_Sphaerodromia_sp
Dromiidae_Lewindromia_unidentata
Dromiidae_Moreiradromia_antillensis
Dromiidae_Conchoecetes_artificiosus
Dromiidae_Lauridromia_dehaani
Latreilliidae_Latreillia_elegans
Latreilliidae_Eplumula_phalangium
Homolidae_Paromola_japonica
Homolidae_Homologenus_malayensis
Homolidae_Homologenus_rostratus
Homolidae_Lamoha_longirostris
Lyreididae_Lyreidus_tridentatus
Lyreididae_Lysirude_griffini
Raninidae_Ranilia_muricata
Raninidae_Ranina_ranina
Raninidae_Raninoides_benedicti
Raninidae_Cosmonotus_grayii
Corystidae_Jonas_distinctus
Goneplacidae_Pycnoplax_suruguensis
Goneplacidae_Carcinoplax_longimana
Goneplacidae_Goneplax_sp
Xanthidae_Williamstimpsonia_denticulatus
Eriphiidae_Eriphia_scabricula
Platyxanthidae_Platyxanthus_orbignyi
Pseudoziidae_Pseudozius_caystrus
Pseudorhombilidae_Thalassoplax_angusta
Pseudorhombilidae_Micropanope_sculptipes
Pseudorhombilidae_Nanoplax_xanthiformis
Pseudorhombilidae_Trapezioplax_tridentata
Pseudorhombilidae_Panoplax_depressa
Pseudorhombilidae_Pseudorhombila_quadridentata
Pseudorhombilidae_Lophoxanthus_lamellipes
Geryonidae_Chaceon_mediterraneus
Geryonidae_Raymanninus_schmitti
Xanthidae_Xanthodius_sternberghii1and2
Xanthidae_Paraxanthus_barbiger
Platyxanthidae_Danielethus_patagonicus
Eriphiidae_Eriphia_verrucosa
Panopeidae_Malacoplax_californiensis
Panopeidae_Eurytium_limosum
Panopeidae_Panopeus_herbstii
Xanthidae_Xantho_pilipes
Trapeziidae_Trapezia_digitalis
Retroplumidae_Retropluma_denticulata
Dairoididae_Dairoides_kusei
Platyxanthidae_Otmaroxanthus_balboai
Pseudoziidae_Euryozius_camachoi
Chasmocarcinidae_Chasmocarcinus_chacei
Cyclodorippidae_Corycodus_sp
Cyclodorippidae_Tymolus_japonicus
Cymonomidae_Cymonomus_quadratus
Panopeidae_Rhithropanopeus_harrisii
Panopeidae_Glyptoplax_smithii
Panopeidae_Dyspanopeus_sayi
Panopeidae_Hexapanopeus_angustifrons
Xanthidae_Platyactaea_dovii
Xanthidae_Paractaea_rufopunctata
Portunidae_Scylla_paramamosain
Ethusidae_Ethusina_abyssicola
Pilumnoididae_Pilumnoides_nudifrons
Mathildellidae_Mathildella_rubra
Cancridae_Anatolikos_japonicus
Cancridae_Cancer_borealis
Cancridae_Cancer_pagurus
Oziidae_Ozius_reticulatus
Oziidae_Epixanthus_sp
Oziidae_Epixanthus_tenuidactylos
Calappidae_Cycloes_granulata
Matutidae_Matuta_sp
Parthenopidae_Ochtholambrus_excavatus
Parthenopidae_Leiolambrus_punctatissimus
Parthenopidae_Rhinolambrus_sp
Platyxanthidae_Homalaspis_plana
Thiidae_Thia_scutellata1and2
Polybiidae_Liocarcinus_depurator
Polybiidae_Bathynectes_longispina
Polybiidae_Necora_puber
Orithyiidae_Orithyia_sinica
Calappidae_Calappa_philargius
Calappidae_Calappa_calappa
Belliidae_Acanthocyclus_albatrossis
Belliidae_Acanthocyclus_hassleri
Cheiragonidae_Telmessus_cheiragonus1and2
Carcinidae_Carcinus_maenas
Panopeidae_Neopanope_packardii
Trichopeltariidae_Trichopeltarion_nobile
Leucosiidae_Tokoyo_eburnea
Leucosiidae_Pyrhila_sp
Leucosiidae_Pyrhila_carinata
Leucosiidae_Ebalia_tuberculosa
Leucosiidae_Praebebalia_longidactyla
Acidopsidae_Acidops_fimbriatus
Acidopsidae_Acidops_sp
Trapeziidae_Trapezia_tigrina
Xanthidae_Allactaea_lithostrota
Xanthidae_Carpoporus_papulosus
Xanthidae_Lybia_tessellata
Xanthidae_Edwardsium_lobipes
Xanthidae_Cataleptodius_floridanus
Portunidae_Charybdis_feriata
Leucosiidae_Randallia_bulligera
Euryplacidae_Frevillea_barbata
Parthenopidae_Cryptopodia_fornicata
Portunidae_Portunus_pelagicus
Portunidae_Portunus_sayi
Cryptochiridae_Hapalocarcinus_marsupialis
Tanaochelidae_Tanaocheles_bidentata
Pilumnidae_Heteropanope_glabra
Pilumnidae_Tiaramedon_spinosum
Pilumnidae_Pilumnus_dasypodus
Pilumnidae_Lobopilumnus_agassizii
Pilumnidae_Danielum_ixbauchac
Bythograeidae_Austinograea_sp1and2
Xanthidae_Xanthias_canaliculatus
Xanthidae_Demania_scaberrima
Xanthidae_Pilodius_areolatus
Xanthidae_Xanthodius_cooksoni
Panopeidae_Acantholobulus_bermudensis
Calappidae_Mursia_sp2
Xanthidae_Cyclodius_obscurus
Xanthidae_Actaea_bifrons
Xanthidae_Atergatis_integerrimus
Calappidae_Acanthocarpus_alexandri
Calappidae_Mursia_sp1
Ovalipidae_Ovalipes_punctatus
Ovalipidae_Ovalipes_stephensoni
Oregoniidae_Chionoecetes_tanneri
Oregoniidae_Hyas_lyratus
Majidae_Maja_brachydactyla
Majidae_Maja_squinado
Majidae_Schizophrys_aspera
Inachidae_Coryrhynchus_riisei
Inachidae_Ericerodes_veleronis
Inachoididae_Inachoides_laevis
Inachoididae_Aepinus_septemspinosus
Inachoididae_Collodes_granosus
Inachoididae_Arachnopsis_filipes
Inachoididae_Anasimus_latus
Epialtidae_Notolopas_mexicanus
Epialtidae_Stenocionops_ovatus
Epialtidae_Leptopisa_setirostris
Epialtidae_Leptopisa_setirostris
Epialtidae_Doclea_canalifera
Epialtidae_Macrocoeloma_camptocerum
Epialtidae_Hyastenus_sp
Epialtidae_Lubinia_dubia
Mithracidae_Microphrys_branchialis
Epialtidae_Taliepus_dentatus
Mithracidae_Mithrax_hispidus
Epialtidae_Pugettia_nipponensis
Mithracidae_Mithraculus_forceps
Mithracidae_Thoe_erosa
Epialtidae_Pisoides_edwardsii
Mithracidae_Ala_cornuta
Epialtidae_Rochinia_hystrix
Mithracidae_Pitho_aculeata
Mithracidae_Teleophrys_ornatus
Mithracidae_Mithraculus_coryphe
Progeryonidae_Rhadinoplax_micropthalmus
Hypothalassiidae_Hypothalassia_armata
Dacryopilumnidae_Dacryopilumnus_rathbunae
Pseudocarcinidae_Pseudocarcinus_gigas_ourdata
Matutidae_Matuta_planipes
Iphiculidae_Iphiculus_convexus
Geryonidae_Chaceon_sp
Menippidae_Menippe_rumphii
Carpiliidae_Carpilius_maculatus
Vultocinidae_Vultocinus_anfractus
Conleyidae_Conleyus_sp
Dairidae_Daira_americana
Aethridae_Hepatus_epheliticus1and2
Trapeziidae_Trapezia_rufopunctata
Chasmocarcinidae_Megaesthesius_yokoyai
Gecarcinidae_Tuerkayana_hirtipes
Gecarcinidae_Cardisoma_guanhumi
Plagusiidae_Plagusia_squamosa
Gecarcinidae_Gecarcinus_lateralis
Percnidae_Percnon_affine_ourdata
Dotillidae_Dotilla_myctiroides
Varunidae_Gaetice_depressus
Varunidae_Eriocheir_japonica
Varunidae_Hemigrapsus_crenulatus
Grapsidae_Pachygrapsus_marmoratus
Varunidae_Hemigrapsus_penicillatus
Heloeciidae_Heloecius_cordiformis
Gecarcinidae_Cardisoma_crassum
Sesarmidae_Armases_cinereum
Sesarmidae_Parasesarma_eumolpe
Sesarmidae_Perisesarma_bidens
Ocypodidae_Paraleptuca_crassipes
Ocypodidae_Ocypode_sinensis
Ocypodidae_Gelasimus_borealis
Ocypodidae_Ocypode_ceratophthalmus
Xenograpsidae_Xenograpsus_testudinatus
Pinnotheridae_Holothuriophilus_pacificus
Pinnotheridae_Pinnaxodes_chilensis
Pinnotheridae_Rathbunixa_pearsei
Pinnotheridae_Pinnixa_cylindrica
Grapsidae_Grapsus_albolineatus
Dotillidae_Tmethypocoelis_ceratophora
Dotillidae_Scopimera_intermedia
Varunidae_Metaplax_longipes
Mictyridae_Mictyris_brevidactylus
Glyptograpsidae_Platychirograpsus_spectabilis
Mictyridae_Mictyris_longicarpus
Varunidae_Cyclograpsus_cinereus
Macrophthalmidae_Macrophthalmus_setosus
Macrophthalmidae_Macrophthalmus_telescopicus
Grapsidae_Metopograpsus_frontalis
Grapsidae_Metopograpsus_messor
Macrophthalmidae_Chaenostoma_boscii
Xenophthalmidae_Xenophthalmus_pinnotheroides
Dorippidae_Paradorippe_australiensis
Euryplacidae_Eucrate_crenata
Euryplacidae_Eucrate_sp
Potamidae_Johora_tiomanensis
Potamidae_Cryptopotamon_anacoluthon
Potamidae_Nanhaipotamon_hongkongense
Gecarcinucidae_Barytelphusa_sp
Gecarcinucidae_Barytelphusa_cunicularis
Gecarcinucidae_Somanniathelphusa_sp
Gecarcinucidae_Spiralothelphusa_senex
Potamonautidae_Platythelphusa_armata
Potamonautidae_Potamonautes_subukia
Potamidae_Isolapotamon_griswoldi
Gecarcinucidae_Gubernatoriana_gubernatoris
Gecarcinucidae_Parathelphusa_sp
Hymenosomatidae_Hymenosoma_orbiculare
Hymenosomatidae_Neorhynchoplax_bovis
Hymenosomatidae_Amarinus_paralacustris
Hymenosomatidae_Elamena_producta
Hymenosomatidae_Halicarcinus_cookii
Hymenosomatidae_Halicarcinus_innominatus
0.02

**Fig. S4**

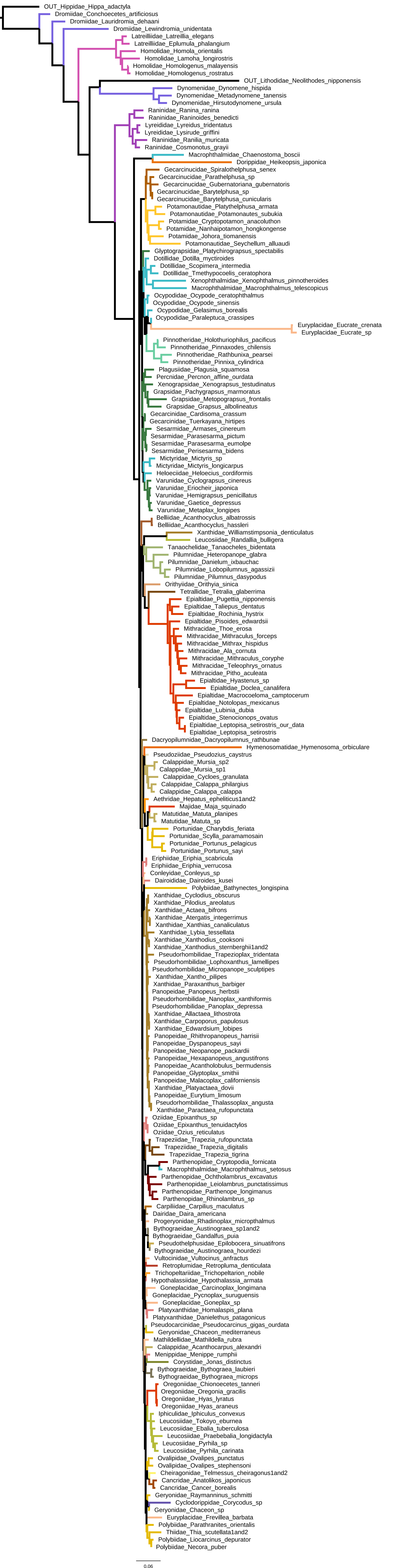
OUT_Hippidae_Hippa_adactyla
Dromiidae_Conchoecetes_artificiosus
Dromiidae_Lauridromia_dehaani
Dromiidae_Lewindromia_unidentata
Latreilliidae_Latreillia_elegans
Latreilliidae_Eplumula_phalangium
Homolidae_Homola_orientalis
Homolidae_Lamoha_longirostris
Homolidae_Homologenus_malayensis
Homolidae_Homologenus_rostratus
OUT_Lithodidae_Neolithodes_nipponensis
Dynomenidae_Dynomene_hispida
Dynomenidae_Metadynomene_tanensis
Dynomenidae_Hirsutodynomene_ursula
Raninidae_Ranina_ranina
Raninidae_Raninoides_benedicti
Lyreididae_Lyreidus_tridentatus
Lyreididae_Lysirude_griffini
Raninidae_Ranilia_muricata
Raninidae_Cosmonotus_grayii
Macrophthalmidae_Chaenostoma_boscii
Dorippidae_Heikeopsis_japonica
Gecarcinucidae_Spiralothelphusa_senex
Gecarcinucidae_Parathelphusa_sp
Gecarcinucidae_Gubernatoriana_gubernatoris
Gecarcinucidae_Barytelphusa_sp
Gecarcinucidae_Barytelphusa_cunicularis
Potamonautidae_Platythelphusa_armata
Potamonautidae_Potamonautes_subukia
Potamidae_Cryptopotamon_anacoluthon
Potamidae_Nanhaipotamon_hongkongense
Potamidae_Johora_tiomanensis
Potamonautidae_Seychellum_alluaudi
Glyptograpsidae_Platychirograpsus_spectabilis
Dotillidae_Dotilla_myctiroides
Dotillidae_Scopimera_intermedia
Dotillidae_Tmethypocoelis_ceratophora
Xenophthalmidae_Xenophthalmus_pinnotheroides
Macrophthalmidae_Macrophthalmus_telescopicus
Ocypodidae_Ocypode_ceratophthalmus
Ocypodidae_Ocypode_sinensis
Ocypodidae_Gelasimus_borealis
Ocypodidae_Paraleptuca_crassipes
Euryplacidae_Eucrate_crenata
Euryplacidae_Eucrate_sp
Pinnotheridae_Holothuriophilus_pacificus
Pinnotheridae_Pinnaxodes_chilensis
Pinnotheridae_Rathbunixa_pearsei
Pinnotheridae_Pinnixa_cylindrica
Plagusiidae_Plagusia_squamosa
Percnidae_Percnon_affine_ourdata
Xenograpsidae_Xenograpsus_testudinatus
Grapsidae_Pachygrapsus_marmoratus
Grapsidae_Metopograpsus_frontalis
Grapsidae_Grapsus_albolineatus
Gecarcinidae_Cardisoma_crassum
Gecarcinidae_Tuerkayana_hirtipes
Sesarmidae_Armases_cinereum
Sesarmidae_Parasesarma_pictum
Sesarmidae_Parasesarma_eumolpe
Sesarmidae_Perisesarma_bidens
Mictyridae_Mictyris_sp
Mictyridae_Mictyris_longicarpus
Heloeciidae_Heloecius_cordiformis
Varunidae_Cyclograpsus_cinereus
Varunidae_Eriocheir_japonica
Varunidae_Hemigrapsus_penicillatus
Varunidae_Gaetice_depressus
Varunidae_Metaplax_longipes
Belliidae_Acanthocyclus_albatrossis
Belliidae_Acanthocyclus_hassleri
Xanthidae_Williamstimpsonia_denticulatus
Leucosiidae_Randallia_bulligera
Tanaochelidae_Tanaocheles_bidentata
Pilumnidae_Heteropanope_glabra
Pilumnidae_Danielum_ixbauchac
Pilumnidae_Lobopilumnus_agassizii
Pilumnidae_Pilumnus_dasypodus
Orithyiidae_Orithyia_sinica
Tetraliidae_Tetralia_glaberrima
Epialtidae_Pugettia_nipponensis
Epialtidae_Taliepus_dentatus
Epialtidae_Rochinia_hystrix
Epialtidae_Pisoides_edwardsii
Mithracidae_Thoe_erosa
Mithracidae_Mithraculus_forceps
Mithracidae_Mithrax_hispidus
Mithracidae_Ala_cornuta
Mithracidae_Mithraculus_coryphe
Mithracidae_Teleophrys_ornatus
Mithracidae_Pitho_aculeata
Epialtidae_Hyastenus_sp
Epialtidae_Doclea_canalifera
Epialtidae_Macrocoeloma_camptocerum
Epialtidae_Notolopas_mexicanus
Epialtidae_Lubinia_dubia
Epialtidae_Stenocionops_ovatus
Epialtidae_Leptopisa_setirostris_our_data
Epialtidae_Leptopisa_setirostris
Dacryopilumnidae_Dacryopilumnus_rathbunae
Hymenosomatidae_Hymenosoma_orbiculare
Pseudoziidae_Pseudozius_caystrus
Calappidae_Mursia_sp2
Calappidae_Mursia_sp1
Calappidae_Cycloes_granulata
Calappidae_Calappa_philargius
Calappidae_Calappa_calappa
Aethridae_Hepatus_epheliticus1and2
Majidae_Maja_squinado
Matutidae_Matuta_planipes
Matutidae_Matuta_sp
Portunidae_Charybdis_feriata
Portunidae_Scylla_paramamosain
Portunidae_Portunus_pelagicus
Portunidae_Portunus_sayi
Eriphiidae_Eriphia_scabricula
Eriphiidae_Eriphia_verrucosa
Conleyidae_Conleyus_sp
Dairoididae_Dairoides_kusei
Polybiidae_Bathynectes_longispina
Xanthidae_Cyclodius_obscurus
Xanthidae_Pilodius_areolatus
Xanthidae_Actaea_bifrons
Xanthidae_Atergatis_integerrimus
Xanthidae_Xanthias_canaliculatus
Xanthidae_Lybia_tessellata
Xanthidae_Xanthodius_cooksoni
Xanthidae_Xanthodius_sternberghii1and2
Pseudorhombilidae_Trapezioplax_tridentata
Pseudorhombilidae_Lophoxanthus_lamellipes
Pseudorhombilidae_Micropanope_sculptipes
Xanthidae_Xantho_pilipes
Xanthidae_Paraxanthus_barbiger
Panopeidae_Panopeus_herbstii
Pseudorhombilidae_Nanoplax_xanthiformis
Pseudorhombilidae_Panoplax_depressa
Xanthidae_Alllactaea_lithostrota
Xanthidae_Carpoporus_papulosus
Xanthidae_Edwardsium_lobipes
Panopeidae_Rhithropanopeus_harrisii
Panopeidae_Dyspanopeus_sayi
Panopeidae_Neopanope_packardii
Panopeidae_Hexapanopeus_angustifrons
Panopeidae_Acantholobulus_bermudensis
Panopeidae_Glyptoplax_smithii
Panopeidae_Malacoplax_californiensis
Xanthidae_Platyactaea_dovii
Panopeidae_Eurytium_limosum
Pseudorhombilidae_Thalassoplax_angusta
Xanthidae_Paractaea_rufopunctata
Oziidae_Epixanthus_sp
Oziidae_Epixanthus_tenuidactylos
Oziidae_Ozius_reticulatus
Trapeziidae_Trapezia_rufopunctata
Trapeziidae_Trapezia_digitalis
Trapeziidae_Trapezia_tigrina
Parthenopidae_Cryptopodia_fornicata
Macrophthalmidae_Macrophthalmus_setosus
Parthenopidae_Ochtholambrus_excavatus
Parthenopidae_Leiolambrus_punctatissimus
Parthenopidae_Parthenope_longimanus
Parthenopidae_Rhinolambrus_sp
Carpiliidae_Carpilius_maculatus
Dairidae_Daira_americana
Progeryonidae_Rhadinoplax_micropthalmus
Bythograeidae_Austinograea_sp1and2
Bythograeidae_Gandalfus_puia
Pseudothelphusidae_Epilobocera_sinuatifrons
Bythograeidae_Austinograea_hourdezi
Vultocinidae_Vultocinus_anfractus
Retroplumidae_Retropluma_denticulata
Trichopeltariidae_Trichopeltarion_nobile
Hypothalassiidae_Hypothalassia_armata
Goneplacidae_Carcinoplax_longimana
Goneplacidae_Pycnoplax_surugensis
Goneplacidae_Goneplax_sp
Platyxanthidae_Homalaspis_plana
Platyxanthidae_Danielethus_patagonicus
Pseudocarcinidae_Pseudocarcinus_gigas_ourdata
Geryonidae_Chaceon_mediterraneus
Mathildellidae_Mathildella_rubra
Calappidae_Acanthocarpus_alexandri
Menippidae_Menippe_rumphii
Corystidae_Jonas_distinctus
Bythograeidae_Bythograea_laubieri
Bythograeidae_Bythograea_microps
Oregoniidae_Chionoecetes_tanneri
Oregoniidae_Oregonia_gracilis
Oregoniidae_Hyas_lyratus
Oregoniidae_Hyas_araneus
Iphiculidae_Iphiculus_convexus
Leucosiidae_Tokoyo_eburnea
Leucosiidae_Ebalia_tuberculosa
Leucosiidae_Praebebalia_longidactyla
Leucosiidae_Pyrhila_sp
Leucosiidae_Pyrhila_carinata
Ovalipidae_Ovalipes_punctatus
Ovalipidae_Ovalipes_stephensoni
Cheiragonidae_Telmessus_cheiragonus1and2
Cancridae_Anatolikos_japonicus
Cancridae_Cancer_borealis
Geryonidae_Raymanninus_schmitti
Cyclodorippidae_Corycodus_sp
Geryonidae_Chaceon_sp
Euryplacidae_Frevillea_barbata
Polybiidae_Parathranites_orientalis
Thiidae_Thia_scutellata1and2
Polybiidae_Liocarcinus_depurator
Polybiidae_Necora_puber
0.06

**Fig. S5**

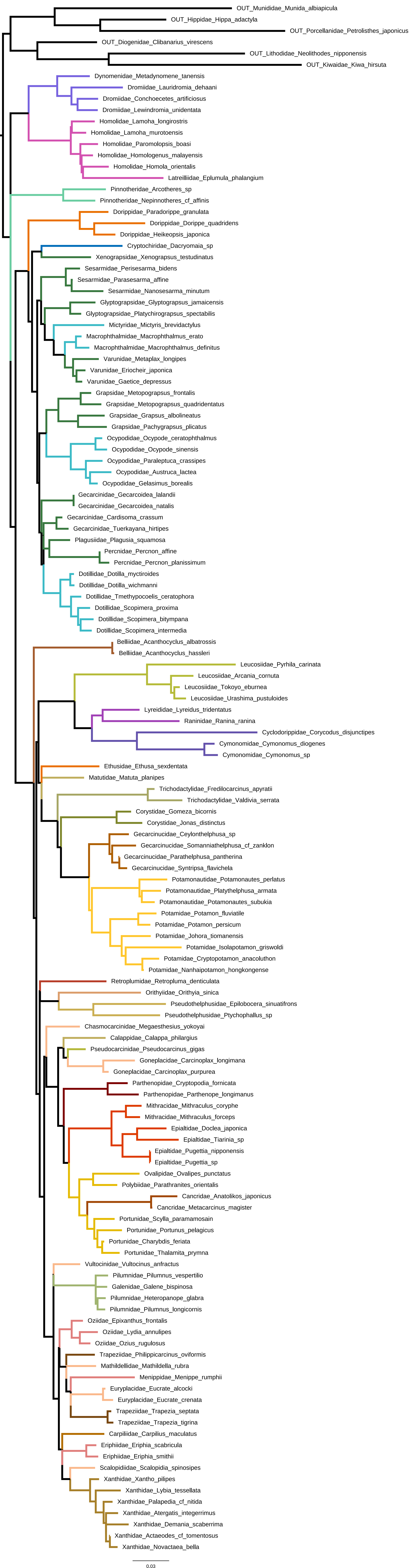
OUT_Munididae_Munida_albiapicula
OUT_Hippidae_Hippa_adactyla
OUT_Porcellanidae_Petrolisthes_japonicus
OUT_Diogenidae_Clibanarius_virescens
OUT_Lithodidae_Neolithodes_nipponensis
OUT_Kiwaidae_Kiwa_hirsuta
Dynomenidae_Metadynomene_tanensis
Dromiidae_Lauridromia_dehaani
Dromiidae_Conchoecetes_artificiosus
Dromiidae_Lewindromia_unidentata
Homolidae_Lamoha_longirostris
Homolidae_Lamoha_murotoensis
Homolidae_Paromolopsis_boasi
Homolidae_Homologenus_malayensis
Homolidae_Homola_orientalis
Latreilliidae_Eplumula_phalangium
Pinnotheridae_Arcotheres_sp
Pinnotheridae_Nepinnotheres_cf_affinis
Dorippidae_Paradorippe_granulata
Dorippidae_Dorippe_quadridens
Dorippidae_Heikeopsis_japonica
Cryptochiridae_Dacryomaia_sp
Xenograpsidae_Xenograpsus_testudinatus
Sesarmidae_Perisesarma_bidens
Sesarmidae_Parasesarma_affine
Sesarmidae_Nanosesarma_minutum
Glyptograpsidae_Glyptograpsus_jamaicensis
Glyptograpsidae_Platychirograpsus_spectabilis
Mictyridae_Mictyris_brevidactylus
Macrophthalmidae_Macrophthalmus_erato
Macrophthalmidae_Macrophthalmus_definitus
Varunidae_Metaplax_longipes
Varunidae_Eriocheir_japonica
Varunidae_Gaetice_depressus
Grapsidae_Metopograpsus_frontalis
Grapsidae_Metopograpsus_quadridentatus
Grapsidae_Grapsus_albolineatus
Grapsidae_Pachygrapsus_plicatus
Ocypodidae_Ocypode_ceratophthalmus
Ocypodidae_Ocypode_sinensis
Ocypodidae_Paraleptuca_crassipes
Ocypodidae_Austruca_lactea
Ocypodidae_Gelasimus_borealis
Gecarcinidae_Gecarcoidea_lalandii
Gecarcinidae_Gecarcoidea_natalis
Gecarcinidae_Cardisoma_crassum
Gecarcinidae_Tuerkayana_hirtipes
Plagusiidae_Plagusia_squamosa
Percnidae_Percnon_affine
Percnidae_Percnon_planissimum
Dotillidae_Dotilla_myctiroides
Dotillidae_Dotilla_wichmanni
Dotillidae_Tmethypocoelis_ceratophora
Dotillidae_Scopimera_proxima
Dotillidae_Scopimera_bitympana
Dotillidae_Scopimera_intermedia
Belliidae_Acanthocyclus_albatrossis
Belliidae_Acanthocyclus_hassleri
Leucosiidae_Pyrhila_carinata
Leucosiidae_Arcania_cornuta
Leucosiidae_Tokoyo_eburnea
Leucosiidae_Urashima_pustuloides
Lyreididae_Lyreidus_tridentatus
Raninidae_Ranina_ranina
Cyclodorippidae_Corycodus_disjunctipes
Cymonomidae_Cymonomus_diogenes
Cymonomidae_Cymonomus_sp
Ethusidae_Ethusa_sexdentata
Matutidae_Matuta_planipes
Trichodactylidae_Fredilocarcinus_apyratii
Trichodactylidae_Valdivia_serrata
Corystidae_Gomeza_bicornis
Corystidae_Jonas_distinctus
Gecarcinucidae_Ceylonthelphusa_sp
Gecarcinucidae_Somanniathelphusa_cf_zanklon
Gecarcinucidae_Parathelphusa_pantherina
Gecarcinucidae_Syntripsa_flavichela
Potamonautidae_Potamonautes_perlatus
Potamonautidae_Platythelphusa_armata
Potamonautidae_Potamonautes_subukia
Potamidae_Potamon_fluviatile
Potamidae_Potamon_persicum
Potamidae_Johora_tiomanensis
Potamidae_Isolapotamon_griswoldi
Potamidae_Cryptopotamon_anacoluthon
Potamidae_Nanhaipotamon_hongkongense
Retroplumidae_Retropluma_denticulata
Orithyiidae_Orithyia_sinica
Pseudothelphusidae_Epilobocera_sinuatifrons
Pseudothelphusidae_Ptychophallus_sp
Chasmocarcinidae_Megaesthesius_yokoyai
Calappidae_Calappa_philargius
Pseudocarcinidae_Pseudocarcinus_gigas
Goneplacidae_Carcinoplax_longimana
Goneplacidae_Carcinoplax_purpurea
Parthenopidae_Cryptopodia_fornicata
Parthenopidae_Parthenope_longimanus
Mithracidae_Mithraculus_coryphe
Mithracidae_Mithraculus_forceps
Epialtidae_Doclea_japonica
Epialtidae_Tiarinia_sp
Epialtidae_Pugettia_nipponensis
Epialtidae_Pugettia_sp
Ovalipidae_Ovalipes_punctatus
Polybiidae_Parathranites_orientalis
Cancridae_Anatolikos_japonicus
Cancridae_Metacarcinus_magister
Portunidae_Scylla_paramamosain
Portunidae_Portunus_pelagicus
Portunidae_Charybdis_feriata
Portunidae_Thalamita_prymna
Vultocinidae_Vultocinus_anfractus
Pilumnidae_Pilumnus_vespertilio
Galenidae_Galene_bispinosa
Pilumnidae_Heteropanope_glabra
Pilumnidae_Pilumnus_longicornis
Oziidae_Epixanthus_frontalis
Oziidae_Lydia_annulipes
Oziidae_Ozius_rugulosus
Trapeziidae_Philippicarcinus_oviformis
Mathildellidae_Mathildella_rubra
Menippidae_Menippe_rumphii
Euryplacidae_Eucrate_alcocki
Euryplacidae_Eucrate_crenata
Trapeziidae_Trapezia_septata
Trapeziidae_Trapezia_tigrina
Carpiliidae_Carpilius_maculatus
Eriphiidae_Eriphia_scabricula
Eriphiidae_Eriphia_smithii
Scalopidiidae_Scalopidia_spinosipes
Xanthidae_Xantho_pilipes
Xanthidae_Lybia_tessellata
Xanthidae_Palapedia_cf_nitida
Xanthidae_Atergatis_integerrimus
Xanthidae_Demania_scaberrima
Xanthidae_Actaeodes_cf_tomentosus
Xanthidae_Novactaea_bella
0.03

**Fig. S6**

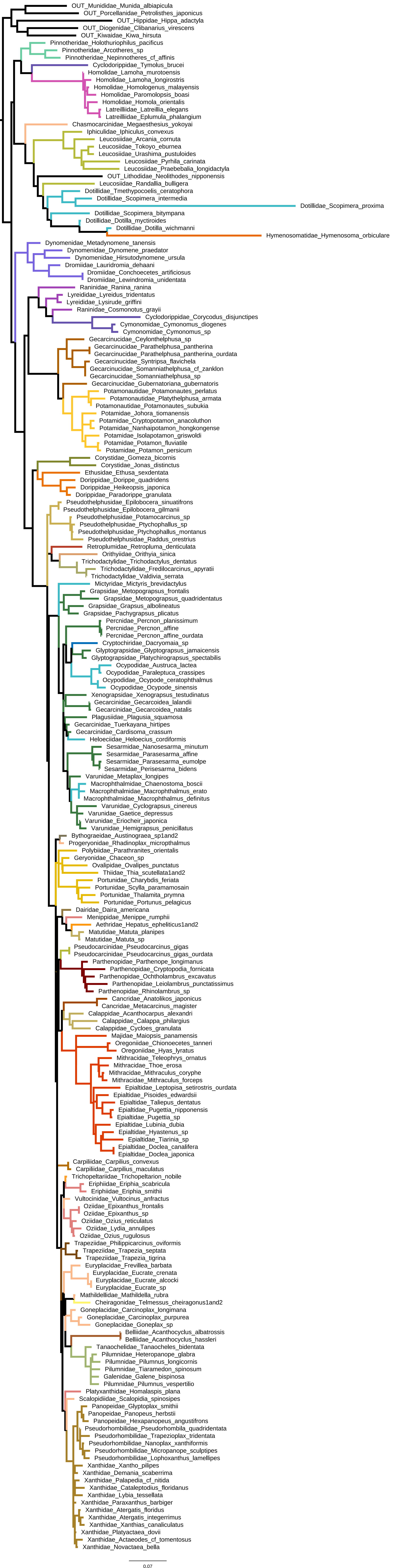
OUT_Munididae_Munida_albiapicula
OUT_Porcellanidae_Petrolisthes_japonicus
OUT_Hippidae_Hippa_adactyla
OUT_Diogenidae_Clibanarius_virescens
OUT_Kiwaidae_Kiwa_hirsuta
Pinnotheridae_Holothuriophilus_pacificus
Pinnotheridae_Arcotheres_sp
Pinnotheridae_Nepinnotheres_cf_affinis
Cyclodorippidae_Tymolus_brucei
Homolidae_Lamoha_murotoensis
Homolidae_Lamoha_longirostris
Homolidae_Homologenus_malayensis
Homolidae_Paromolopsis_boasi
Homolidae_Homola_orientalis
Latreilliidae_Latreillia_elegans
Latreilliidae_Eplumula_phalangium
Chasmocarcinidae_Megaesthesius_yokoyai
Iphiculidae_Iphiculus_convexus
Leucosiidae_Arcania_cornuta
Leucosiidae_Tokoyo_eburnea
Leucosiidae_Urashima_pustuloides
Leucosiidae_Pyrhila_carinata
Leucosiidae_Praebebalia_longidactyla
OUT_Lithodidae_Neolithodes_nipponensis
Leucosiidae_Randallia_bulligera
Dotillidae_Tmethypocoelis_ceratophora
Dotillidae_Scopimera_intermedia
Dotillidae_Scopimera_proxima
Dotillidae_Scopimera_bitympana
Dotillidae_Dotilla_myctiroides
Dotillidae_Dotilla_wichmanni
Hymenosomatidae_Hymenosoma_orbiculare
Dynomenidae_Metadynomene_tanensis
Dynomenidae_Dynomene_praedator
Dynomenidae_Hirsutodynomene_ursula
Dromiidae_Lauridromia_dehaani
Dromiidae_Conchoecetes_artificiosus
Dromiidae_Lewindromia_unidentata
Raninidae_Ranina_ranina
Lyreididae_Lyreidus_tridentatus
Lyreididae_Lysirude_griffini
Raninidae_Cosmonotus_grayii
Cyclodorippidae_Corycodus_disjunctipes
Cymonomidae_Cymonomus_diogenes
Cymonomidae_Cymonomus_sp
Gecarcinucidae_Ceylonthelphusa_sp
Gecarcinucidae_Parathelphusa_pantherina
Gecarcinucidae_Parathelphusa_pantherina_ourdata
Gecarcinucidae_Syntripsa_flavichela
Gecarcinucidae_Somanniathelphusa_cf_zanklon
Gecarcinucidae_Somanniathelphusa_sp
Gecarcinucidae_Gubernatoriana_gubernatoris
Potamonautidae_Potamonautes_perlatus
Potamonautidae_Platythelphusa_armata
Potamonautidae_Potamonautes_subukia
Potamidae_Johora_tiomanensis
Potamidae_Cryptopotamon_anacoluthon
Potamidae_Nanhaipotamon_hongkongense
Potamidae_Isolapotamon_griswoldi
Potamidae_Potamon_fluviatile
Potamidae_Potamon_persicum
Corystidae_Gomeza_bicornis
Corystidae_Jonas_distinctus
Ethusidae_Ethusa_sexdentata
Dorippidae_Dorippe_quadridens
Dorippidae_Heikeopsis_japonica
Dorippidae_Paradorippe_granulata
Pseudothelphusidae_Epilobocera_sinuatifrons
Pseudothelphusidae_Epilobocera_gilmanii
Pseudothelphusidae_Potamocarcinus_sp
Pseudothelphusidae_Ptychophallus_sp
Pseudothelphusidae_Ptychophallus_montanus
Pseudothelphusidae_Raddus_orestrius
Retroplumidae_Retropluma_denticulata
Orithyiidae_Orithyia_sinica
Trichodactylidae_Trichodactylus_dentatus
Trichodactylidae_Fredilocarcinus_apyratii
Trichodactylidae_Valdivia_serrata
Mictyridae_Mictyris_brevidactylus
Grapsidae_Metopograpsus_frontalis
Grapsidae_Metopograpsus_quadridentatus
Grapsidae_Grapsus_albolineatus
Grapsidae_Pachygrapsus_plicatus
Percnidae_Percnon_planissimum
Percnidae_Percnon_affine
Percnidae_Percnon_affine_ourdata
Cryptochiridae_Dacryomaia_sp
Glyptograpsidae_Glyptograpsus_jamaicensis
Glyptograpsidae_Platychirograpsus_spectabilis
Ocypodidae_Austruca_lactea
Ocypodidae_Paraleptuca_crassipes
Ocypodidae_Ocypode_ceratophthalmus
Ocypodidae_Ocypode_sinensis
Xenograpsidae_Xenograpsus_testudinatus
Gecarcinidae_Gecarcoidea_lalandii
Gecarcinidae_Gecarcoidea_natalis
Plagusiidae_Plagusia_squamosa
Gecarcinidae_Tuerkayana_hirtipes
Gecarcinidae_Cardisoma_crassum
Heloeciidae_Heloecius_cordiformis
Sesarmidae_Nanosesarma_minutum
Sesarmidae_Parasesarma_affine
Sesarmidae_Parasesarma_eumolpe
Sesarmidae_Perisesarma_bidens
Varunidae_Metaplax_longipes
Macrophthalmidae_Chaenostoma_boscii
Macrophthalmidae_Macrophthalmus_erato
Macrophthalmidae_Macrophthalmus_definitus
Varunidae_Cyclograpsus_cinereus
Varunidae_Gaetice_depressus
Varunidae_Eriocheir_japonica
Varunidae_Hemigrapsus_penicillatus
Bythograeidae_Austinograea_sp1and2
Progeryonidae_Rhadinoplax_micropthalmus
Polybiidae_Parathranites_orientalis
Geryonidae_Chaceon_sp
Ovalipidae_Ovalipes_punctatus
Thiidae_Thia_scutellata1and2
Portunidae_Charybdis_feriata
Portunidae_Scylla_paramamosain
Portunidae_Thalamita_prymna
Portunidae_Portunus_pelagicus
Dairidae_Daira_americana
Menippidae_Menippe_rumphii
Aethridae_Hepatus_epheliticus1and2
Matutidae_Matuta_planipes
Matutidae_Matuta_sp
Pseudocarcinidae_Pseudocarcinus_gigas
Pseudocarcinidae_Pseudocarcinus_gigas_ourdata
Parthenopidae_Parthenope_longimanus
Parthenopidae_Cryptopodia_fornicata
Parthenopidae_Ochtholambrus_excavatus
Parthenopidae_Leiolambrus_punctatissimus
Parthenopidae_Rhinolambrus_sp
Cancridae_Anatolikos_japonicus
Cancridae_Metacarcinus_magister
Calappidae_Acanthocarpus_alexandri
Calappidae_Calappa_philargius
Calappidae_Cycloes_granulata
Majidae_Maiopsis_panamensis
Oregoniidae_Chionoecetes_tanneri
Oregoniidae_Hyas_lyratus
Mithracidae_Teleophrys_ornatus
Mithracidae_Thoe_erosa
Mithracidae_Mithraculus_coryphe
Mithracidae_Mithraculus_forceps
Epialtidae_Leptopisa_setirostris_ourdata
Epialtidae_Pisoides_edwardsii
Epialtidae_Taliepus_dentatus
Epialtidae_Pugettia_nipponensis
Epialtidae_Pugettia_sp
Epialtidae_Lubinia_dubia
Epialtidae_Hyastenus_sp
Epialtidae_Tiarinia_sp
Epialtidae_Doclea_canalifera
Epialtidae_Doclea_japonica
Carpiliidae_Carpilius_convexus
Carpiliidae_Carpilius_maculatus
Trichopeltariidae_Trichopeltarion_nobile
Eriphiidae_Eriphia_scabricula
Eriphiidae_Eriphia_smithii
Vultocinidae_Vultocinus_anfractus
Oziidae_Epixanthus_frontalis
Oziidae_Epixanthus_sp
Oziidae_Ozius_reticulatus
Oziidae_Lydia_annulipes
Oziidae_Ozius_rugulosus
Trapeziidae_Philippicarcinus_oviformis
Trapeziidae_Trapezia_septata
Trapeziidae_Trapezia_tigrina
Euryplacidae_Frevillea_barbata
Euryplacidae_Eucrate_crenata
Euryplacidae_Eucrate_alcocki
Euryplacidae_Eucrate_sp
Mathildellidae_Mathildella_rubra
Cheiragonidae_Telmessus_cheiragonus1and2
Goneplacidae_Carcinoplax_longimana
Goneplacidae_Carcinoplax_purpurea
Goneplacidae_Goneplax_sp
Belliidae_Acanthocyclus_albatrossis
Belliidae_Acanthocyclus_hassleri
Tanaochelidae_Tanaocheles_bidentata
Pilumnidae_Heteropanope_glabra
Pilumnidae_Pilumnus_longicornis
Pilumnidae_Tiaramedon_spinosum
Galenidae_Galene_bispinosa
Pilumnidae_Pilumnus_vespertilio
Platyxanthidae_Homalaspis_plana
Scalopidiidae_Scalopidia_spinosipes
Panopeidae_Glyptoplax_smithii
Panopeidae_Panopeus_herbstii
Panopeidae_Hexapanopeus_angustifrons
Pseudorhombilidae_Pseudorhombila_quadridentata
Pseudorhombilidae_Trapezioplax_tridentata
Pseudorhombilidae_Nanoplax_xanthiformis
Pseudorhombilidae_Micropanope_sculptipes
Pseudorhombilidae_Lophoxanthus_lamellipes
Xanthidae_Xantho_pilipes
Xanthidae_Demania_scaberrima
Xanthidae_Palapedia_cf_nitida
Xanthidae_Cataleptodius_floridanus
Xanthidae_Lybia_tessellata
Xanthidae_Paraxanthus_barbiger
Xanthidae_Atergatis_floridus
Xanthidae_Atergatis_integerrimus
Xanthidae_Xanthias_canaliculatus
Xanthidae_Platyactaea_dovii
Xanthidae_Actaeodes_cf_tomentosus
Xanthidae_Novactaea_bella
0.07

**Fig. S7**

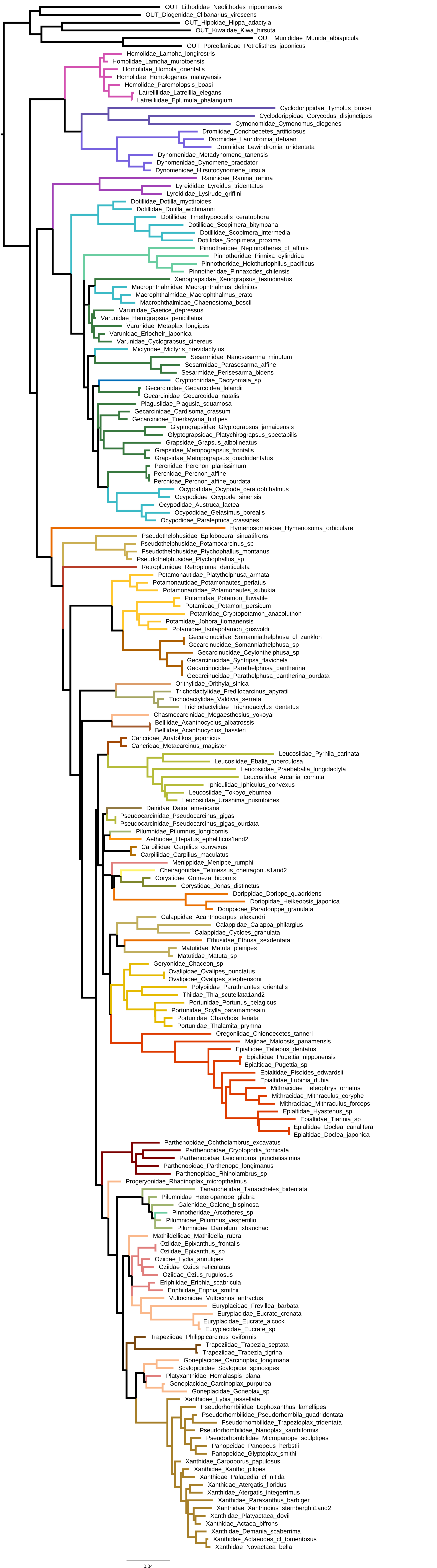
OUT_Lithodidae_Neolithodes_nipponensis
OUT_Diogenidae_Clibanarius_virescens
OUT_Hippidae_Hippa_adactyla
OUT_Kiwaidae_Kiwa_hirsuta
OUT_Munididae_Munida_albiapicula
OUT_Porcellanidae_Petrolisthes_japonicus
Homolidae_Lamoha_longirostris
Homolidae_Lamoha_murotoensis
Homolidae_Homola_orientalis
Homolidae_Homologenus_malayensis
Homolidae_Paromolopsis_boasi
Latreilliidae_Latreillia_elegans
Latreilliidae_Eplumula_phalangium
Cyclodorippidae_Tymolus_brucei
Cyclodorippidae_Corycodus_disjunctipes
Cymonomidae_Cymonomus_diogenes
Dromiidae_Conchoecetes_artificiosus
Dromiidae_Lauridromia_dehaani
Dromiidae_Lewindromia_unidentata
Dynomenidae_Metadynomene_tanensis
Dynomenidae_Dynomene_praedator
Dynomenidae_Hirsutodynomene_ursula
Raninidae_Ranina_ranina
Lyreididae_Lyreidus_tridentatus
Lyreididae_Lysirude_griffini
Dotillidae_Dotilla_myctiroides
Dotillidae_Dotilla_wichmanni
Dotillidae_Tmethypocoelis_ceratophora
Dotillidae_Scopimera_bitympana
Dotillidae_Scopimera_intermedia
Dotillidae_Scopimera_proxima
Pinnotheridae_Nepinnotheres_cf_affinis
Pinnotheridae_Pinnixa_cylindrica
Pinnotheridae_Holothuriophilus_pacificus
Pinnotheridae_Pinnaxodes_chilensis
Xenograpsidae_Xenograpsus_testudinatus
Macrophthalmidae_Macrophthalmus_definitus
Macrophthalmidae_Macrophthalmus_erato
Macrophthalmidae_Chaenostoma_boscii
Varunidae_Gaetice_depressus
Varunidae_Hemigrapsus_penicillatus
Varunidae_Metaplax_longipes
Varunidae_Eriocheir_japonica
Varunidae_Cyclograpsus_cinereus
Mictyridae_Mictyris_brevidactylus
Sesarmidae_Nanosesarma_minutum
Sesarmidae_Parasesarma_affine
Sesarmidae_Perisesarma_bidens
Cryptochiridae_Dacryomaia_sp
Gecarcinidae_Gecarcoidea_lalandii
Gecarcinidae_Gecarcoidea_natalis
Plagusiidae_Plagusia_squamosa
Gecarcinidae_Cardisoma_crassum
Gecarcinidae_Tuerkayana_hirtipes
Glyptograpsidae_Glyptograpsus_jamaicensis
Glyptograpsidae_Platychirograpsus_spectabilis
Grapsidae_Grapsus_albolineatus
Grapsidae_Metopograpsus_frontalis
Grapsidae_Metopograpsus_quadridentatus
Percnidae_Percnon_planissimum
Percnidae_Percnon_affine
Percnidae_Percnon_affine_ourdata
Ocypodidae_Ocypode_ceratophthalmus
Ocypodidae_Ocypode_sinensis
Ocypodidae_Austruca_lactea
Ocypodidae_Gelasimus_borealis
Ocypodidae_Paraleptuca_crassipes
Hymenosomatidae_Hymenosoma_orbiculare
Pseudothelphusidae_Epilobocera_sinuatifrons
Pseudothelphusidae_Potamocarcinus_sp
Pseudothelphusidae_Ptychophallus_montanus
Pseudothelphusidae_Ptychophallus_sp
Retroplumidae_Retropluma_denticulata
Potamonautidae_Platythelphusa_armata
Potamonautidae_Potamonautes_perlatus
Potamonautidae_Potamonautes_subukia
Potamidae_Potamon_fluviatile
Potamidae_Potamon_persicum
Potamidae_Cryptopotamon_anacoluthon
Potamidae_Johora_tiomanensis
Potamidae_Isolapotamon_griswoldi
Gecarcinucidae_Somanniathelphusa_cf_zanklon
Gecarcinucidae_Somanniathelphusa_sp
Gecarcinucidae_Ceylonthelphusa_sp
Gecarcinucidae_Syntripsa_flavichela
Gecarcinucidae_Parathelphusa_pantherina
Gecarcinucidae_Parathelphusa_pantherina_ourdata
Orithyiidae_Orithyia_sinica
Trichodactylidae_Fredilocarcinus_apyratii
Trichodactylidae_Valdivia_serrata
Trichodactylidae_Trichodactylus_dentatus
Chasmocarcinidae_Megaesthesius_yokoyai
Belliidae_Acanthocyclus_albatrossis
Belliidae_Acanthocyclus_hassleri
Cancridae_Anatolikos_japonicus
Cancridae_Metacarcinus_magister
Leucosiidae_Pyrhila_carinata
Leucosiidae_Ebalia_tuberculosa
Leucosiidae_Praebebalia_longidactyla
Leucosiidae_Arcania_cornuta
Iphiculidae_Iphiculus_convexus
Leucosiidae_Tokoyo_eburnea
Leucosiidae_Urashima_pustuloides
Dairidae_Daira_americana
Pseudocarcinidae_Pseudocarcinus_gigas
Pseudocarcinidae_Pseudocarcinus_gigas_ourdata
Pilumnidae_Pilumnus_longicornis
Aethridae_Hepatus_epheliticus1and2
Carpiliidae_Carpilius_convexus
Carpiliidae_Carpilius_maculatus
Menippidae_Menippe_rumphii
Cheiragonidae_Telmessus_cheiragonus1and2
Corystidae_Gomeza_bicornis
Corystidae_Jonas_distinctus
Dorippidae_Dorippe_quadridens
Dorippidae_Heikeopsis_japonica
Dorippidae_Paradorippe_granulata
Calappidae_Acanthocarpus_alexandri
Calappidae_Calappa_philargius
Calappidae_Cycloes_granulata
Ethusidae_Ethusa_sexdentata
Matutidae_Matuta_planipes
Matutidae_Matuta_sp
Geryonidae_Chaceon_sp
Ovalipidae_Ovalipes_punctatus
Ovalipidae_Ovalipes_stephensoni
Polybiidae_Parathranites_orientalis
Thiidae_Thia_scutellata1and2
Portunidae_Portunus_pelagicus
Portunidae_Scylla_paramamosain
Portunidae_Charybdis_feriata
Portunidae_Thalamita_prymna
Oregoniidae_Chionoecetes_tanneri
Majidae_Maiopsis_panamensis
Epialtidae_Taliepus_dentatus
Epialtidae_Pugettia_nipponensis
Epialtidae_Pugettia_sp
Epialtidae_Pisoides_edwardsii
Epialtidae_Lubinia_dubia
Mithracidae_Teleophrys_ornatus
Mithracidae_Mithraculus_coryphe
Mithracidae_Mithraculus_forceps
Epialtidae_Hyastenus_sp
Epialtidae_Tiarinia_sp
Epialtidae_Doclea_canalifera
Epialtidae_Doclea_japonica
Parthenopidae_Ochtholambrus_excavatus
Parthenopidae_Cryptopodia_fornicata
Parthenopidae_Leiolambrus_punctatissimus
Parthenopidae_Parthenope_longimanus
Parthenopidae_Rhinolambrus_sp
Progeryonidae_Rhadinoplax_microptthalmus
Tanaochelidae_Tanaocheles_bidentata
Pilumnidae_Heteropanope_glabra
Galenidae_Galene_bispinosa
Pinnotheridae_Arcotheres_sp
Pilumnidae_Pilumnus_vespertilio
Pilumnidae_Danielum_ixbauchac
Mathildellidae_Mathildella_rubra
Oziidae_Epixanthus_frontalis
Oziidae_Epixanthus_sp
Oziidae_Lydia_annulipes
Oziidae_Ozius_reticulatus
Oziidae_Ozius_rugulosus
Eriphiidae_Eriphia_scabricula
Eriphiidae_Eriphia_smithii
Vultocinidae_Vultocinus_anfractus
Euryplacidae_Frevillea_barbata
Euryplacidae_Eucrate_crenata
Euryplacidae_Eucrate_alcocki
Euryplacidae_Eucrate_sp
Trapeziidae_Philippicarcinus_oviformis
Trapeziidae_Trapezia_septata
Trapeziidae_Trapezia_tigrina
Goneplacidae_Carcinoplax_longimana
Scalopidiidae_Scalopidia_spinosipes
Platyxanthidae_Homalaspis_plana
Goneplacidae_Carcinoplax_purpurea
Goneplacidae_Goneplax_sp
Xanthidae_Lybia_tessellata
Pseudorhombilidae_Lophoxanthus_lamellipes
Pseudorhombilidae_Pseudorhombila_quadridentata
Pseudorhombilidae_Trapezioplax_tridentata
Pseudorhombilidae_Nanoplax_xanthiformis
Pseudorhombilidae_Micropanope_sculptipes
Panopeidae_Panopeus_herbstii
Panopeidae_Glyptoplax_smithii
Xanthidae_Carpoporus_papulosus
Xanthidae_Xantho_pilipes
Xanthidae_Palapedia_cf_nitida
Xanthidae_Atergatis_floridus
Xanthidae_Atergatis_integerrimus
Xanthidae_Paraxanthus_barbiger
Xanthidae_Xanthodius_sternberghii1and2
Xanthidae_Platyactaea_dovii
Xanthidae_Actaea_bifrons
Xanthidae_Demania_scaberrima
Xanthidae_Actaeodes_cf_tomentosus
Xanthidae_Novactaea_bella
0.04

#### Fig. S8

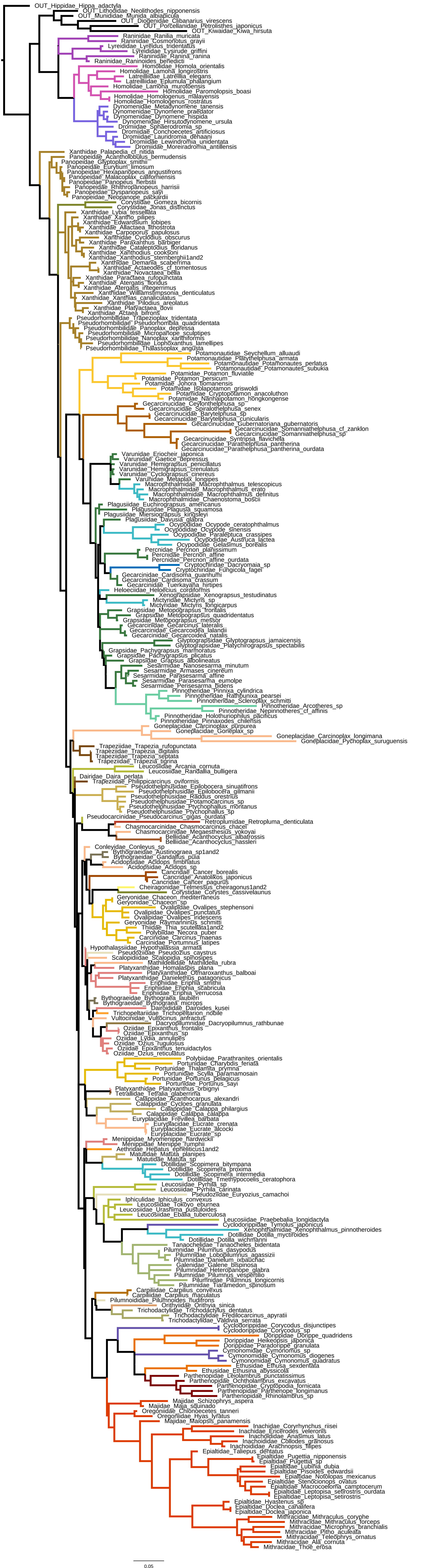

OUT_Hippidae_Hippa_adactyla
OUT_Lithodidae_Neolithodes_nipponensis
OUT_Munididae_Munida_albiapicula
OUT_Diogenidae_Clibanarius_virescens
OUT_Porcellanidae_Petrolisthes_japonicus
OUT_Kiwaidae_Kiwa_hirsuta
Raninidae_Ranilia_muricata
Raninidae_Cosmonotus_grayii
Lyreididae_Lyreidus_tridentatus
Lyreididae_Lysirude_griffini
Raninidae_Ranina_ranina
Raninidae_Raninoides_benedicti
Homolidae_Homola_orientalis
Homolidae_Lamoha_longirostris
Latreilliidae_Latreillia_elegans
Latreilliidae_Eplumula_phalangium
Homolidae_Lamoha_murotoensis
Homolidae_Paromolopsis_boasi
Homolidae_Homologenus_malayensis
Homolidae_Homologenus_rostratus
Dynomenidae_Metadynomene_tanensis
Dynomenidae_Dynomene_praedator
Dynomenidae_Dynomene_hispida
Dynomenidae_Hirsutodynomene_ursula
Dromiidae_Sphaerodromia_sp
Dromiidae_Conchoecetes_artificiosus
Dromiidae_Lauridromia_dehaani
Dromiidae_Lewindromia_unidentata
Dromiidae_Moreiradromia_antillensis
Xanthidae_Palapedia_cf_nitida
Panopeidae_Acantholobulus_bermudensis
Panopeidae_Glyptoplax_smithii
Panopeidae_Eurytium_limosum
Panopeidae_Hexapanopeus_angustifrons
Panopeidae_Malacoplax_californiensis
Panopeidae_Panopeus_herbstii
Panopeidae_Rhithropanopeus_harrisii
Panopeidae_Dyspanopeus_sayi
Panopeidae_Neopanope_packardii
Corystidae_Gomeza_bicornis
Corystidae_Jonas_distinctus
Xanthidae_Lybia_tessellata
Xanthidae_Xantho_pilipes
Xanthidae_Edwardsium_lobipes
Xanthidae_Allactaea_lithostrota
Xanthidae_Carpoporus_papulosus
Xanthidae_Cyclodius_obscurus
Xanthidae_Paraxanthus_barbiger
Xanthidae_Cataleptodius_floridanus
Xanthidae_Xanthodius_cooksoni
Xanthidae_Xanthodius_sternberghii1and2
Xanthidae_Demania_scaberrima
Xanthidae_Actaeodes_cf_tomentosus
Xanthidae_Novactaea_bella
Xanthidae_Paractaea_rufopunctata
Xanthidae_Atergatis_floridus
Xanthidae_Atergatis_integerrimus
Xanthidae_Williamstimpsonia_denticulatus
Xanthidae_Xanthias_canaliculatus
Xanthidae_Pilodius_areolatus
Xanthidae_Platyactaea_dovii
Xanthidae_Actaea_bifrons
Pseudorhombilidae_Trapezioplax_tridentata
Pseudorhombilidae_Pseudorhombila_quadridentata
Pseudorhombilidae_Panoplax_depressa
Pseudorhombilidae_Micropanope_sculptipes
Pseudorhombilidae_Nanoplax_xanthiformis
Pseudorhombilidae_Lophoxanthus_lamellipes
Pseudorhombilidae_Thalassoplax_angusta
Potamonautidae_Seychellum_alluaudi
Potamonautidae_Platythelphusa_armata
Potamonautidae_Potamonautes_perlatus
Potamonautidae_Potamonautes_subukia
Potamidae_Potamon_fluviatile
Potamidae_Potamon_persicum
Potamidae_Johora_tiomanensis
Potamidae_Isolapotamon_griswoldi
Potamidae_Cryptopotamon_anacoluthon
Potamidae_Nanhaipotamon_hongkongense
Gecarcinucidae_Ceylonthelphusa_sp
Gecarcinucidae_Spiralothelphusa_senex
Gecarcinucidae_Barytelphusa_sp
Gecarcinucidae_Barytelphusa_cunicularis
Gecarcinucidae_Gubernatoriana_gubernatoris
Gecarcinucidae_Somanniathelphusa_cf_zanklon
Gecarcinucidae_Somanniathelphusa_sp
Gecarcinucidae_Syntripsa_flavichela
Gecarcinucidae_Parathelphusa_pantherina
Gecarcinucidae_Parathelphusa_pantherina_ourdata
Varunidae_Eriocheir_japonica
Varunidae_Gaetice_depressus
Varunidae_Hemigrapsus_penicillatus
Varunidae_Hemigrapsus_crenulatus
Varunidae_Cyclograpsus_cinereus
Varunidae_Metaplax_longipes
Macrophthalmidae_Macrophthalmus_telescopicus
Macrophthalmidae_Macrophthalmus_erato
Macrophthalmidae_Macrophthalmus_definitus
Macrophthalmidae_Chaenostoma_boscii
Plagusiidae_Euchirograpsus_americanus
Plagusiidae_Plagusia_squamosa
Plagusiidae_Miersiograpsus_kingsleyi
Plagusiidae_Davusia_glabra
Ocypodidae_Ocypode_ceratophthalmus
Ocypodidae_Ocypode_sinensis
Ocypodidae_Paraleptuca_crassipes
Ocypodidae_Austruca_lactea
Ocypodidae_Gelasimus_borealis
Percnidae_Percnon_planissimum
Percnidae_Percnon_affine
Percnidae_Percnon_affine_ourdata
Cryptochiridae_Dacryomaia_sp
Cryptochiridae_Fungicola_fagei
Gecarcinidae_Cardisoma_guanhumi
Gecarcinidae_Cardisoma_crassum
Gecarcinidae_Tuerkayana_hirtipes
Heloeciidae_Heloecius_cordiformis
Xenograpsidae_Xenograpsus_testudinatus
Mictyridae_Mictyris_sp
Mictyridae_Mictyris_longicarpus
Grapsidae_Metopograpsus_frontalis
Grapsidae_Metopograpsus_quadridentatus
Grapsidae_Metopograpsus_messor
Gecarcinidae_Gecarcinus_lateralis
Gecarcinidae_Gecarcoidea_lalandii
Gecarcinidae_Gecarcoidea_natalis
Glyptograpsidae_Glyptograpsus_jamaicensis
Glyptograpsidae_Platychirograpsus_spectabilis
Grapsidae_Pachygrapsus_marmoratus
Grapsidae_Pachygrapsus_plicatus
Grapsidae_Grapsus_albolineatus
Sesarmidae_Nanosesarma_minutum
Sesarmidae_Armases_cinereum
Sesarmidae_Parasesarma_affine
Sesarmidae_Parasesarma_eumolpe
Sesarmidae_Perisesarma_bidens
Pinnotheridae_Pinnixa_cylindrica
Pinnotheridae_Rathbunixa_pearsei
Pinnotheridae_Scleroplax_schmitti
Pinnotheridae_Arcotheres_sp
Pinnotheridae_Nepinnotheres_cf_affinis
Pinnotheridae_Holothuriophilus_pacificus
Pinnotheridae_Pinnaxodes_chilensis
Goneplacidae_Carcinoplax_purpurea
Goneplacidae_Goneplax_sp
Goneplacidae_Carcinoplax_longimana
Goneplacidae_Pycnoplax_suruguensis
Trapeziidae_Trapezia_rufopunctata
Trapeziidae_Trapezia_digitalis
Trapeziidae_Trapezia_septata
Trapeziidae_Trapezia_tigrina
Leucosiidae_Arcania_cornuta
Leucosiidae_Randallia_bulligera
Dairidae_Daira_perlata
Trapeziidae_Philippicarcinus_oviformis
Pseudothelphusidae_Epilobocera_sinuatifrons
Pseudothelphusidae_Epilobocera_gilmanii
Pseudothelphusidae_Raddus_orestrius
Pseudothelphusidae_Potamocarcinus_sp
Pseudothelphusidae_Ptychophallus_montanus
Pseudothelphusidae_Ptychophallus_sp
Pseudocarcinidae_Pseudocarcinus_gigas_ourdata
Retroplumidae_Retropluma_denticulata
Chasmocarcinidae_Chasmocarcinus_chacei
Chasmocarcinidae_Megaesthesius_yokoyai
Belliidae_Acanthocyclus_albatrossis
Belliidae_Acanthocyclus_hassleri
Conleyidae_Conleyus_sp
Bythograeidae_Austinograea_sp1and2
Bythograeidae_Gandalfus_puia
Acidopsidae_Acidops_fimbriatus
Acidopsidae_Acidops_sp
Cancridae_Cancer_borealis
Cancridae_Anatolikos_japonicus
Cancridae_Cancer_pagurus
Cheiragonidae_Telmessus_cheiragonus1and2
Corystidae_Corystes_cassivelaunus
Geryonidae_Chaceon_mediterraneus
Geryonidae_Chaceon_sp
Ovalipidae_Ovalipes_stephensoni
Ovalipidae_Ovalipes_punctatus
Ovalipidae_Ovalipes_iridescens
Geryonidae_Raymanninus_schmitti
Thiidae_Thia_scutellata1and2
Polybiidae_Necora_puber
Carcinidae_Carcinus_maenas
Carcinidae_Portumnus_latipes
Hypothalassiidae_Hypothalassia_armata
Pseudoziidae_Pseudozius_caystrus
Scalopidiidae_Scalopidia_spinosipes
Mathildellidae_Mathildella_rubra
Platyxanthidae_Homalaspis_plana
Platyxanthidae_Otmaroxanthus_balboai
Platyxanthidae_Danielethus_patagonicus
Eriphiidae_Eriphia_smithii
Eriphiidae_Eriphia_scabricula
Eriphiidae_Eriphia_verrucosa
Bythograeidae_Bythograea_laubieri
Bythograeidae_Bythograea_microps
Dairoididae_Dairoides_kusei
Trichopeltariidae_Trichopeltarion_nobile
Vultocinidae_Vultocinus_anfractus
Dacryopilumnidae_Dacryopilumnus_rathbunae
Oziidae_Epixanthus_frontalis
Oziidae_Epixanthus_sp
Oziidae_Lydia_annulipes
Oziidae_Ozius_rugulosus
Oziidae_Epixanthus_tenuidactylos
Oziidae_Ozius_reticulatus
Polybiidae_Parathranites_orientalis
Portunidae_Charybdis_feriata
Portunidae_Thalamita_prymna
Portunidae_Scylla_paramamosain
Portunidae_Portunus_pelagicus
Portunidae_Portunus_sayi
Platyxanthidae_Platyxanthus_orbignyi
Tetraliidae_Tetralia_glaberrima
Calappidae_Acanthocarpus_alexandri
Calappidae_Cycloes_granulata
Calappidae_Calappa_philargius
Calappidae_Calappa_calappa
Euryplacidae_Frevillea_barbata
Euryplacidae_Eucrate_crenata
Euryplacidae_Eucrate_alcocki
Euryplacidae_Eucrate_sp
Menippidae_Myomenippe_hardwickii
Menippidae_Menippe_rumphii
Aethridae_Hepatus_epheliticus1and2
Matutidae_Matuta_planipes
Matutidae_Matuta_sp
Dotillidae_Scopimera_bitympana
Dotillidae_Scopimera_proxima
Dotillidae_Scopimera_intermedia
Dotillidae_Tmethypocoelis_ceratophora
Leucosiidae_Pyrhila_sp
Leucosiidae_Pyrhila_carinata
Pseudoziidae_Euryozius_camachoi
Iphiculidae_Iphiculus_convexus
Leucosiidae_Tokoyo_eburnea
Leucosiidae_Urashima_pustuloides
Leucosiidae_Ebalia_tuberculosa
Leucosiidae_Praebebalia_longidactyla
Cyclodorippidae_Tymolus_japonicus
Xenophthalmidae_Xenophthalmus_pinnotheroides
Dotillidae_Dotilla_myctiroides
Dotillidae_Dotilla_wichmanni
Tanaochelidae_Tanaocheles_bidentata
Pilumnidae_Pilumnus_dasypodus
Pilumnidae_Lobopilumnus_agassizii
Pilumnidae_Danielum_ixbauchac
Galenidae_Galene_bispinosa
Pilumnidae_Heteropanope_glabra
Pilumnidae_Pilumnus_vespertilio
Pilumnidae_Pilumnus_longicornis
Pilumnidae_Tiaramedon_spinosum
Carpiliidae_Carpilius_convexus
Carpiliidae_Carpilius_maculatus
Pilumnoididae_Pilumnoides_nudifrons
Orithyiidae_Orithyia_sinica
Trichodactylidae_Trichodactylus_dentatus
Trichodactylidae_Fredilocarcinus_apyratii
Trichodactylidae_Valdivia_serrata
Cyclodorippidae_Corycodus_disjunctipes
Cyclodorippidae_Corycodus_sp
Dorippidae_Dorippe_quadridens
Dorippidae_Heikeopsis_japonica
Dorippidae_Paradorippe_granulata
Cymonomidae_Cymonomus_sp
Cymonomidae_Cymonomus_diogenes
Cymonomidae_Cymonomus_quadratus
Ethusidae_Ethusa_sexdentata
Ethusidae_Ethusina_abyssicola
Parthenopidae_Leiolambrus_punctatissimus
Parthenopidae_Ochtholambrus_excavatus
Parthenopidae_Cryptopodia_fornicata
Parthenopidae_Parthenope_longimanus
Parthenopidae_Rhinolambrus_sp
Majidae_Schizophrys_aspera
Majidae_Maja_squinado
Oregoniidae_Chionoecetes_tanneri
Oregoniidae_Hyas_lyratus
Majidae_Maiopsis_panamensis
Inachidae_Coryrhynchus_riisei
Inachidae_Ericerodes_veleronis
Inachoididae_Anasimus_latus
Inachoididae_Collodes_granosus
Inachoididae_Arachnopsis_filipes
Epialtidae_Taliepus_dentatus
Epialtidae_Pugettia_nipponensis
Epialtidae_Pugettia_sp
Epialtidae_Lubinia_dubia
Epialtidae_Pisoides_edwardsii
Epialtidae_Notolopas_mexicanus
Epialtidae_Stenocionops_ovatus
Epialtidae_Macrocoeloma_camptocerum
Epialtidae_Leptopisa_setirostris_ourdata
Epialtidae_Leptopisa_setirostris
Epialtidae_Hyastenus_sp
Epialtidae_Doclea_canalifera
Epialtidae_Doclea_japonica
Mithracidae_Mithraculus_coryphe
Mithracidae_Mithraculus_forceps
Mithracidae_Microphrys_branchialis
Mithracidae_Pitho_aculeata
Mithracidae_Teleophrys_ornatus
Mithracidae_Ala_cornuta
Mithracidae_Thoe_erosa
0.05

Fig. S9

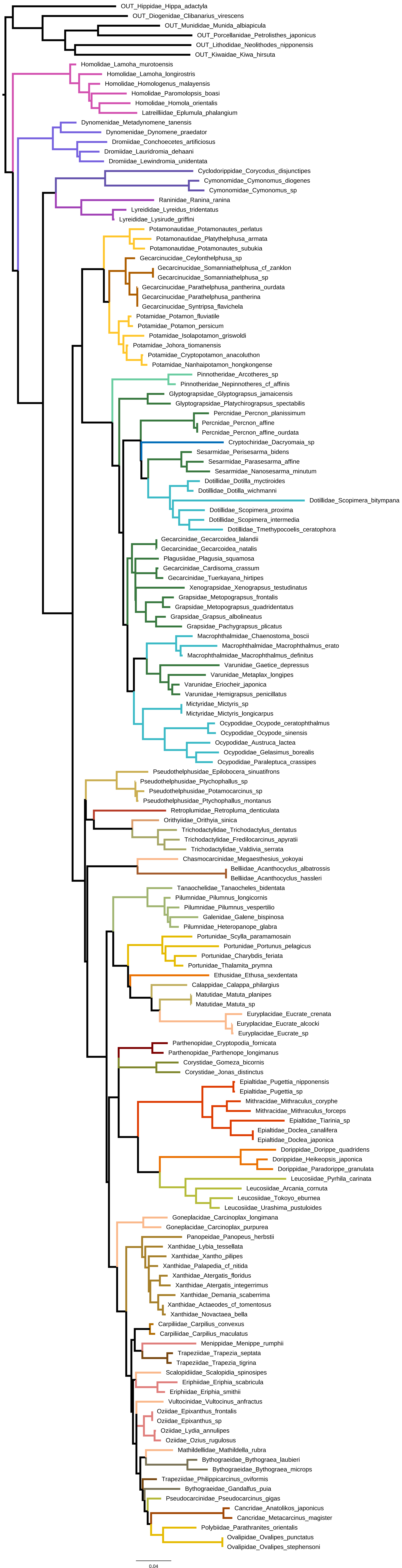
OUT_Hippidae_Hippa_adactyla
OUT_Diogenidae_Clibanarius_virescens
OUT_Munididae_Munida_albiapicula
OUT_Porcellanidae_Petrolisthes_japonicus
OUT_Lithodidae_Neolithodes_nipponensis
OUT_Kiwaidae_Kiwa_hirsuta
Homolidae_Lamoha_murotoensis
Homolidae_Lamoha_longirostris
Homolidae_Homologenus_malayensis
Homolidae_Paromolopsis_boasi
Homolidae_Homola_orientalis
Latreilliidae_Eplumula_phalangium
Dynomenidae_Metadynomene_tanensis
Dynomenidae_Dynomene_praedator
Dromiidae_Conchoecetes_artificiosus
Dromiidae_Lauridromia_dehaani
Dromiidae_Lewindromia_unidentata
Cyclodorippidae_Corycodus_disjunctipes
Cymonomidae_Cymonomus_diogenes
Cymonomidae_Cymonomus_sp
Raninidae_Ranina_ranina
Lyreididae_Lyreidus_tridentatus
Lyreididae_Lysirude_griffini
Potamonautidae_Potamonautes_perlatus
Potamonautidae_Platythelphusa_armata
Potamonautidae_Potamonautes_subukia
Gecarcinucidae_Ceylonthelphusa_sp
Gecarcinucidae_Somanniathelphusa_cf_zanklon
Gecarcinucidae_Somanniathelphusa_sp
Gecarcinucidae_Parathelphusa_pantherina_ourdata
Gecarcinucidae_Parathelphusa_pantherina
Gecarcinucidae_Syntripsa_flavichela
Potamidae_Potamon_fluviatile
Potamidae_Potamon_persicum
Potamidae_Isolapotamon_griswoldi
Potamidae_Johora_tiomanensis
Potamidae_Cryptopotamon_anacoluthon
Potamidae_Nanhaipotamon_hongkongense
Pinnotheridae_Arcotheres_sp
Pinnotheridae_Nepinnotheres_cf_affinis
Glyptograpsidae_Glyptograpsus_jamaicensis
Glyptograpsidae_Platychirograpsus_spectabilis
Percnidae_Percnon_planissimum
Percnidae_Percnon_affine
Percnidae_Percnon_affine_ourdata
Cryptochiridae_Dacryomaia_sp
Sesarmidae_Perisesarma_bidens
Sesarmidae_Parasesarma_affine
Sesarmidae_Nanosesarma_minutum
Dotillidae_Dotilla_myctiroides
Dotillidae_Dotilla_wichmanni
Dotillidae_Scopimera_bitympana
Dotillidae_Scopimera_proxima
Dotillidae_Scopimera_intermedia
Dotillidae_Tmethypocoelis_ceratophora
Gecarcinidae_Gecarcoidea_lalandii
Gecarcinidae_Gecarcoidea_natalis
Plagusiidae_Plagusia_squamosa
Gecarcinidae_Cardisoma_crassum
Gecarcinidae_Tuerkayana_hirtipes
Xenograpsidae_Xenograpsus_testudinatus
Grapsidae_Metopograpsus_frontalis
Grapsidae_Metopograpsus_quadridentatus
Grapsidae_Grapsus_albolineatus
Grapsidae_Pachygrapsus_plicatus
Macrophthalmidae_Chaenostoma_boscii
Macrophthalmidae_Macrophthalmus_erato
Macrophthalmidae_Macrophthalmus_definitus
Varunidae_Gaetice_depressus
Varunidae_Metaplax_longipes
Varunidae_Eriocheir_japonica
Varunidae_Hemigrapsus_penicillatus
Mictyridae_Mictyris_sp
Mictyridae_Mictyris_longicarpus
Ocypodidae_Ocypode_ceratophthalmus
Ocypodidae_Ocypode_sinensis
Ocypodidae_Austruca_lactea
Ocypodidae_Gelasimus_borealis
Ocypodidae_Paraleptuca_crassipes
Pseudothelphusidae_Epilobocera_sinuatifrons
Pseudothelphusidae_Ptychophallus_sp
Pseudothelphusidae_Potamocarcinus_sp
Pseudothelphusidae_Ptychophallus_montanus
Retroplumidae_Retropluma_denticulata
Orithyiidae_Orithyia_sinica
Trichodactylidae_Trichodactylus_dentatus
Trichodactylidae_Fredilocarcinus_apyratii
Trichodactylidae_Valdivia_serrata
Chasmocarcinidae_Megaesthesius_yokoyai
Belliidae_Acanthocyclus_albatrossis
Belliidae_Acanthocyclus_hassleri
Tanaocheliidae_Tanaocheles_bidentata
Pilumnidae_Pilumnus_longicornis
Pilumnidae_Pilumnus_vespertilio
Galenidae_Galene_bispinosa
Pilumnidae_Heteropanope_glabra
Portunidae_Scylla_paramamosain
Portunidae_Portunus_pelagicus
Portunidae_Charybdis_feriata
Portunidae_Thalamita_prymna
Ethusidae_Ethusa_sexdentata
Calappidae_Calappa_philargius
Matutidae_Matuta_planipes
Matutidae_Matuta_sp
Euryplacidae_Eucrate_crenata
Euryplacidae_Eucrate_alcocki
Euryplacidae_Eucrate_sp
Parthenopidae_Cryptopodia_fornicata
Parthenopidae_Parthenope_longimanus
Corystidae_Gomeza_bicornis
Corystidae_Jonas_distinctus
Epialtidae_Pugettia_nipponensis
Epialtidae_Pugettia_sp
Mithracidae_Mithraculus_coryphe
Mithracidae_Mithraculus_forceps
Epialtidae_Tiarinia_sp
Epialtidae_Doclea_canalifera
Epialtidae_Doclea_japonica
Dorippidae_Dorippe_quadridens
Dorippidae_Heikeopsis_japonica
Dorippidae_Paradorippe_granulata
Leucosiidae_Pyrhila_carinata
Leucosiidae_Arcania_cornuta
Leucosiidae_Tokoyo_eburnea
Leucosiidae_Urashima_pustuloides
Goneplacidae_Carcinoplax_longimana
Goneplacidae_Carcinoplax_purpurea
Panopeidae_Panopeus_herbstii
Xanthidae_Lybia_tessellata
Xanthidae_Xantho_pilipes
Xanthidae_Palapedia_cf_nitida
Xanthidae_Atergatis_floridus
Xanthidae_Atergatis_integerrimus
Xanthidae_Demania_scaberrima
Xanthidae_Actaeodes_cf_tomentosus
Xanthidae_Novactaea_bella
Carpiliidae_Carpilius_convexus
Carpiliidae_Carpilius_maculatus
Menippidae_Menippe_rumphii
Trapeziidae_Trapezia_septata
Trapeziidae_Trapezia_tigrina
Scalopidiidae_Scalopidia_spinosipes
Eriphiidae_Eriphia_scabricula
Eriphiidae_Eriphia_smithii
Vultocinidae_Vultocinus_anfractus
Oziidae_Epixanthus_frontalis
Oziidae_Epixanthus_sp
Oziidae_Lydia_annulipes
Oziidae_Ozius_rugulosus
Mathildellidae_Mathildella_rubra
Bythograeidae_Bythograea_laubieri
Bythograeidae_Bythograea_microps
Trapeziidae_Philippicarcinus_oviformis
Bythograeidae_Gandalfus_puia
Pseudocarcinidae_Pseudocarcinus_gigas
Cancridae_Anatolikos_japonicus
Cancridae_Metacarcinus_magister
Polybiidae_Parathranites_orientalis
Ovalipidae_Ovalipes_punctatus
Ovalipidae_Ovalipes_stephensoni
0.04

**Fig. S10**

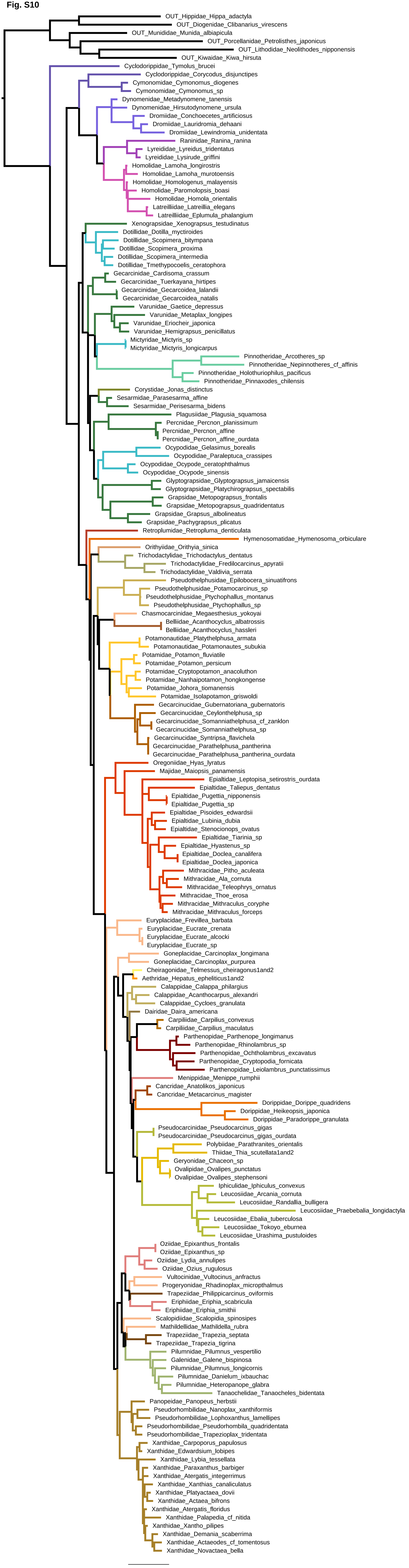

OUT_Hippidae_Hippa_adactyla
OUT_Diogenidae_Clibanarius_virescens
OUT_Munididae_Munida_albiapicula
OUT_Porcellanidae_Petrolisthes_japonicus
OUT_Lithodidae_Neolithodes_nipponensis
OUT_Kiwaidae_Kiwa_hirsuta
Cyclodorippidae_Tymolus_brucei
Cyclodorippidae_Corycodus_disjunctipes
Cymonomidae_Cymonomus_diogenes
Cymonomidae_Cymonomus_sp
Dynomenidae_Metadynomene_tanensis
Dynomenidae_Hirsutodynomene_ursula
Dromiidae_Conchoecetes_artificiosus
Dromiidae_Lauridromia_dehaani
Dromiidae_Lewindromia_unidentata
Raninidae_Ranina_ranina
Lyreididae_Lyreidus_tridentatus
Lyreididae_Lysirude_griffini
Homolidae_Lamoha_longirostris
Homolidae_Lamoha_murotoensis
Homolidae_Homologenus_malayensis
Homolidae_Paromolopsis_boasi
Homolidae_Homola_orientalis
Latreilliidae_Latreillia_elegans
Latreilliidae_Eplumula_phalangium
Xenograpsidae_Xenograpsus_testudinatus
Dotillidae_Dotilla_myctiroides
Dotillidae_Scopimera_bitympana
Dotillidae_Scopimera_proxima
Dotillidae_Scopimera_intermedia
Dotillidae_Tmethypocoelis_ceratophora
Gecarcinidae_Cardisoma_crassum
Gecarcinidae_Tuerkayana_hirtipes
Gecarcinidae_Gecarcoidea_lalandii
Gecarcinidae_Gecarcoidea_natalis
Varunidae_Gaetice_depressus
Varunidae_Metaplax_longipes
Varunidae_Eriocheir_japonica
Varunidae_Hemigrapsus_penicillatus
Mictyridae_Mictyris_sp
Mictyridae_Mictyris_longicarpus
Pinnotheridae_Arcotheres_sp
Pinnotheridae_Nepinnotheres_cf_affinis
Pinnotheridae_Holothuriophilus_pacificus
Pinnotheridae_Pinnaxodes_chilensis
Corystidae_Jonas_distinctus
Sesarmidae_Parasesarma_affine
Sesarmidae_Perisesarma_bidens
Plagusiidae_Plagusia_squamosa
Percnidae_Percnon_planissimum
Percnidae_Percnon_affine
Percnidae_Percnon_affine_ourdata
Ocypodidae_Gelasimus_borealis
Ocypodidae_Paraleptuca_crassipes
Ocypodidae_Ocypode_ceratophthalmus
Ocypodidae_Ocypode_sinensis
Glyptograpsidae_Glyptograpsus_jamaicensis
Glyptograpsidae_Platychirograpsus_spectabilis
Grapsidae_Metopograpsus_frontalis
Grapsidae_Metopograpsus_quadridentatus
Grapsidae_Grapsus_albolineatus
Grapsidae_Pachygrapsus_plicatus
Retroplumidae_Retropluma_denticulata
Hymenosomatidae_Hymenosoma_orbiculare
Orithyiidae_Orithyia_sinica
Trichodactylidae_Trichodactylus_dentatus
Trichodactylidae_Fredilocarcinus_apyratii
Trichodactylidae_Valdivia_serrata
Pseudothelphusidae_Epilobocera_sinuatifrons
Pseudothelphusidae_Potamocarcinus_sp
Pseudothelphusidae_Ptychophallus_montanus
Pseudothelphusidae_Ptychophallus_sp
Chasmocarcinidae_Megaesthesius_yokoyai
Belliidae_Acanthocyclus_albatrossis
Belliidae_Acanthocyclus_hassleri
Potamonautidae_Platythelphusa_armata
Potamonautidae_Potamonautes_subukia
Potamidae_Potamon_fluviatile
Potamidae_Potamon_persicum
Potamidae_Cryptopotamon_anacoluthon
Potamidae_Nanhaipotamon_hongkongense
Potamidae_Johora_tiomanensis
Potamidae_Isolapotamon_griswoldi
Gecarcinucidae_Gubernatoriana_gubernatoris
Gecarcinucidae_Ceylonthelphusa_sp
Gecarcinucidae_Somanniathelphusa_cf_zanklon
Gecarcinucidae_Somanniathelphusa_sp
Gecarcinucidae_Syntripsa_flavichela
Gecarcinucidae_Parathelphusa_pantherina
Gecarcinucidae_Parathelphusa_pantherina_ourdata
Oregoniidae_Hyas_lyratus
Majidae_Maiopsis_panamensis
Epialtidae_Leptopisa_setirostris_ourdata
Epialtidae_Taliepus_dentatus
Epialtidae_Pugettia_nipponensis
Epialtidae_Pugettia_sp
Epialtidae_Pisoides_edwardsii
Epialtidae_Lubinia_dubia
Epialtidae_Stenocionops_ovatus
Epialtidae_Tiarinia_sp
Epialtidae_Hyastenus_sp
Epialtidae_Doclea_canalifera
Epialtidae_Doclea_japonica
Mithracidae_Pitho_aculeata
Mithracidae_Ala_cornuta
Mithracidae_Teleophrys_ornatus
Mithracidae_Thoe_erosa
Mithracidae_Mithraculus_coryphe
Mithracidae_Mithraculus_forceps
Euryplacidae_Frevillea_barbata
Euryplacidae_Eucrate_crenata
Euryplacidae_Eucrate_alcocki
Euryplacidae_Eucrate_sp
Goneplacidae_Carcinoplax_longimana
Goneplacidae_Carcinoplax_purpurea
Cheiragonidae_Telmessus_cheiragonus1and2
Aethridae_Hepatus_epheliticus1and2
Calappidae_Calappa_philargius
Calappidae_Acanthocarpus_alexandri
Calappidae_Cycloes_granulata
Dairidae_Daira_americana
Carpiliidae_Carpilius_convexus
Carpiliidae_Carpilius_maculatus
Parthenopidae_Parthenope_longimanus
Parthenopidae_Rhinolambrus_sp
Parthenopidae_Ochtholambrus_excavatus
Parthenopidae_Cryptopodia_fornicata
Parthenopidae_Leiolambrus_punctatissimus
Menippidae_Menippe_rumphii
Cancridae_Anatolikos_japonicus
Cancridae_Metacarcinus_magister
Dorippidae_Dorippe_quadridens
Dorippidae_Heikeopsis_japonica
Dorippidae_Paradorippe_granulata
Pseudocarcinidae_Pseudocarcinus_gigas
Pseudocarcinidae_Pseudocarcinus_gigas_ourdata
Polybiidae_Parathranites_orientalis
Thiidae_Thia_scutellata1and2
Geryonidae_Chaceon_sp
Ovalipidae_Ovalipes_punctatus
Ovalipidae_Ovalipes_stephensoni
Iphiculidae_Iphiculus_convexus
Leucosiidae_Arcania_cornuta
Leucosiidae_Randallia_bulligera
Leucosiidae_Praebebalia_longidactyla
Leucosiidae_Ebalia_tuberculosa
Leucosiidae_Tokoyo_eburnea
Leucosiidae_Urashima_pustuloides
Oziidae_Epixanthus_frontalis
Oziidae_Epixanthus_sp
Oziidae_Lydia_annulipes
Oziidae_Ozius_rugulosus
Vultocinidae_Vultocinus_anfractus
Progeryonidae_Rhadinoplax_micropthalmus
Trapeziidae_Philippicarcinus_oviformis
Eriphiidae_Eriphia_scabricula
Eriphiidae_Eriphia_smithii
Scalopidiidae_Scalopidia_spinosipes
Mathildellidae_Mathildella_rubra
Trapeziidae_Trapezia_septata
Trapeziidae_Trapezia_tigrina
Pilumnidae_Pilumnus_vespertilio
Galenidae_Galene_bispinosa
Pilumnidae_Pilumnus_longicornis
Pilumnidae_Danielum_ixbauchac
Pilumnidae_Heteropanope_glabra
Tanaochelidae_Tanaocheles_bidentata
Panopeidae_Panopeus_herbstii
Pseudorhombilidae_Nanoplax_xanthiformis
Pseudorhombilidae_Lophoxanthus_lamellipes
Pseudorhombilidae_Pseudorhombila_quadridentata
Pseudorhombilidae_Trapezioplax_tridentata
Xanthidae_Carpoporus_papulosus
Xanthidae_Edwardsium_lobipes
Xanthidae_Lybia_tessellata
Xanthidae_Paraxanthus_barbiger
Xanthidae_Atergatis_integerrimus
Xanthidae_Xanthias_canaliculatus
Xanthidae_Platyactaea_dovii
Xanthidae_Actaea_bifrons
Xanthidae_Atergatis_floridus
Xanthidae_Palapedia_cf_nitida
Xanthidae_Xantho_pilipes
Xanthidae_Demania_scaberrima
Xanthidae_Actaeodes_cf_tomentosus
Xanthidae_Novactaea_bella
0.06

Fig. S11

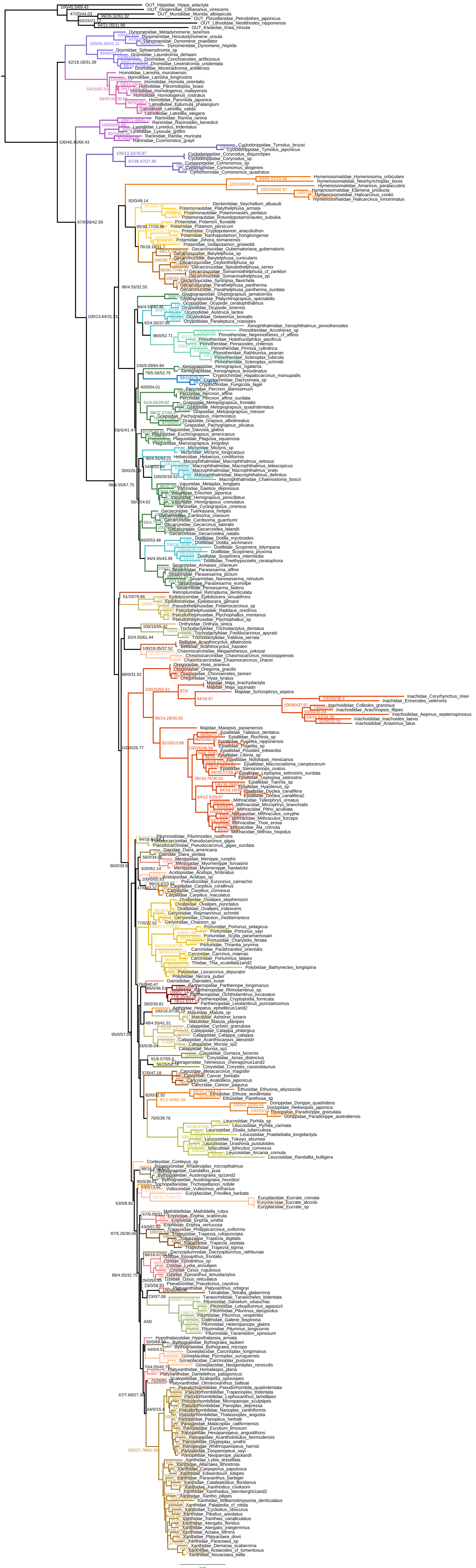

**Fig. S12**

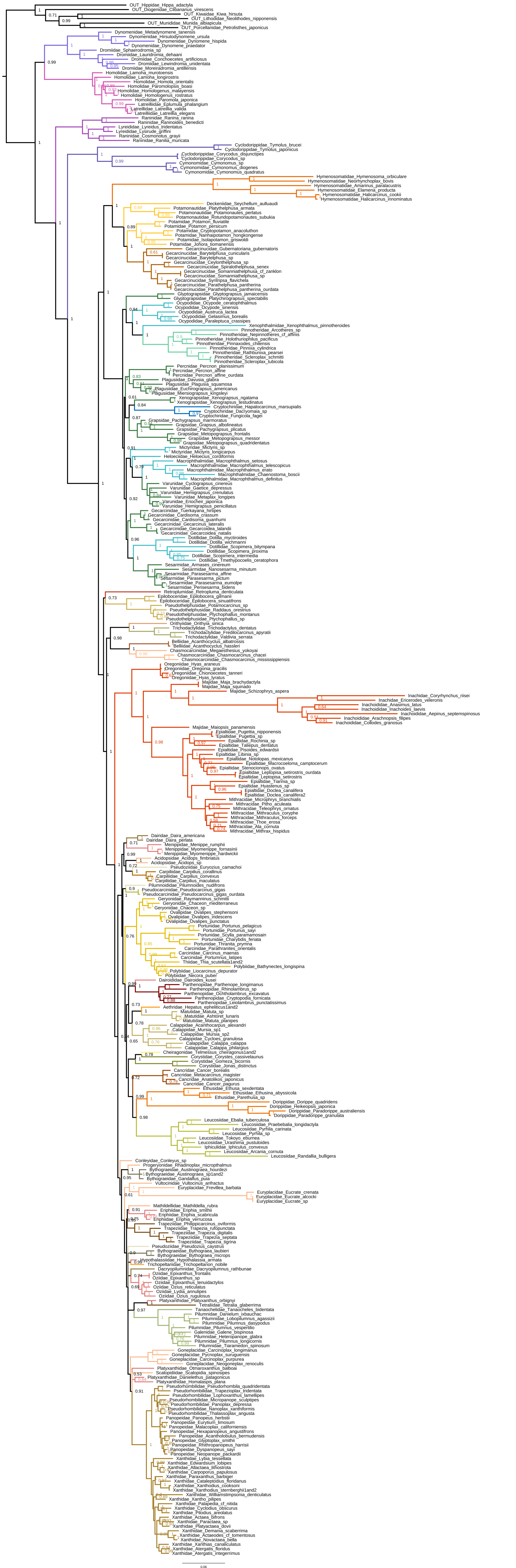

**Fig. S13**

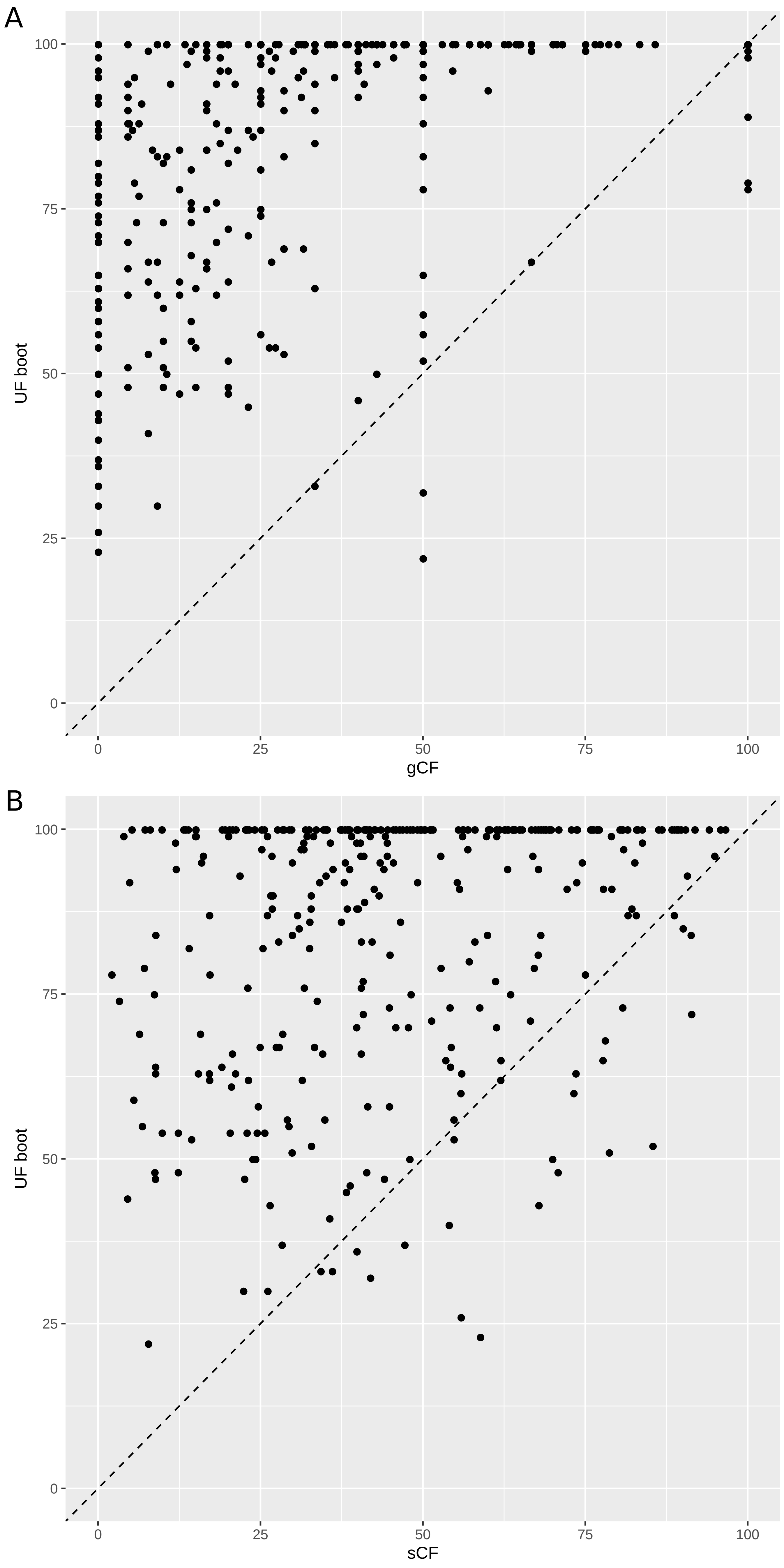

Fig. S14

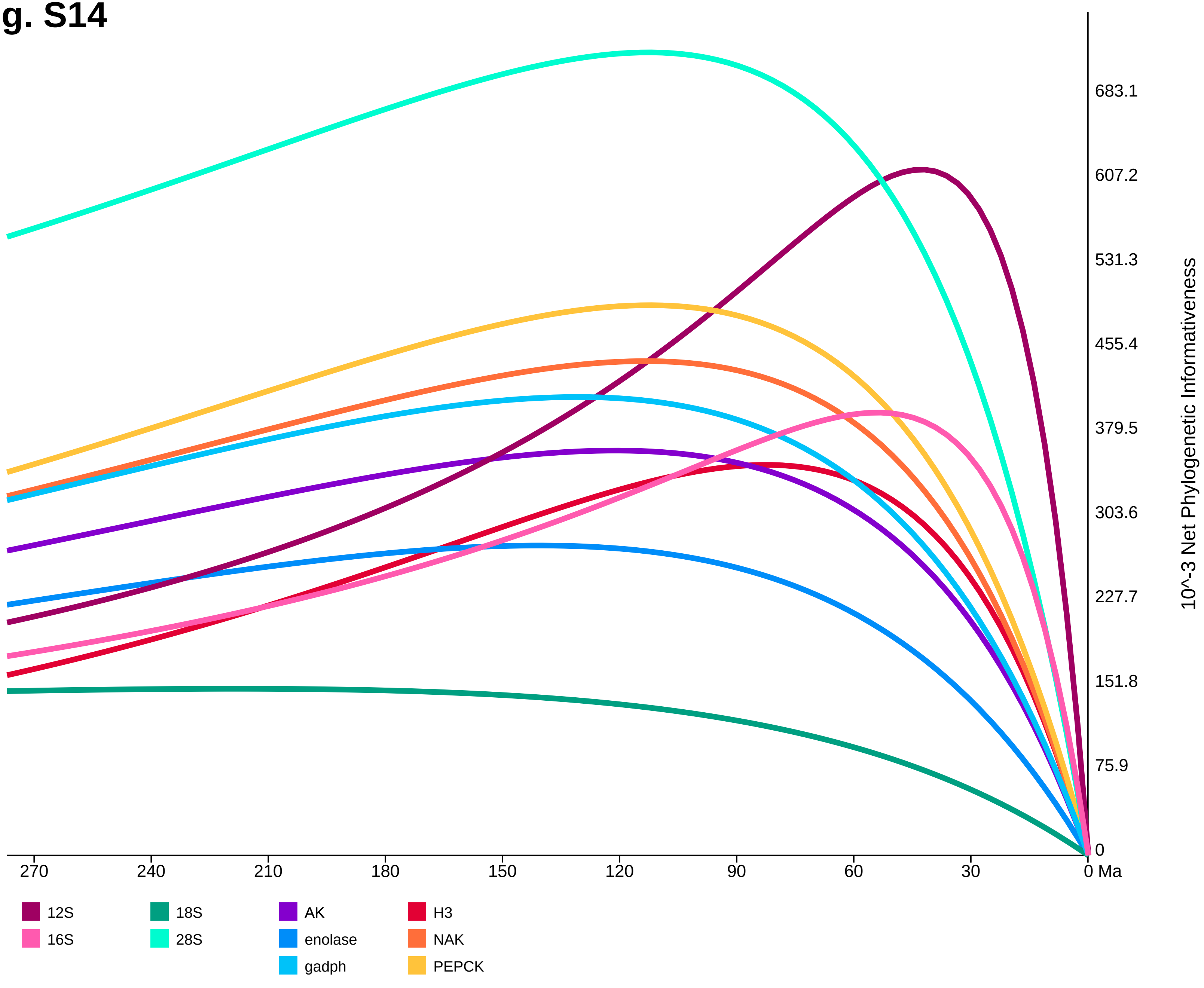
683.1
607.2
531.3
455.4
379.5
303.6
227.7
151.8
75.9
0
10^-3 Net Phylogenetic Informativeness
270
240
210
180
150
120
90
60
30
0 Ma
12S
16S
18S
28S
AK
enolase
gadph
H3
NAK
PEPCK

**Fig. S15**

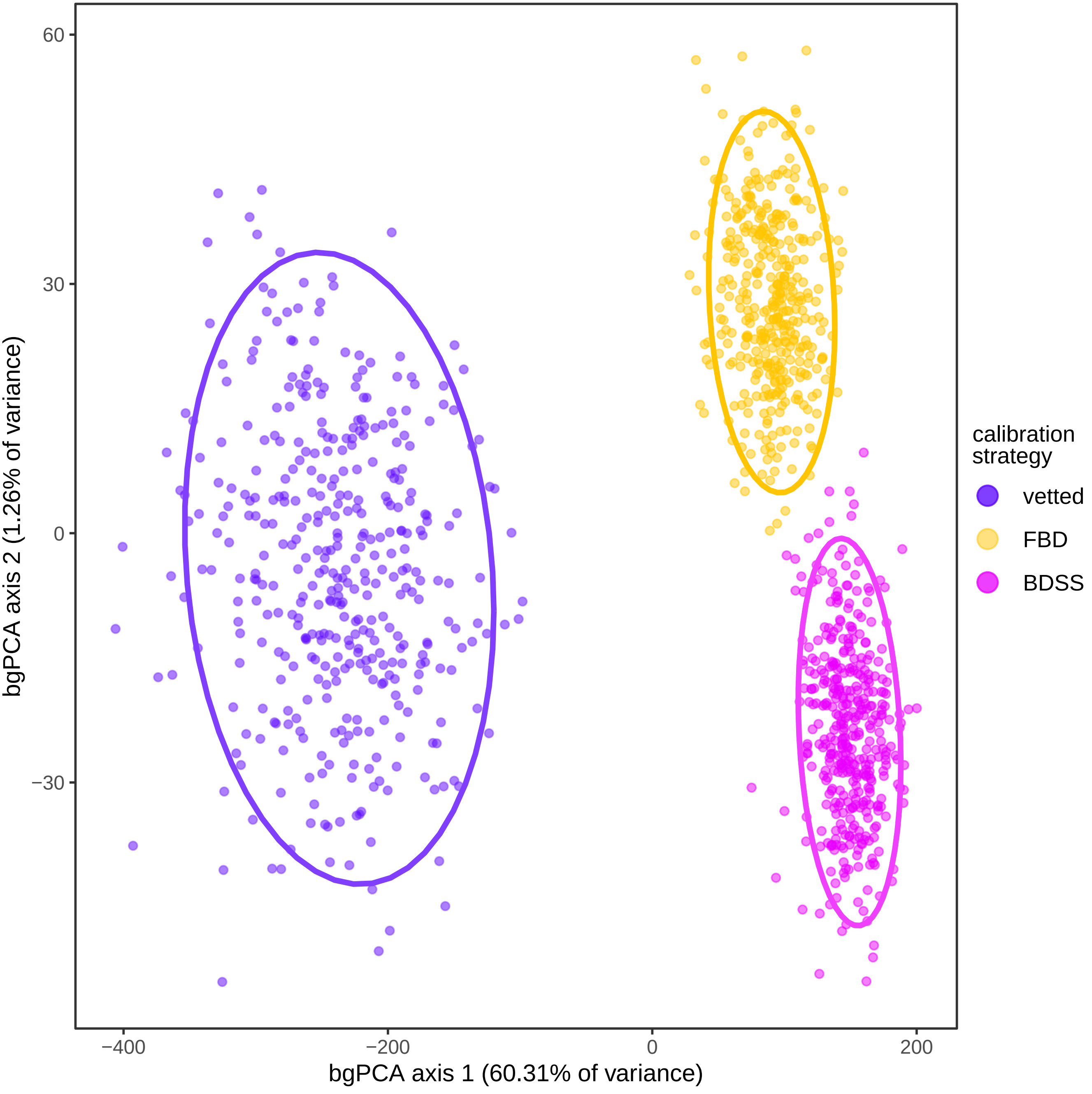
60
30
0
−30
−400
−200
0
200
bgPCA axis 1 (60.31% of variance)
bgPCA axis 2 (1.26% of variance)
calibration strategy
vetted
FBD
BDSS

Fig. S16

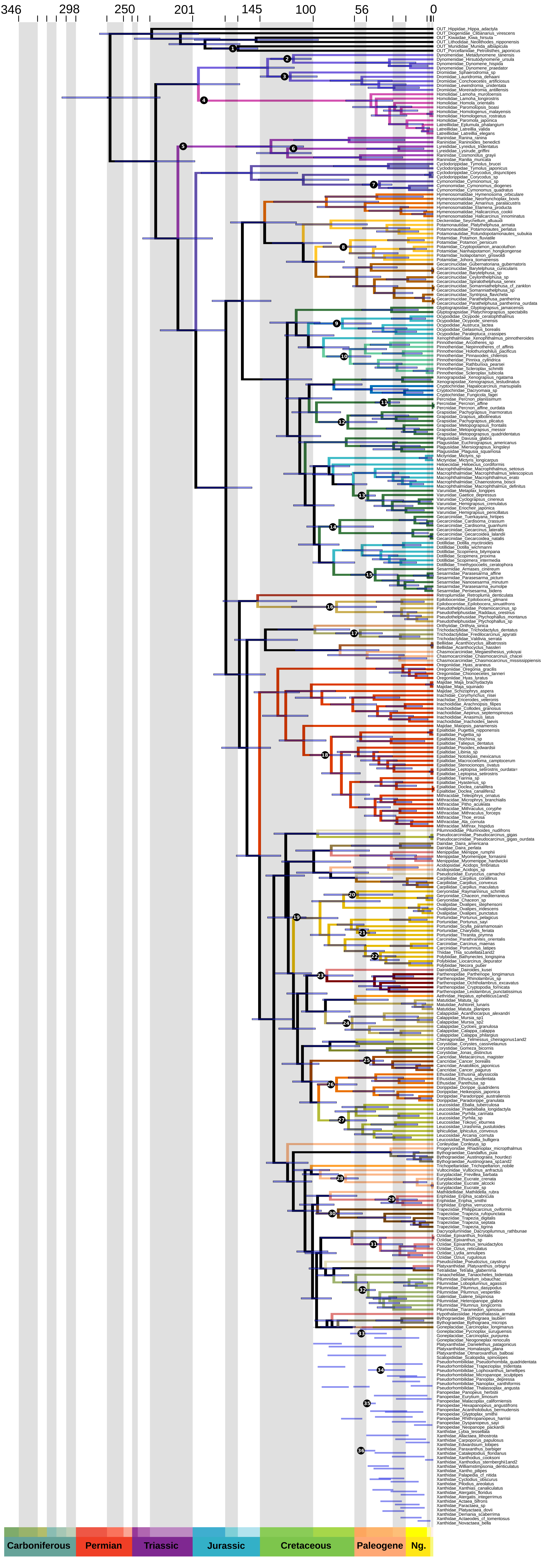
346
298
250
201
145
100
56
0
Carboniferous
Permian
Triassic
Jurassic
Cretaceous
Paleogene
Ng.

**Fig. S17**

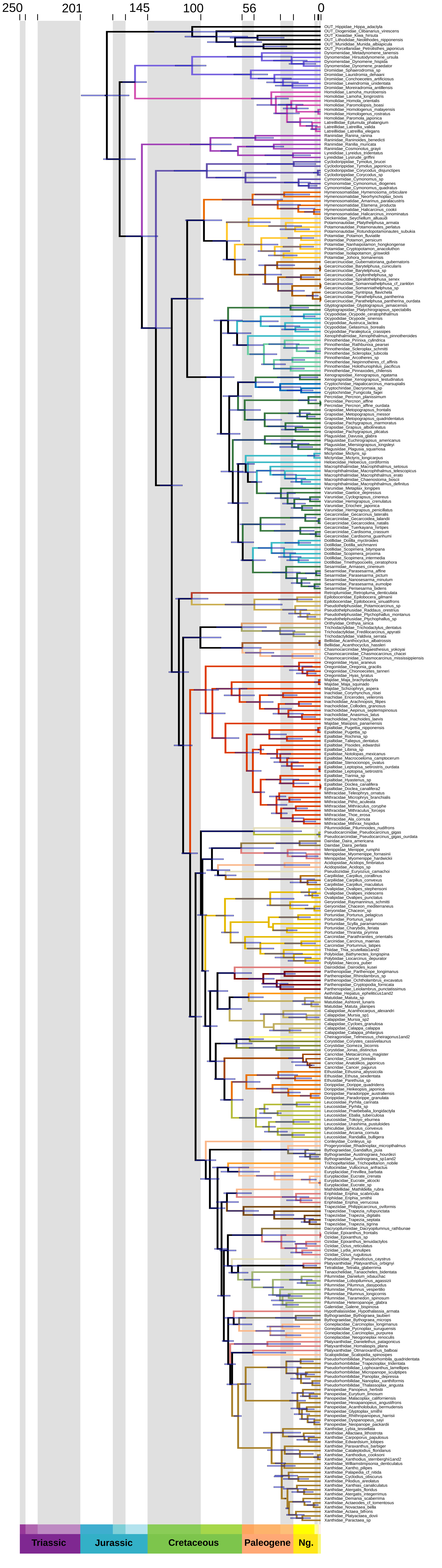
250
201
145
100
56
0
Triassic
Jurassic
Cretaceous
Paleogene
Ng.

**Fig. S18**

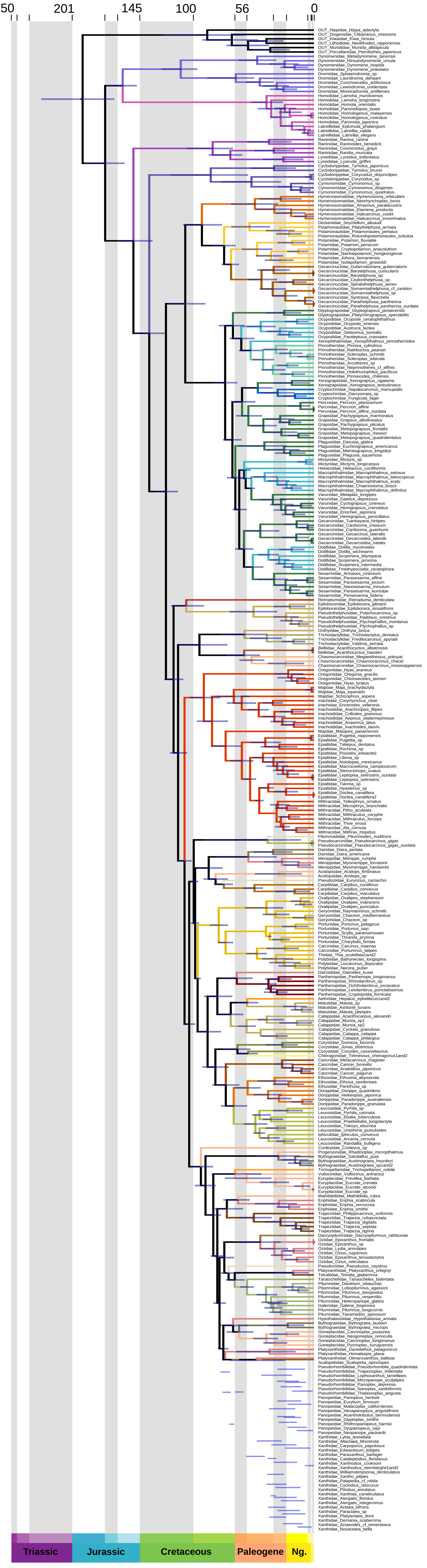
250
201
145
100
56
0
Triassic
Jurassic
Cretaceous
Paleogene
Ng.

### Fig. S19

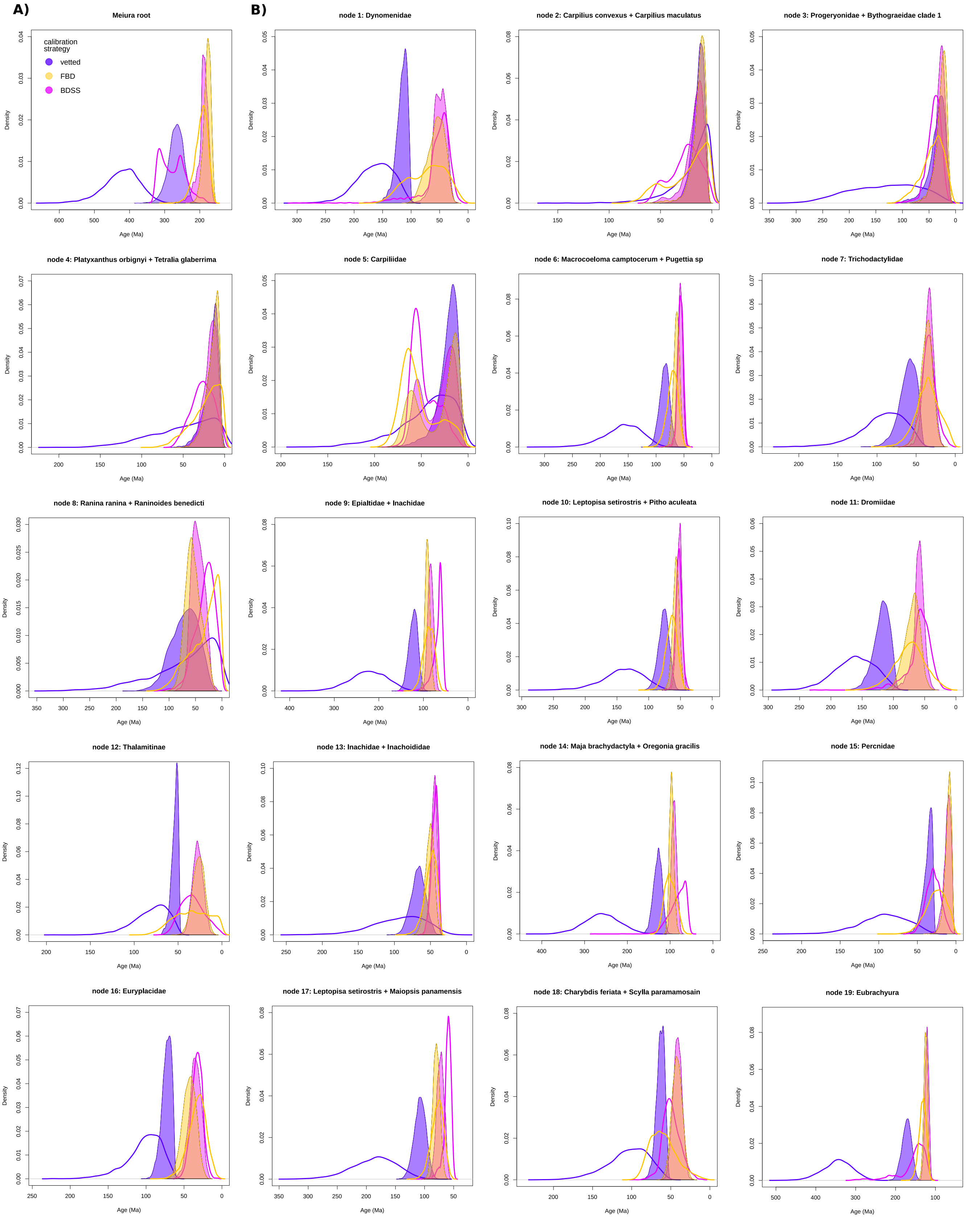

A)
B)
Meiura root
calibration strategy
vetted
FBD
BDSS
node 1: Dynomenidae
node 2: Carpilius convexus + Carpilius maculatus
node 3: Progeryonidae + Bythograeidae clade 1
node 4: Platyxanthus orbignyi + Tetralia glaberrima
node 5: Carpiliidae
node 6: Macrocoeloma camptocerum + Pugettia sp
node 7: Trichodactylidae
node 8: Ranina ranina + Raninoides benedicti
node 9: Epialtidae + Inachidae
node 10: Leptopisa setirostris + Pitho aculeata
node 11: Dromiidae
node 12: Thalamitinae
node 13: Inachidae + Inachoididae
node 14: Maja brachydactyla + Oregonia gracilis
node 15: Percnidae
node 16: Euryplacidae
node 17: Leptopisa setirostris + Maiopsis panamensis
node 18: Charybdis feriata + Scylla paramamosain
node 19: Eubrachyura
Density
Age (Ma)

### Fig. S20

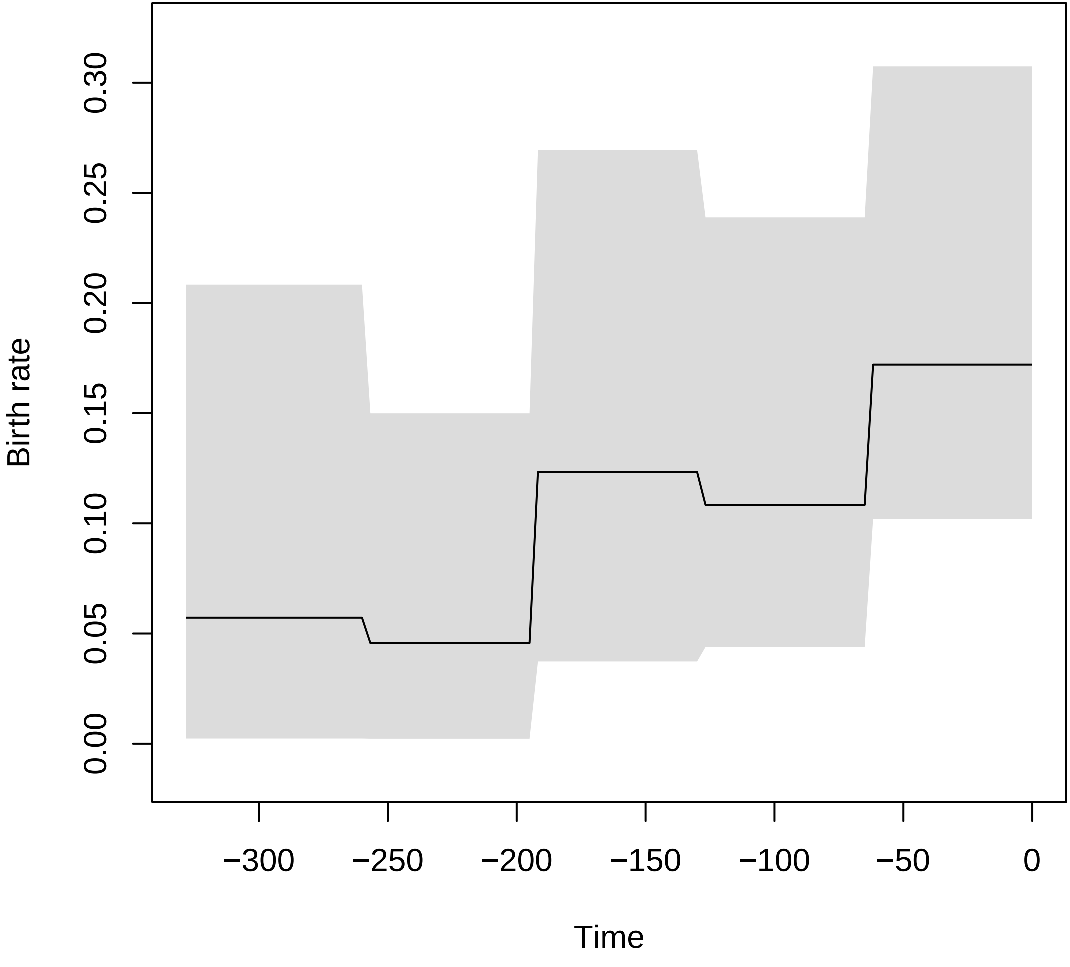

Birth rate
0.00
0.05
0.10
0.15
0.20
0.25
0.30
−300
−250
−200
−150
−100
−50
0
Time

**Fig. S21**

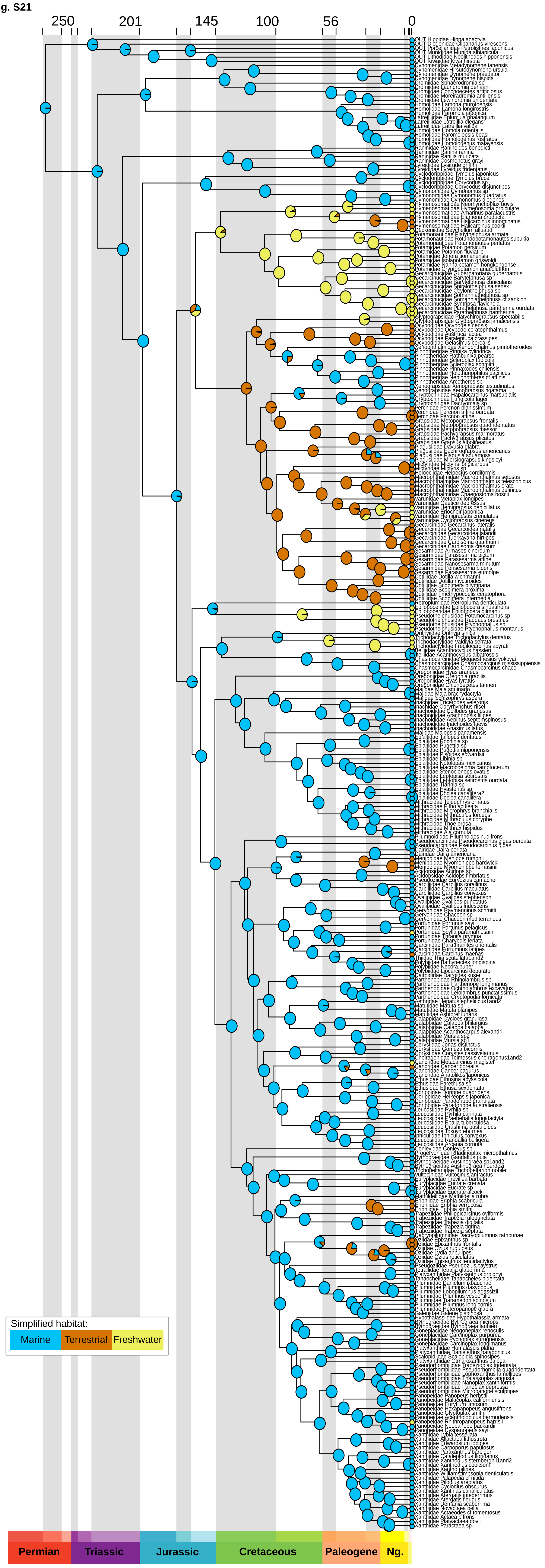
250
201
145
100
56
0
Simplified habitat:
Marine
Terrestrial
Freshwater
Permian
Triassic
Jurassic
Cretaceous
Paleogene
Ng.

**Fig. S22**

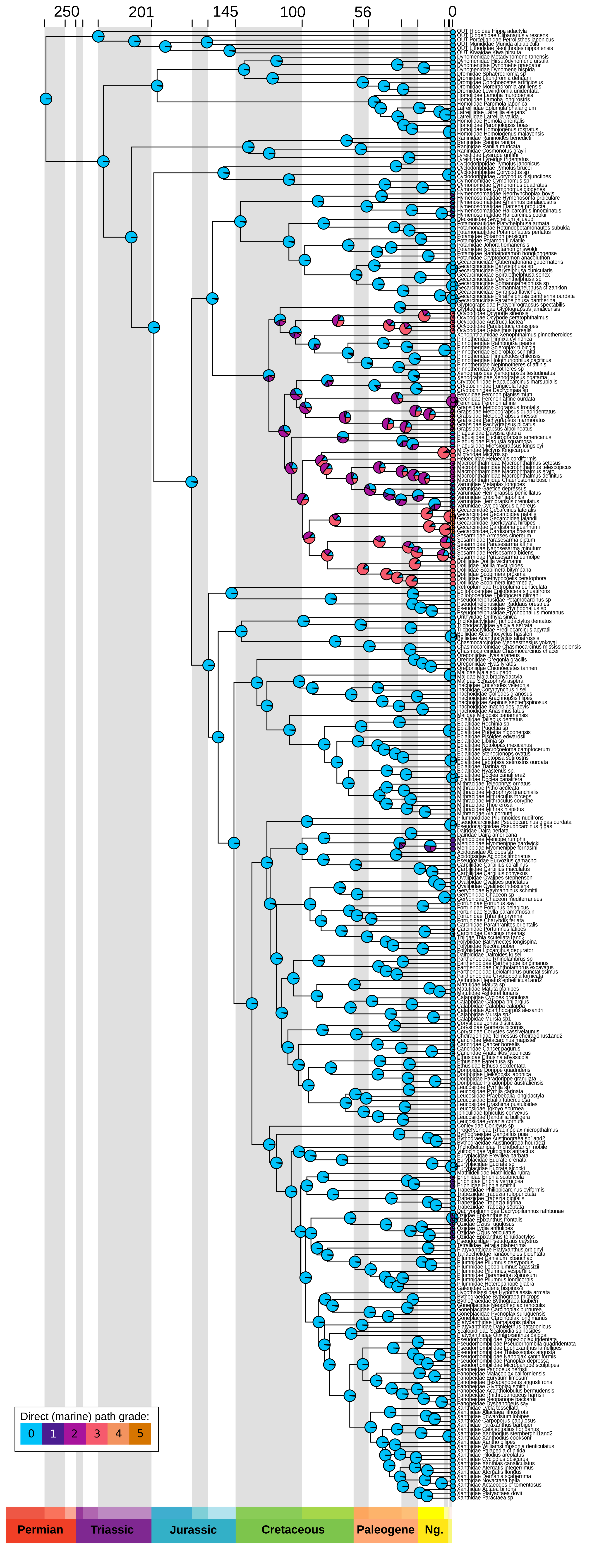
250
201
145
100
56
0
Direct (marine) path grade:
0
1
2
3
4
5
Permian
Triassic
Jurassic
Cretaceous
Paleogene
Ng.

Fig. S23

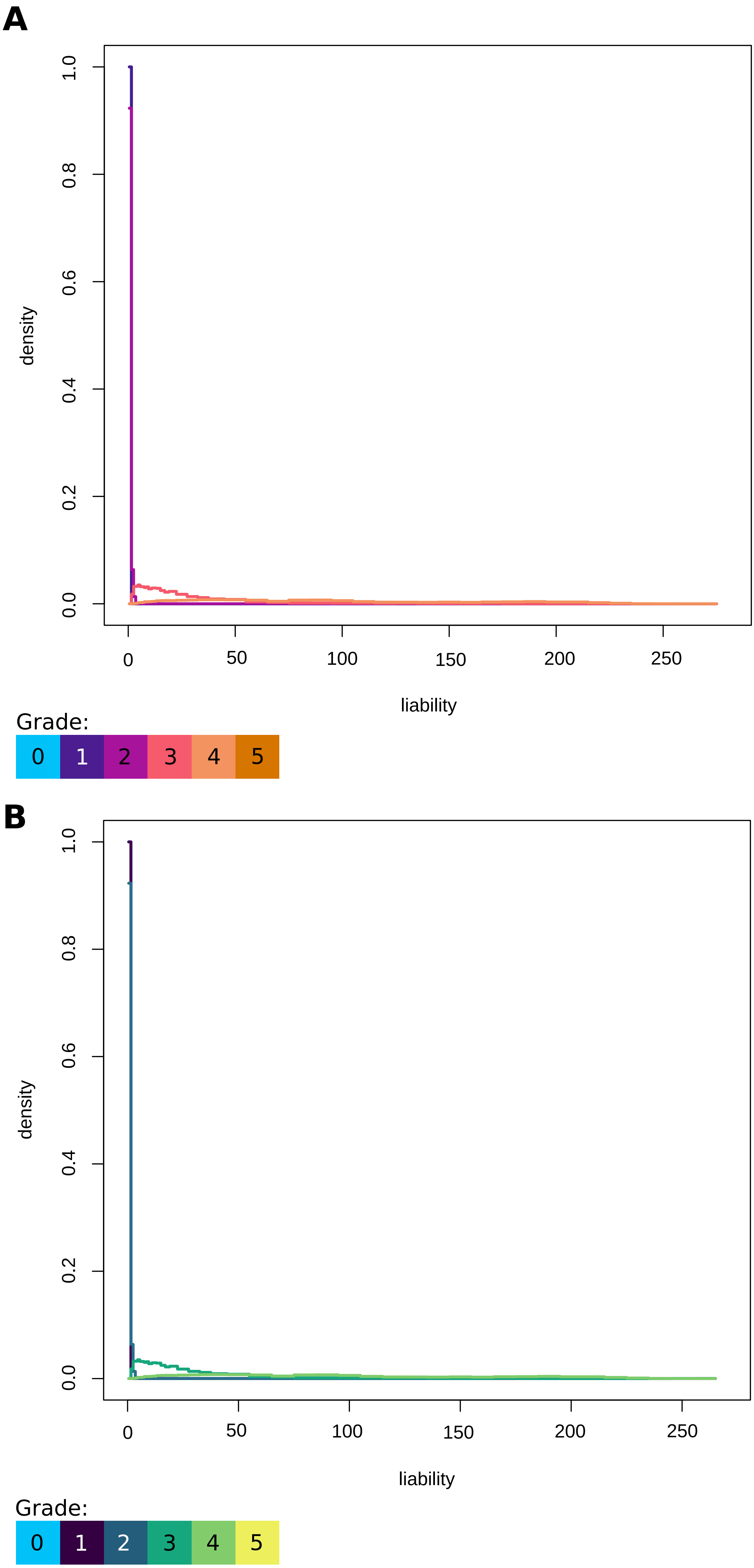
A
density
1.0
0.8
0.6
0.4
0.2
0.0
0
50
100
150
200
250
liability
Grade:
0
1
2
3
4
5
B
density
1.0
0.8
0.6
0.4
0.2
0.0
0
50
100
150
200
250
liability
Grade:
0
1
2
3
4
5

**Fig. S24**

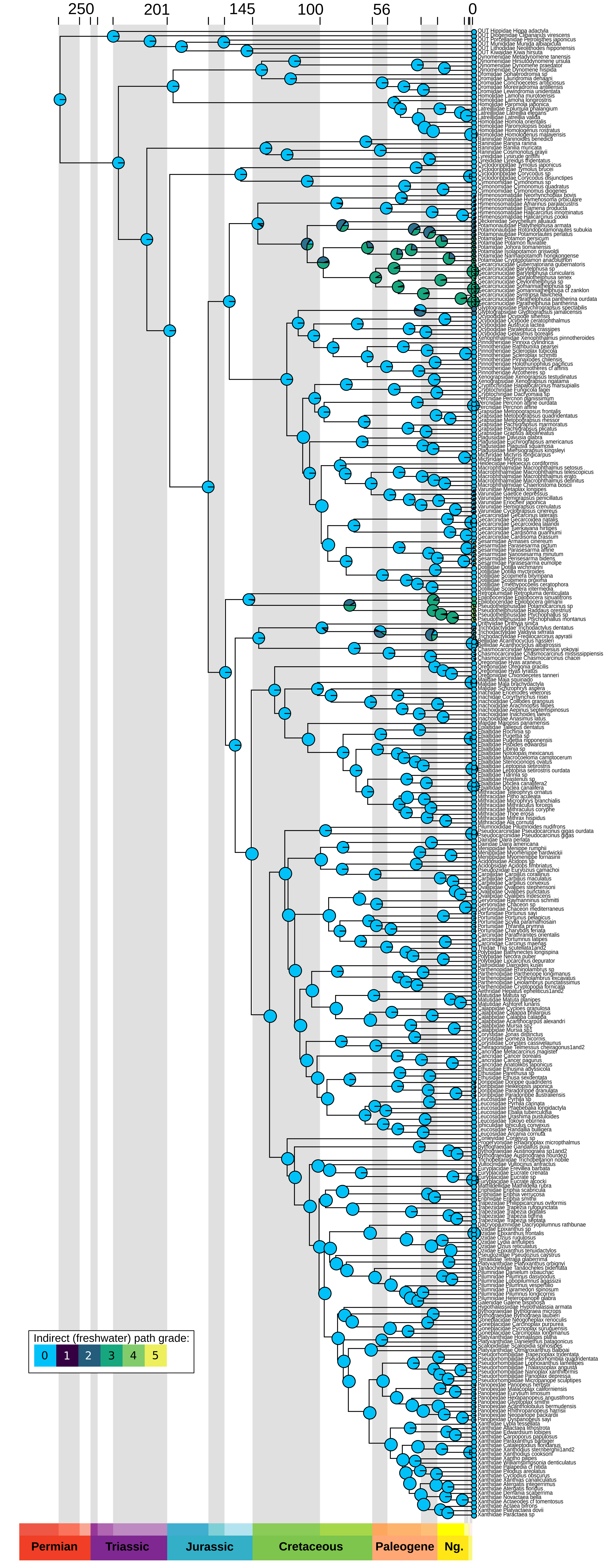
250
201
145
100
56
0
Indirect (freshwater) path grade:
0
1
2
3
4
5
Permian
Triassic
Jurassic
Cretaceous
Paleogene
Ng.

#### Fig. S25

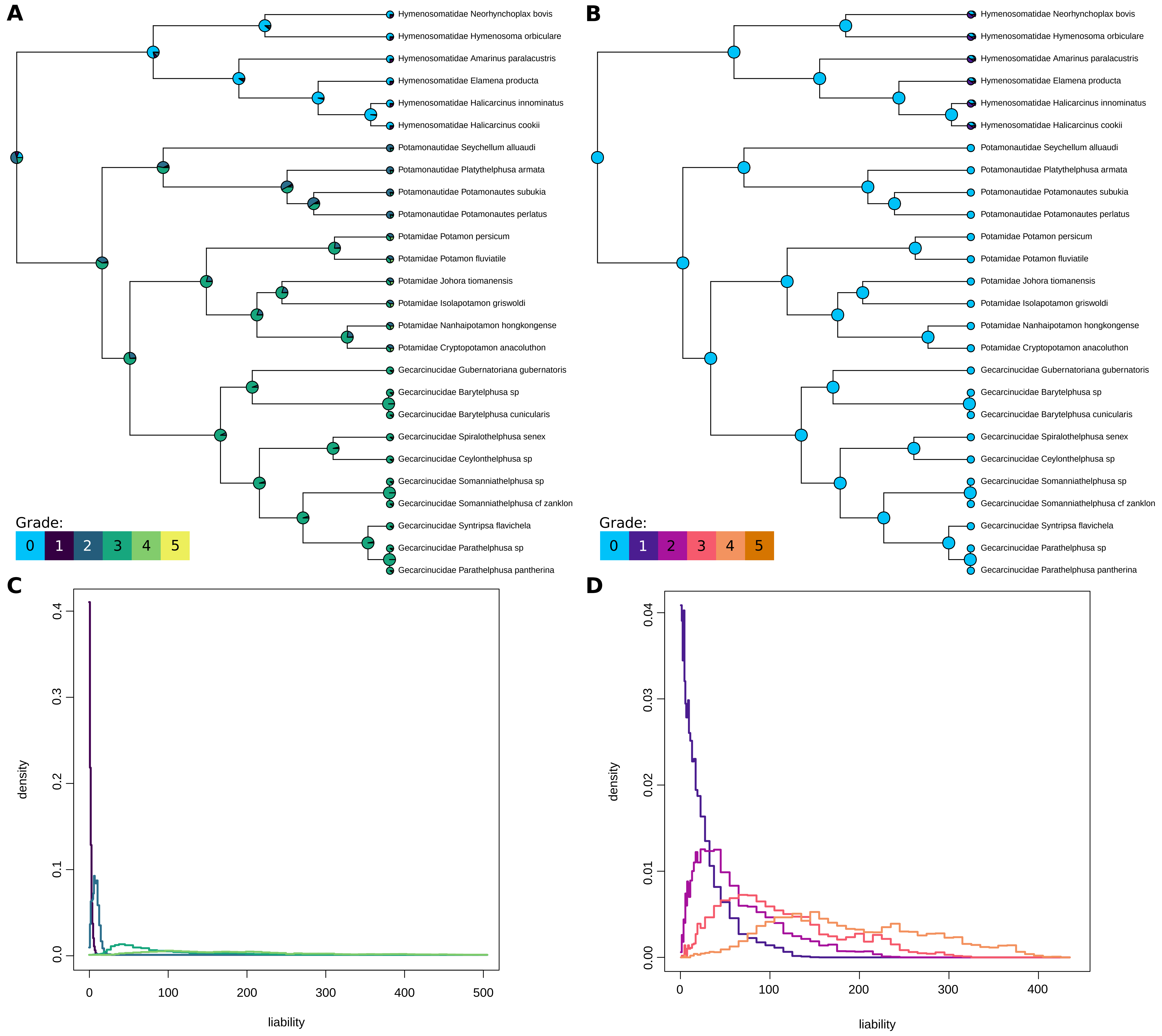
A
B
Hymenosomatidae Neorhynchoplax bovis
Hymenosomatidae Hymenosoma orbiculare
Hymenosomatidae Amarinus paralacustris
Hymenosomatidae Elamena producta
Hymenosomatidae Halicarcinus innominatus
Hymenosomatidae Halicarcinus cookii
Potamonautidae Seychellum alluaudi
Potamonautidae Platythelphusa armata
Potamonautidae Potamonautes subukia
Potamonautidae Potamonautes perlatus
Potamidae Potamon persicum
Potamidae Potamon fluviatile
Potamidae Johora tiomanensis
Potamidae Isolapotamon griswoldi
Potamidae Nanhaipotamon hongkongense
Potamidae Cryptopotamon anacoluthon
Gecarcinucidae Gubernatoriana gubernatoris
Gecarcinucidae Barytelphusa sp
Gecarcinucidae Barytelphusa cunicularis
Gecarcinucidae Spiralothelphusa senex
Gecarcinucidae Ceylonthelphusa sp
Gecarcinucidae Somanniathelphusa sp
Gecarcinucidae Somanniathelphusa cf zanklon
Gecarcinucidae Syntripsa flavichela
Gecarcinucidae Parathelphusa sp
Gecarcinucidae Parathelphusa pantherina
Grade:
0 1 2 3 4 5
C
D
density
liability
0.0 0.1 0.2 0.3 0.4
0 100 200 300 400 500
0.00 0.01 0.02 0.03 0.04
0 100 200 300 400

Fig. S26

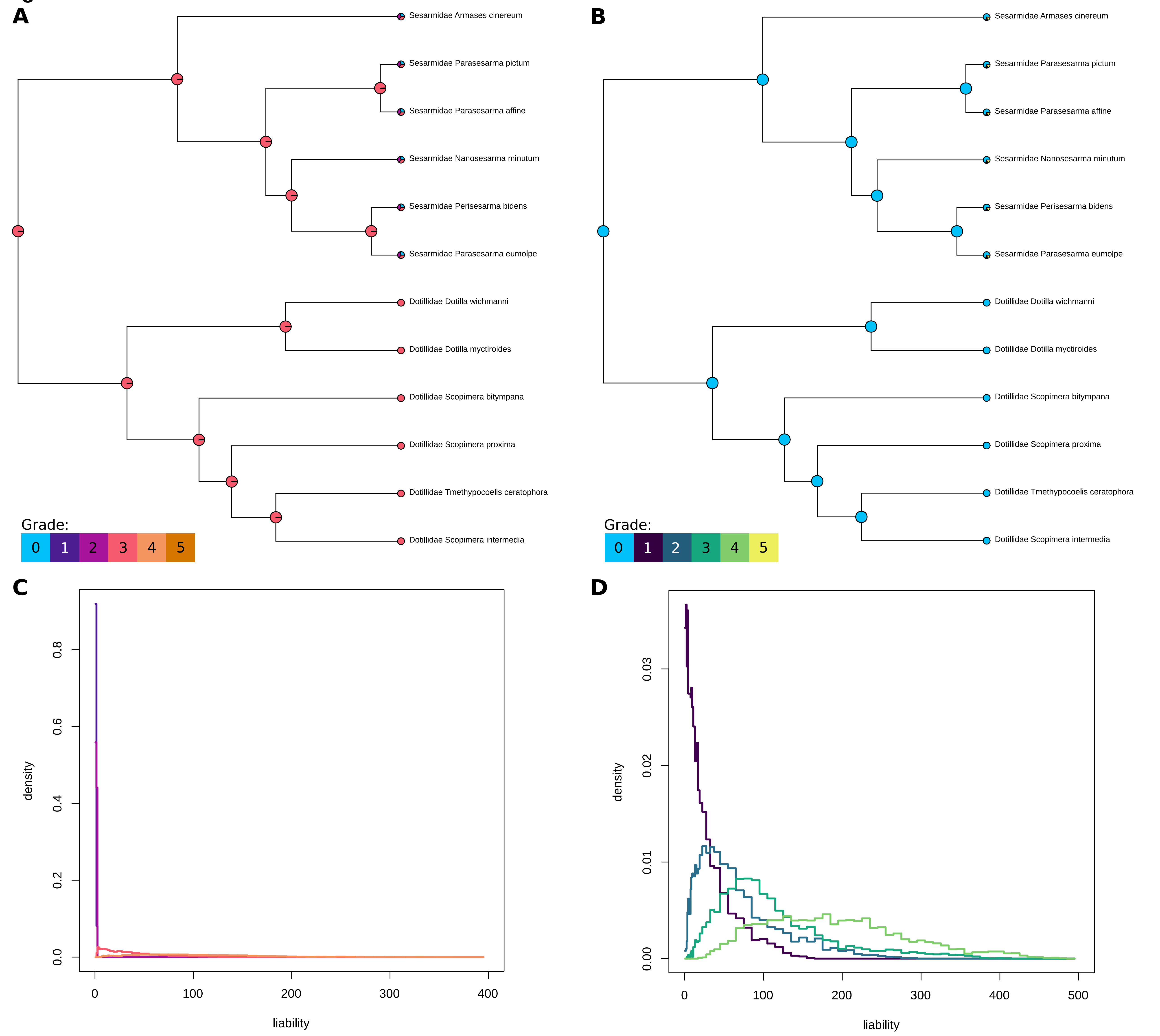

A
Sesarmidae Armases cinereum
Sesarmidae Parasesarma pictum
Sesarmidae Parasesarma affine
Sesarmidae Nanosesarma minutum
Sesarmidae Perisesarma bidens
Sesarmidae Parasesarma eumolpe
Dotillidae Dotilla wichmanni
Dotillidae Dotilla myctiroides
Dotillidae Scopimera bitympana
Dotillidae Scopimera proxima
Dotillidae Tmethypocoelis ceratophora
Dotillidae Scopimera intermedia
Grade:
0 1 2 3 4 5
B
Sesarmidae Armases cinereum
Sesarmidae Parasesarma pictum
Sesarmidae Parasesarma affine
Sesarmidae Nanosesarma minutum
Sesarmidae Perisesarma bidens
Sesarmidae Parasesarma eumolpe
Dotillidae Dotilla wichmanni
Dotillidae Dotilla myctiroides
Dotillidae Scopimera bitympana
Dotillidae Scopimera proxima
Dotillidae Tmethypocoelis ceratophora
Dotillidae Scopimera intermedia
Grade:
0 1 2 3 4 5
C
density
0.0
0.2
0.4
0.6
0.8
0
100
200
300
400
liability
D
density
0.00
0.01
0.02
0.03
0
100
200
300
400
500
liability
